# Comprehensive analysis of RNA-chromatin, RNA- and DNA-protein interactions

**DOI:** 10.1101/2024.03.13.584417

**Authors:** Daniil A. Khlebnikov, Arina A. Nikolskaya, Anastasia A. Zharikova, Andrey A. Mironov

## Abstract

RNA-chromatin interactome data is considered to be one of the noisiest types of data in biology. This is due to protein-coding RNA contacts and non-specific interactions between RNA and chromatin caused by protocol specifics. Therefore, finding regulatory interactions between certain transcripts and genome loci requires a wide range of filtering techniques to obtain significant results. Using data on pairwise interactions between these molecules, we propose a concept of triad interaction involving RNA, protein and a DNA locus. The constructed triads show significantly less noise contacts and are more significant when compared to a background model for generating pairwise interactions. RNA-chromatin contacts data can be used to validate the proposed triad object as positive (Red-ChIP experiment) or negative (RADICL-Seq NPM) controls. Our approach also filters RNA-chromatin contacts in chromatin regions associated with protein functions based on ChromHMM annotation.

## Introduction

It is well recognised that eukaryotic RNAs play multiple roles beyond their direct role in transporting transcribed genetic information in the nucleus: major advances include understanding the functions of non-coding RNAs such as XIST, NEAT1, MALAT1, HOTAIR, FIRRE, etc. [1], [2]. The study of new potential functions of chromatin-associated RNAs is of interest to molecular biology in the context of searching for DNA loci and genes whose interaction with RNA leads to gene expression regulation [3].

When studying the pool of RNA-chromatin contacts, there are two main approaches to designing the interaction detection experiment. ‘One-to-all’ experiments, such as RAP [4], CHART-seq [5], and ChIRP-seq [6], focus on single RNA contacts across the genome by pulling down RNA-chromatin complexes of a particular RNA. These experiments are considered to be the gold standard for a further class of methods of the ‘all-to-all’ type. These methods address the challenge of identifying pairwise contacts between all RNAs and every genomic locus. The key step in the experimental protocol is to fix interacting RNAs and DNA loci by bridge ligation of interacting nucleic acids. This fixation predominantly occurs through proteins. Methods to search for RNA-chromatin interactions include iMARGI [7], [8], RADICL-Seq [9], GRID-Seq [10], [11], [12] and Red-C [13]. This study analyses data from GRID-Seq, RADICL-Seq and Red-C.

The all-to-all experiments show that most of the RNA-chromatin contacts detected are those of protein-coding RNAs [14]. The presence of regulatory functions for these RNAs has not been well established and this observation reflects a high level of non-specific interactions. For most RNAs, the density of contacts near the gene of that RNA peaks and then declines in a power law fashion [13], [14], a phenomenon we call RD-scaling. This is caused by the contacts of incompletely transcribed nascent RNA, as well as by RNA that diffuses away from the site of transcription. Importantly, RD-scaling is a feature of both all-to-all and one-to-all data. However, the RNA-chromatin interaction data reveals several other types of biases. First, contacts may be non-specific, as RNA can contact chromatin proteins such as histones, albeit with lower affinity. The probability of observing contact depends on chromatin accessibility, as heterochromatin regions are less accessible to nucleases and less susceptible to degradation by sonication. Normalisation to background is applied to compensate for this bias. Various peak calling methods are used to reduce the influence of these biases and partially suppress the noise. The most commonly used approaches include MACS2 peak calling [15] and custom dataset-specific peak calling algorithms (GRID-Seq [10], [11], [12]). These approaches do not take into account the bias introduced by RD-scaling. A recently developed BaRDIC peak-calling algorithm is designed to account for these biases [16]. The RNA-protein interaction data can also contain a significant amount of non-specific interactions [17], [18], and appropriate peak calling algorithms are also used to suppress the noise. Red-ChIP [19] detection of RNA-chromatin contacts enriched for EZH2 and CTCF proteins by immunoprecipitation was developed to isolate contacts mediated by these proteins. A ChRD-PET [20] protocol that includes H3K4me3 immunoprecipitation and ChIP-Seq filtering of contacts has been proposed for the study and analysis of RNA-chromatin interactions in *Oryza sativa*.

We compared three types of data: RNA-chromatin contacts (RD-contacts), protein-chromatin contacts (PD-contacts), and protein-RNA contacts (PR-contacts) to understand which proteins mediate the contacts of which RNAs with chromatin. The data used in this study includes RNA-chromatin interactions from RADICL-Seq [9], GRID-Seq [10], [11], [12], and Red-C [13], DNA-protein interactions from ChIP-seq [21], and RNA-protein interactions from RIP-Seq [22], fRIP-Seq [23], and eCLIP [24], [25]. The intersection of these data is referred to as interaction triads, as shown in **Figure 1A**. Pairwise contacts within the triads are denoted as RDt-contacts, PDt-contacts, and PRt-contacts. The proposed concept of interaction triads involves a two-stage filtering process of RNA-DNA interaction data using available RNA-protein and DNA-protein interactomes.

**Fig. 1.**
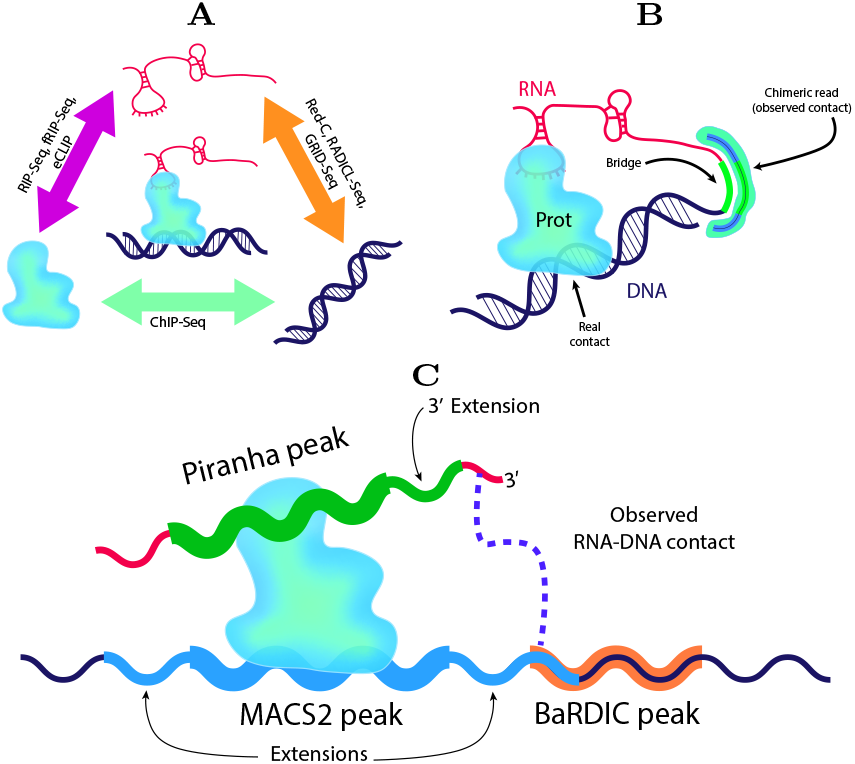
**(A)** The concept of constructing RNA-protein-DNA triads using pairwise interaction data. **(B)** The observed contacts in the all-to-all data may be located away from the actual contacts. This creates inaccuracy in the data. **(C)** To obtain more accurate data on protein-mediated RNA-DNA interactions, peak expansion is required.

This approach enabled us to decrease the aforementioned noise level. We assessed the statistical significance of the triads obtained. When estimating the biological significance of the triads, we observed an association between the triads and the ChromHMM and SPIN putative chromatin states. Additionally, we analysed the evolutionary stability of the identified triads by comparing human and mouse data. We demonstrate that “triads” of RNA-protein-DNA interactions are an excellent validation target for testing the consistency of different methods in identifying biologically relevant contacts and filtering out data noise.

## Materials & Methods

### Initial data

To construct triads of RNA-protein-DNA interactions mediated by one of the 47 nuclear proteins, we used various types of data. These included processed ChIP-Seq experiment data that defined DNA loci interacting with the target protein, raw data from the RIP-, fRIP-, and eCLIP-Seq experiments that defined sites on RNA that interact with the target protein, and our previously processed data for RNA-chromatin interactomes of the Red-C, GRID-Seq, and RADICL-Seq experiments for human (K562) and mouse (mESC) cell lines from RNA-Chrom database [14]. All IDs used from the databases can be found in **Suppl. Table S1**.

ChIP-Seq data were obtained from the public biological databases ENCODE, ReMap [26] and GTRD [27] as DNA peaks derived from the MACS2 peak caller with a q-value threshold of 0.05. Standardisation of data processing and peak calling protocols was verified.

### RNA-Protein interactions data processing

We processed data from RNA-protein immunoprecipitation experiments to determine RNA interactions for target proteins in a unified pipeline. Raw RNA reads from experimental and control replicates were mapped to the human genome version GRCh38.p13 using the hisat2 aligner (v2.2.1) [28] and then binned across the human genome (bin size 300 bp) for more accurate peak searching. The reads were binned and then analysed using the Piranha [29] RNA-protein peak-calling software. The input data was specified as the background distribution for normalization, and a filtering q-value threshold of 0.05 was applied. The resulting peaks were annotated using the gene annotation from the RNA-Chrom database. Mapped reads were annotated with the same annotation. For each PR-interaction, the fold change of contact was calculated as the ratio of the fraction of gene counts in the experimental replicate to the fraction of gene counts in the control replicate, with an additional pseudo-count of 0.01. Genomic intervals were manipulated using the bedtools package (v2.27.1) [30].

### Triads construction

BaRDIC [16] peak calling software was used to process data from RNA-chromatin interactions (BaRDIC parameters are listed in **Suppl. Table S2**). Only contacts falling within BaRDIC peaks were selected, as we are interested in individual contacts. It should be noted that experimental RD-contacts detection assumes spatial proximity ligation, meaning that the observed contact may not be the same as the actual contact (**Figure 1B**). To construct triads, the protein ChIP-seq peaks were extended by 2 Kb on either side of the peak, and the Piranha peaks were extended by 100 bp on the 3’ end (**Figure 1C, Suppl. Table S3**). The RD-contacts were then intersected with PD-contacts and PR-contacts (RIP-Seq / fRIP-Seq / eCLIP) for each protein separately. Additionally, we discarded the RD-contacts if the length of intersection with PD-or PR-peaks was less than 19 bases. Filtering was necessary to reduce the number of false-positive triads resulting from the intersection of close contacts with a peak. Due to the inaccuracies and noise associated with RNA-chromatin contact data, this filtering may discard some true-positive triads along with the false-positive ones. However, we assume that for datasets with extended peaks, the proportion of true-positive triads discarded is minimal. The triads were visualized using the circos-v0.69.9 suite [31]. Red-ChIP data were processed according to the protocol of the study presenting the method [19].

### Triad annotation using ChromHMM and SPIN

The triads obtained in the previous step were annotated with grouped annotations of ChromHMM chromatin states and SPIN chromatin compartments. The annotations were taken from the corresponding articles of their origin [32], [33] and [34]. To simplify the analysis, we combined annotation states into groups (refer to **Suppl. Table S4** for ChromHMM, **Suppl. Table S5** for SPIN).

### RNA-Chromatin contacts orthologs search

The ortho2align [35] software was used to select bidirectional best hits from the coordinates of RNAs involved in triads formed by each protein in human and mouse cell lines. Next, the DNA contacts of each RNA were extracted in both cell lines, and the nearest DNA loci [36] among the contacts were searched for orthologous RNA pairs from different organisms. Finally, bedtools was used to annotate the nearest genes. The correlation of genomic intervals was carried out using StereoGene [37]. The parameters for the runs can be found in **Suppl. Table S6**.

Data visualizations and analyses not previously mentioned were performed using in-house scripts in Python, bash, and R. Data simulations were conducted to verify the randomness of the resulting data using Python scripts. All scripts are available in the corresponding GitHub repository (github.com/dkhlebn/shift_IP_peaks).

## Results

We have selected 47 eukaryotic nuclear proteins that have available RNA and DNA immunoprecipitation data. These proteins are involved in chromatin remodeling and modification, regulation of transcription, and regulation of RNA processing and splicing. The selected proteins include PRC2 proteins (EZH2, SUZ12), several ribonucleoproteins (hnRNPC, H, K, L, U, UL1), various splicing factors (SRSF, U2AF family proteins), and chromatin remodelers (HDAC1, DNMT1, PCAF, LSD1). The proteins were divided into functional groups (**Suppl. Table S7**) to simplify the analysis and draw meaningful conclusions for protein groups rather than individual proteins.

### Constructed triads demonstrate less noise than original data

**Figure 2A-C** displays examples of RD-triad graphic visualizations that we constructed. It is evident that trans-RDt-contacts make up the majority of triad contacts. **Suppl. Table S8** displays the distribution of cis- and trans-contacts numbers for protein triads. It is clear that, with the exception of the triads mediated by the FUS protein, most proteins exhibit predominantly trans-contacts in the filtered contacts.

**Fig. 2.**
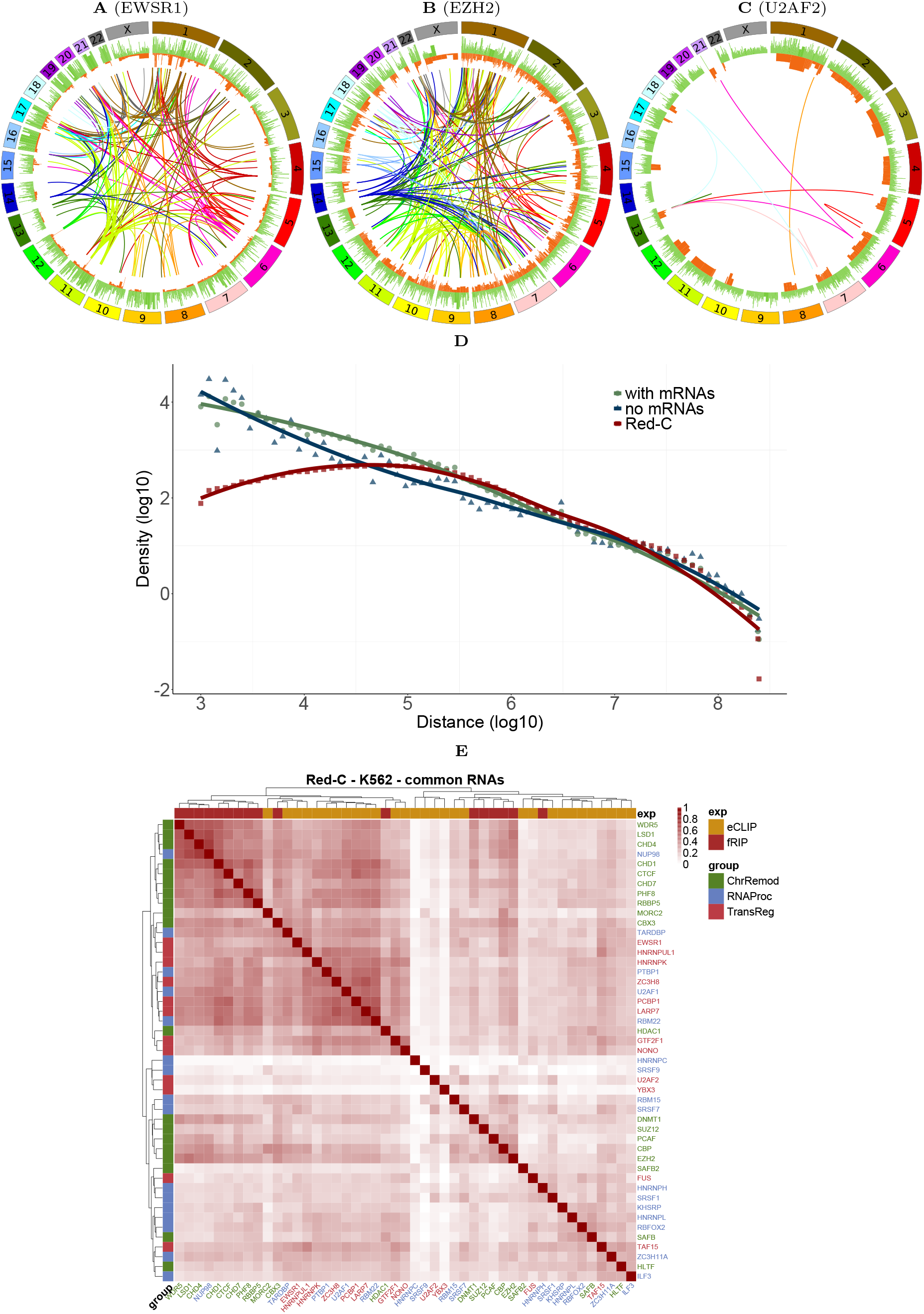
RNA-DNA interaction triads for three proteins: **(A-C)** Schematic diagrams show the interaction triads of EWSR1 **(A)**, EZH2 **(B)**, and U2AF2 **(C)** proteins. (see **Suppl. Fig. S1** for the remaining proteins). The outer histogram (green) represents PR-peaks, while the inner histogram track (red) represents PD-peaks. The arc connecting the locations of the RNA and DNA involved in the triad is coloured according to the chromosome on which the RNA-part is located. Only triads that are not mediated by protein-coding RNAs are shown. **(D)** RDt scaling in cis-triads based on data from the Red-C experiment. **(E)** Jaccard’s score was used to measure the similarity of triad datasets based on shared RNAs between each pair of datasets.

Any existing cis-contacts have RDt-scaling similar to that of the original RNA-chromatin interaction data **Figure 2D**. A paired Wilcoxon test was performed for each RD-experiment (**Table 1**). The statistics for select RNAs are shown in the **Suppl. Table S9**. The ratio of cis-to-trans contacts decreases when comparing triads to RD-experiment data for all protocols except GRID-Seq, specifically for Red-C and RADICL-Seq.

**Table 1.**
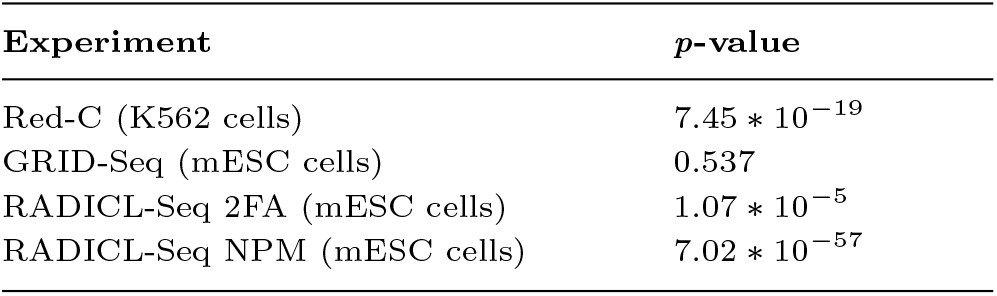
Test results for changes in the ratio of cis-to trans-contacts.

This suggests that the interaction triad data we obtained are qualitatively similar and may be worth further investigation.

The intersections of the sets of RNAs forming triads with each of the proteins were analyzed to identify common contacts (**Figure 2E**). Proteins with similar functions were found to be clustered together, which can be attributed to their participation in a common process and a larger pool of common RNAs forming triads with them.

**Table 2** presents the number of triads found in total and by RNA and protein types. It is known [38], [13], [16] that contacts of protein-coding RNAs can account for up to 80% of all contacts found in RNA-chromatin interactome experimental data. However, the share of mRNA contacts decreases significantly when filtering the data to construct triads (**Table 2, Suppl. Table S8**). It is evident that the proportion of non-coding RNAs has increased significantly. However, the total number of contacts mediated by any one of the 47 proteins accounts for no more than 10% of the original dataset of RNA-chromatin interaction contacts. The unavailability of data for all RNA-chromatin associated proteins, incomplete and noisy RNA-chromatin interactome data and other pairwise protein to nucleic acids interaction data explain this. Our proposed method of triad construction can also filter out noisy PD- and PR-contacts in other pairwise data, as demonstrated in **Suppl. Table S10, S11, S12, S13, S14**.

**Table 2.**
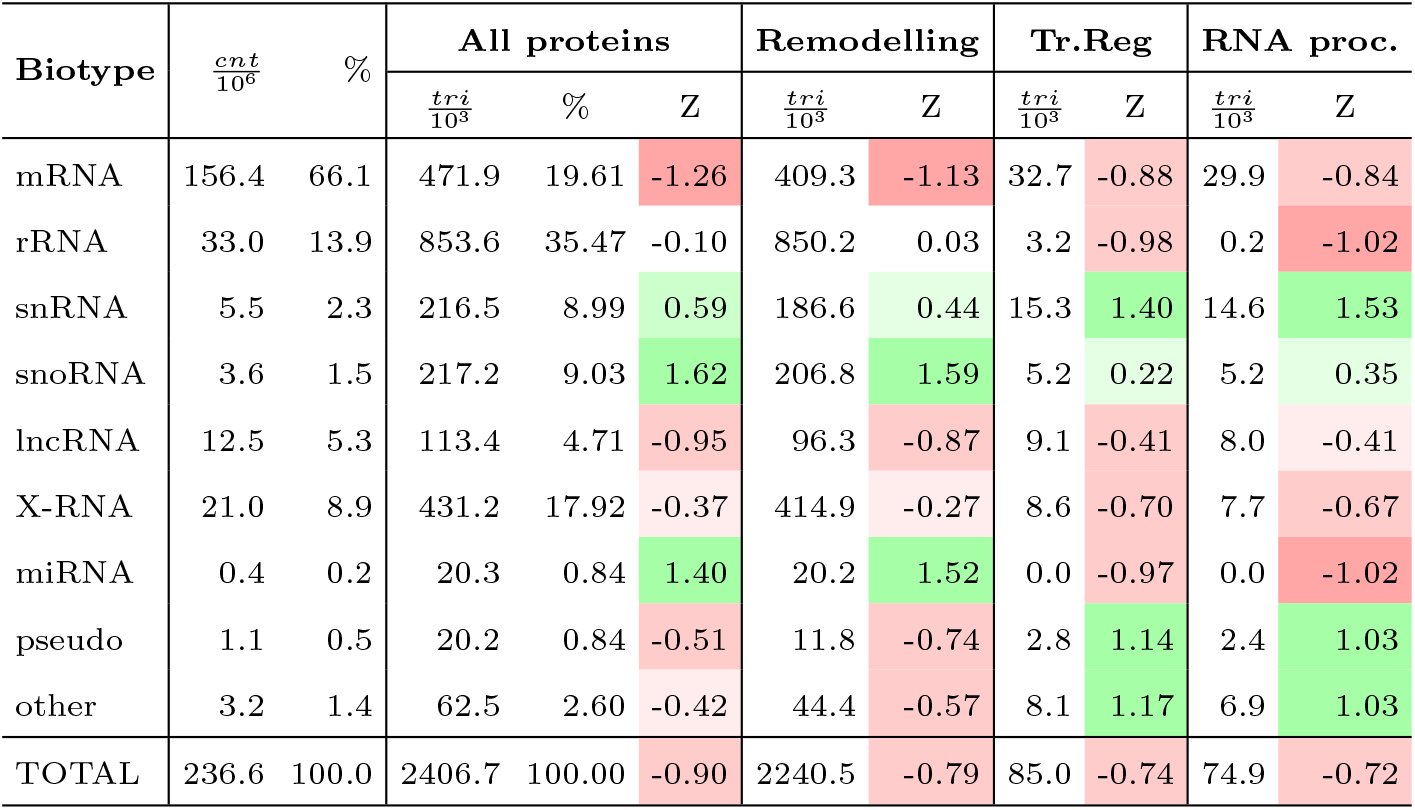
Triads composition by RNA biotypes for all proteins split by function. pseudo – pseudogenes; *cnt* – number of observed contacts; *tri* – number of triades; *Z* – z-score of 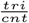 distribution over the RNA biotypes.

We attempted to compare triads assembled on one-to-all and all-to-all RNA-DNA interaction data of Malat1 and Hdac2 RNA for mESC cells using data from RAP and ChIRP-Seq experiments. However, we were again unable to draw any interpretable conclusions due to the incompleteness and noisiness of the pairwise interaction data.

The descriptive statistics for the content of the constructed triads suggest that the results obtained provide a clearer sample of RDt contacts mediated by the proteins studied. This sample can be characterised by a much lower level of noisy cis contacts and RD contacts formed by mRNAs.

### Constructed triads are statistically significant

To evaluate the statistical significance of the obtained triads, we created a background model for PR- and PD-contact data that maintains the biological structure of the original data. To achieve this, we shuffled the PD-peaks and PR-interactions of the studied proteins and constructed triads using these datasets. As our focus was on studying RNA-DNA interactions, we did not alter the RD-contacts data during the simulation.

When simulating ChIP-Seq peaks of proteins, we aimed to preserve the intrinsic structure of PD-contacts, which consisted of two components. Firstly, we had to preserve the profile of interactions, i.e. the height of ChIP-Seq peaks. Secondly, we had to consider and preserve each peak annotation by nuclear A/B-compartment. We retained the annotation of the peak by A- and B-compartments as it contains crucial biological sinformation about protein-DNA interactions. To obtain the shuffled ChIP-Seq tracks, each protein peak was randomly shifted up-or downstream by a number of megabases (1, 3, 5, 7, or 10) predetermined independently for each chromosome. If the ChIP-Seq peak in the original data was in the A-compartment, its shifted version in the generated data would also be in the A-compartment (**Figure 3A**, upper panel).

**Fig. 3.**
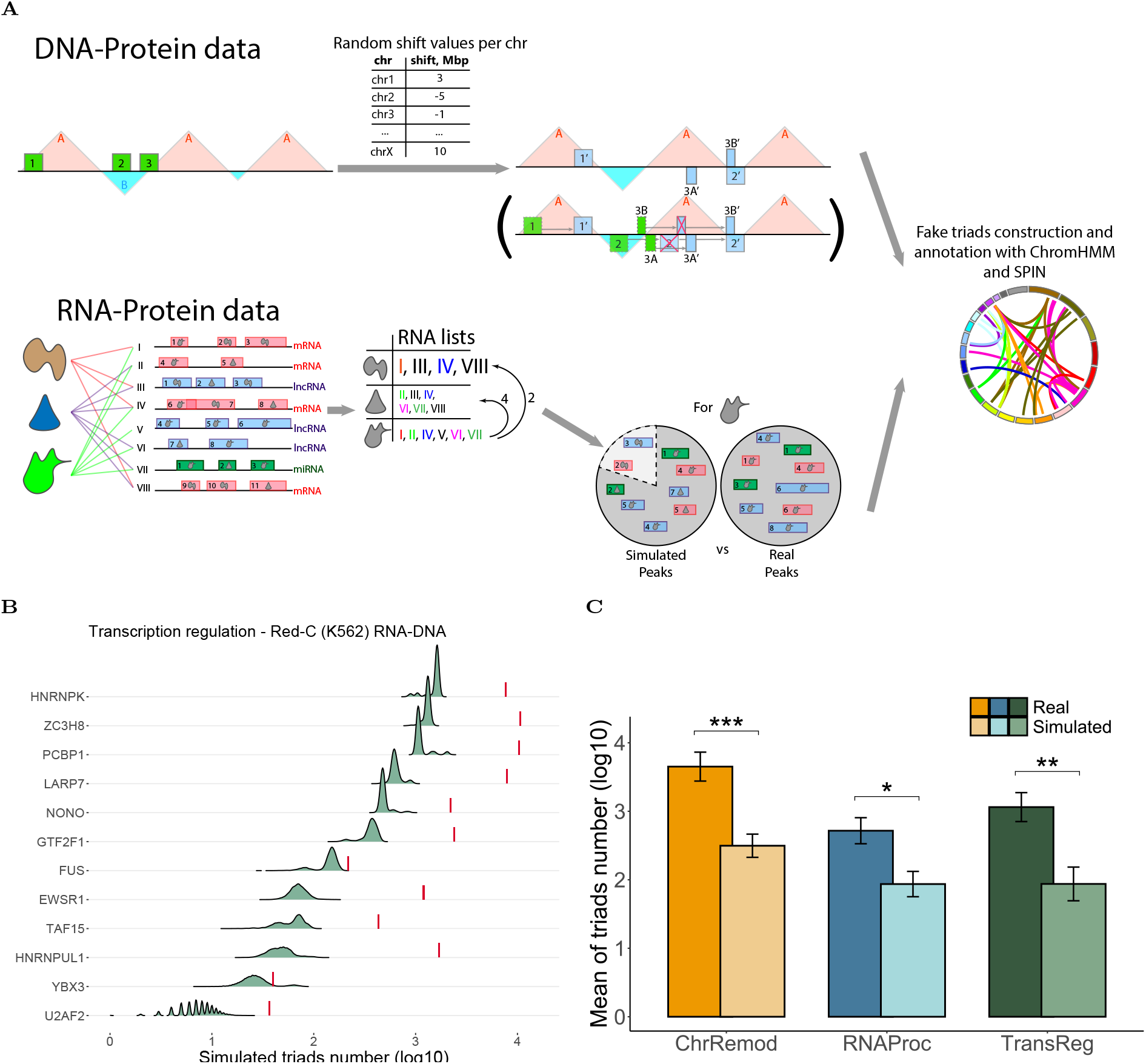
Simulation scheme of the background model used to generate random data for assessing the randomness of the triads. **(A)** The upper panel displays the shift value in megabases generated for each chromosome at every step (from -10 to 10 Mbp). Then, the ChIP-Seq peaks on each chromosome are shifted by the corresponding number of megabases. Peak must remain in the same type of chromatin compartment. If the shift causes the peak to move to the opposite compartment type, it is relocated to the nearest compartment of the required type. The bottom panel presents proteins as lists of RNAs that proteins interact with, based on real RNA peak data. For each protein, a pool of ‘related’ and ‘unrelated’ proteins is defined by the distance between their RNA lists. The pool of PR-interaction peaks for a particular protein is formed from the set of all peaks of ‘related’ proteins (including the protein’s original PR-peaks themselves) and the set of peaks of ‘unrelated’ proteins in a 4:1 ratio. This preserves the number and RNA biotype composition of the original RNA-protein peaks. The triad assembly pipeline is fed with resulting data, and this process is repeated 10 000 times to generate a sample of sufficient magnitude for drawing meaningful statistics. The number of simulated triads for proteins functionally described as transcriptional regulators in the K562 cell line is shown in the distributions in **(B)**, with the real triad counts denoted by vertical red lines. For information on other protein groups and all cell lines, see **Suppl. Fig. S4, S5, S6. (C)** Barplots displaying the number of simulated triads for each functional group of proteins.

Regarding PR-interactions, we aimed to preserve the biotype-specific content of the PR interaction repertoire for each protein by considering its relationships with other proteins in the dataset. This implies that a protein’s shuffled PR-interactions should be estimated by the set of PR-interactions of proteins that share numerous qualitative PR-interactions with it. As experimental data defining PR-interactions may contain some level of noise, we also included PR-interactions from other proteins to account for this (**Figure 3A**, lower panel). The dataset’s proteins were considered as the set of peaks of their PR-interactions and the list of RNAs on which their peaks are located. A distance matrix was obtained based on the magnitude of intersections of the RNA lists between each pair of proteins. Proteins whose distance to some protein was in the top third of all distances to that protein were labelled as ‘related’ for that target protein. The pool of PR-contact peaks for each protein was then formed as follows: The PR-peaks of both ‘related’ and ‘unrelated’ proteins were merged into separate pools. From these, a pool of shuffled PR-peaks for the target protein was drawn with replacement in a 4:1 ratio, while preserving the total number and RNA biotype composition of the original peak set. It is important to note that the protein ‘closeness’ obtained from RNA lists resembles the actual protein clustering by function (**Suppl. Fig. S2**). The data of DNA- and RNA-protein interactions were shuffled and then passed to a triad constructing pipeline for analysis (**Figure 3A**, right of both panels).

The triad counts on shuffled data from the background model are lower than those on real data for most of the proteins, as shown in **Figure 3B-C** (with *p*-values for each protein provided in **Suppl. Table S15**). The only exceptions are the triads based on data for hnRNPL, YBX3, KHSRP, RBFOX2, SUZ12 (K562 cell line), and WDR5 (mESC cell line). For YBX3 and KHSRP proteins, this could be due to unreliable RNA and/or DNA interaction data, which may contain a lot of noise. It is important to note that in some simulations, the number of triads obtained for hnRNPL, RBFOX2, and SUZ12 was even higher than the actual number. This may indicate either significant noise in the data or the presence/absence of some association of these proteins with the RNA-chromatin interactome. The behaviour of the number of simulated triads for the WDR5 protein appears to be influenced by the quality of the original ChIP-Seq data. The analysis of strand cross-correlations reveals a poor pattern of read shift correlation, indicating noisy data (see **Suppl. Fig. S3**).

Taken together, the simulation results demonstrate that the data obtained from real triads is statistically significant. It would be practically unattainable to obtain such a number of triads if the nature of pairwise interactions were different.

### RADICL-Seq NPM: protein absence affects the construction of triads

Formaldehyde crosslinking primarily fixes protein contacts. Therefore, RD-contacts in the RADICL-Seq NPM [9] experiment, where samples were treated with proteinase K before RNA-DNA ligation, can serve as a negative control for triads. The resulting triads can then be compared to triads obtained from RD-contacts constructed from standard 2% formaldehyde (2FA) crosslinking.

In the case of NPM, the triads will only reflect noise since there are no proteins present. We analysed RDt-contacts of EZH2, hnRNPK, SUZ12, and WDR5 proteins. For SUZ12, we used triads constructed from fRIP and eCLIP data (see section **”Pairwise interaction data can influence the construction of triads”**).We compared common triads derived from these data (**Figure 4A-C**, **Suppl. Fig. S7**) by selecting those in which the DNA parts overlap and the RNA parts belong to identical transcripts. The metrics obtained indicate that the SUZ12 and EZH2 proteins have overlapping contacts, as they are both functional units of the PRC2 complex. Additionally, the fRIP and eCLIP data for SUZ12 show a significant number of shared contacts. In contrast, the WDR5 protein in both 2FA and NPM data exhibits a large number of common contacts with other proteins. However, this effect disappears in the 2FA data with contacts common with NPM data removed. The correlation of DNA parts was also examined using StereoGene ([37]) and a similar pattern was observed (**Figure 4D-F**)).

**Fig. 4.**
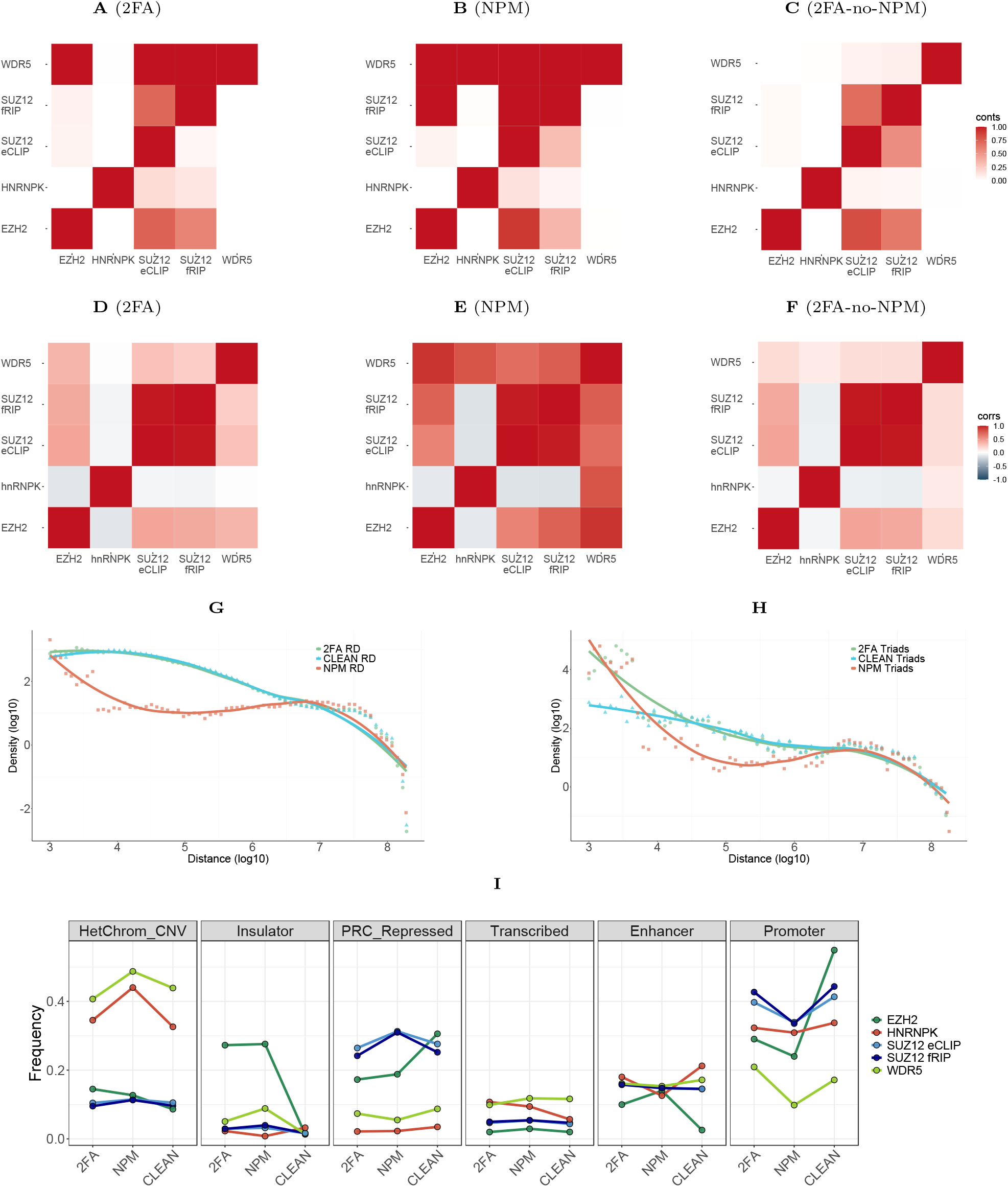
The RADICL-Seq data for various proteins and experimental phases, namely 2FA, NPM, and 2FA-no-NPM (representing triads present in 2FA but not in NPM). **(A-C)** The ratio of shared contacts in triads for the RADICL-Seq 2FA experiment **(A)**, RADICL-Seq NPM experiment **(B)**, and RADICL-Seq 2FA without RADICL-Seq NPM **(C)** is shown. The heatmaps may appear asymmetrical because of the normalization process by RDt-contacts set magnitude. Values greater than 1 are coloured the same as those equal to 1. **(D-F)** Correlation coefficients of DNA-parts tracks of the contacts for the RADICL-Seq 2FA experiment **(D)**, RADICL NPM experiment **(E)**, and RADICL-Seq 2FA without RADICL-Seq NPM **(F)** are shown. **(G, H)** RDt-scaling plots for cis-contacts of the RADICL-Seq experiment **(G)** and for RADICL-Seq-based triads **(H). (I)** Ratios of ChromHMM states distribution of DNA-parts for RADICL-Seq 2FA, RADICL-Seq NPM and RADICL-Seq 2FA without RADICL-Seq NPM experiments.

The NPM data show a different RD-scaling pattern, peaking at distances up to 1 Kb and then declining to uniformly distributed values as the distance from the gene increases. This effect seems to be partially present in the RADICL-Seq 2FA data as well, and is eliminated if the data are cleaned from contacts shared between the RADICL-Seq 2FA and RADICL-Seq NPM experiments (**Figure 4G-H**).

The distribution of DNA-parts over ChromHMM annotation states is of interest, as shown in **Figure 4I**. Different trends can be observed in the ratio change of DNA parts falling into a certain ChromHMM state. A pattern arises where this ratio changes drastically when filtering 2FA-based contacts from those shared with NPM-based dataset for some states and some proteins. For further information, see section **”Constructed triads are associated with genomic annotations”**.

The NPM data contains more contacts in total, which suggests low specificity and potentially higher noise. This observation is supported by the number of simulated triads distribution for RADICL-Seq 2FA and RADICL-Seq NPM data (see **Suppl. Fig. S8**). To increase the specificity of the data, subtracting the NPM contacts from the 2FA data is necessary.

### Pairwise interaction data can influence the construction of triads

The availability of data for various pairwise interactions of RNA, protein and DNA allows us to compare the triads that can be constructed from them. As a source of RNA-protein interactions, we used data from different experimental protocols, namely fRIP-Seq and eCLIP-Seq. Additionally, GRID-Seq and RADICL-Seq data were obtained for the same cell line, mouse embryonic stem cells (mESC). The sets of triads constructed from the different data were compared. RADICL-Seq 2FA data were used for this analysis. The data from the NPM experiment were filtered out (as described in the **”RADICL-Seq NPM: protein absence affects the construction of triads”** section).

The triads for the SUZ12 protein, constructed from different initial RNA-protein interaction data, are consistent. There are more RNAs forming triads in a shared set of contacts than in either of the two separate datasets (**Figure 5A**). In addition, the volume of the shared contacts set is greater (**Figure 5B**). We compared the biotype content of RNAs involved in the fRIP- and eCLIP-based triads (**Figure 5C**). Comparison of RNA biotype representation in triads revealed that RNAs from eCLIP-based triads are primarily ribosomal, whereas RNAs in fRIP-Seq-based triads exhibit greater diversity in composition.

**Fig. 5.**
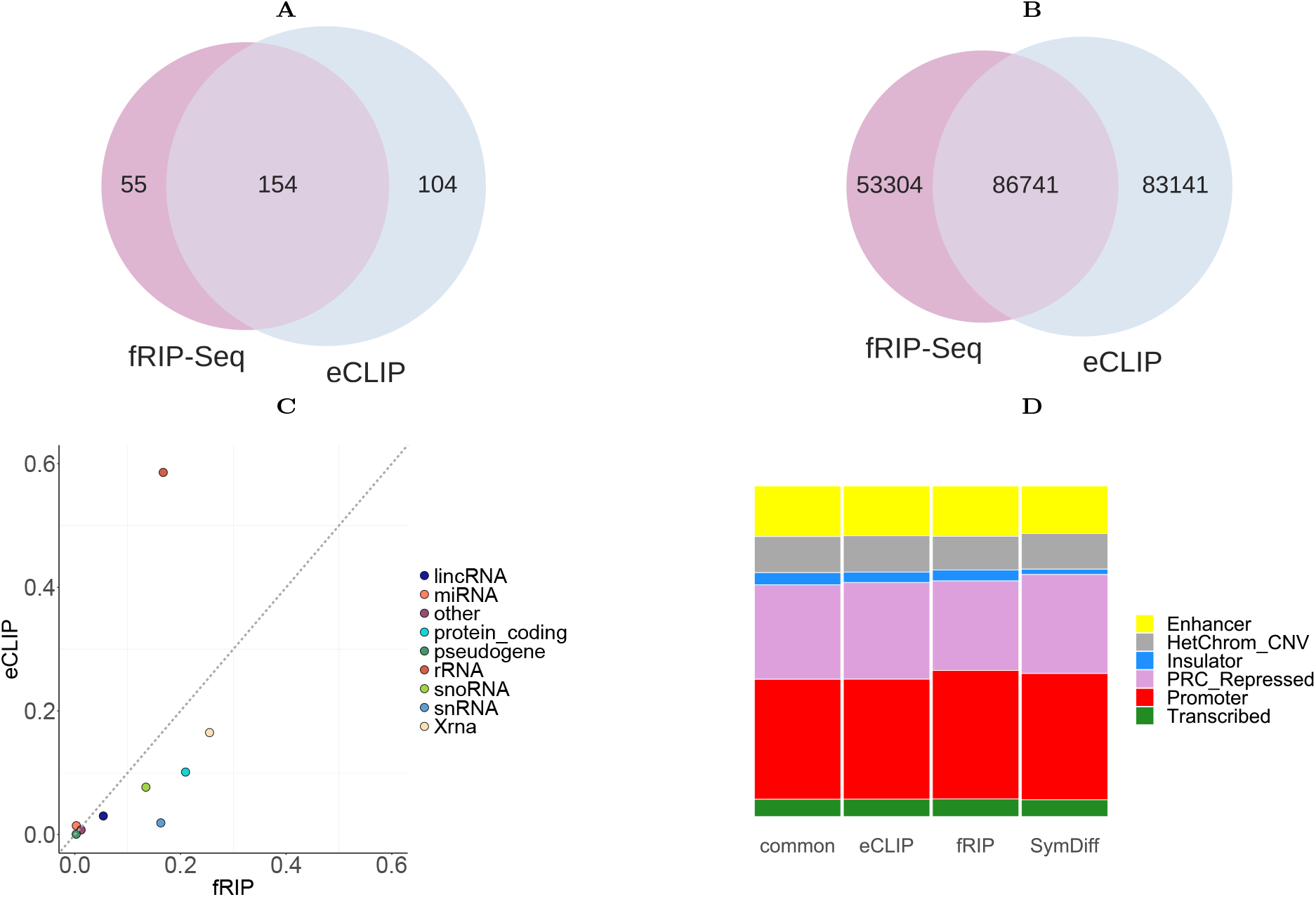
Comparison of SUZ12-mediated triads based on eCLIP- and fRIP-Seq PR-data. **(A, B)** Euler’s diagrams for common RNAs **(A)** and shared RD-contacts **(B)** for these datasets (a contact from one dataset is considered shared if its DNA-part intersects with any DNA-part of at least one contact in the opposing dataset and RNA-parts of these contacts belong to the same transcript). **(C)** RNA biotype composition comparison of the transcripts from which the RNA parts of triads originated. **(D)** Ratios of ChromHMM states distribution for DNA-parts of SUZ12-mediated triads based on eCLIP and fRIP-Seq data.

The RDt-contacts of fRIP- and eCLIP-based triads were divided into four groups:

- RDt-contacts of fRIP-based triads;
- RDt-contacts of eCLIP-based triads;
- RDt-contacts shared between fRIP- and eCLIP-based triads;
- the union of RDt-contacts only present in either fRIP-or eCLIP-based triads (essentially the symmetrical difference of the first two sets).

The relationship between the DNA-parts of these RDt-contacts and the ChromHMM annotation states was examined (**Figure 5D**). The resulting distributions show that although the RNA forming the triads are less consistent between fRIP- and eCLIP-based triads, the DNA loci they interact with and the loci left when constructing triads are consistent.

The aim of this comparison is to determine whether triads constructed on eCLIP or fRIP-Seq data are more biologically relevant. The choice of experimental protocol for constructing triads should be based on this criterion. Assuming that many regulatory functions of non-coding or unannotated RNAs are of great interest for further investigation, we selected triads with a higher proportion of biotypes of long non-coding RNAs and XRNAs. XRNAs are consistently assembled from RNA contacts mapped to unannotated genomic regions in gene deserts. Based on the stated condition, the optimal choice among all the sets is the triads shared between those constructed using fRIP-Seq and eCLIP (**Suppl. Fig. S9**). This is in accordance with common sense, as using a consensus of two experiments adds validity to the objects in that consensus, thereby increasing confidence in data obtained from noisy experiments. An additional argument in favour of the already demonstrated consistency of the discussed triads is the high correlation between the tracks of their DNA parts. This correlation coefficient of 0.89 (*p*-value *∼* 0), obtained using StereoGene, is supported by the cross-correlation function declining within 5 Kb (**Suppl. Fig. S10**).

We compared the effects of different types of RNA-chromatin interactome sequencing experiments on the construction of the RDt-contacts set. We used the triads of EZH2, hnRNPK, SUZ12 and WDR5 proteins constructed from GRID-Seq and RADICL-Seq RD-interactions data in cells of the mESC cell line. The RNA parts in triads of each protein were not consistent, as there were no common RNA parts between them. This was observed in both data filtered from NPM contacts (2FA-no-NPM) and the original 2FA data (**Suppl. Fig. S11**). Previous research has shown that all-to-all RD-data, specifically its DNA parts, begin to exhibit consistency over relatively large distances (starting from 100 Kb) for different experimental protocols [38]. These RNA-chromatin interactions may not represent the complete and accurate sample of functional RNA contacts at DNA loci due to their noisy nature and significant incompleteness. To address this, we extended the DNA parts of the contacts by 500 kb upstream and downstream from the 5’- and 3’-ends of the DNA part of the contact. We then used StereoGene to verify the consistency of these extended intervals (**Figure 6A-D**). We observed a significant correlation of DNA part tracks for EZH2 and hnRNPK proteins (**Table 3**). However, the distribution of correlation coefficients is bimodal, with modes at 0 and 1. This indicates that there are consistent regions between the GRID- and RADICL-Seq DNA parts, which are of main interest to study in light of triads construction. The remaining contacts, tending towards 0 and not resulting in a non-zero correlation, appear to be noise. However, it is important to note that RNA-chromatin interactome data is incomplete, and uncorrelated loci may indicate a contact that is detected in one experiment but lost in another. The most reliable loci are those with a high correlation between data from both experiments. Regrettably, due to the limited number of WDR5 protein-mediated RDt-contacts and the quality of the PR- and PD-interactions data, it is not possible to draw any meaningful conclusions regarding the consistency of triads mediated by this protein.

**Table 3.** The correlation between DNA parts of triads tracks for GRID-Seq and RADICL-Seq-based triads.

| Protein | Correlation | *p*-value |
| --- | --- | --- |
| EZH2 | 0.6525 | $2.7 * 10^{-50}$ |
| hnRNPK | 0.6597 | $1.6 * 10^{-3}$ |
| SUZ12 | 0.039 | $1.1 * 10^{-243}$ |
| WDR5 | -0.078 | 0.6724 |

**Fig. 6.**
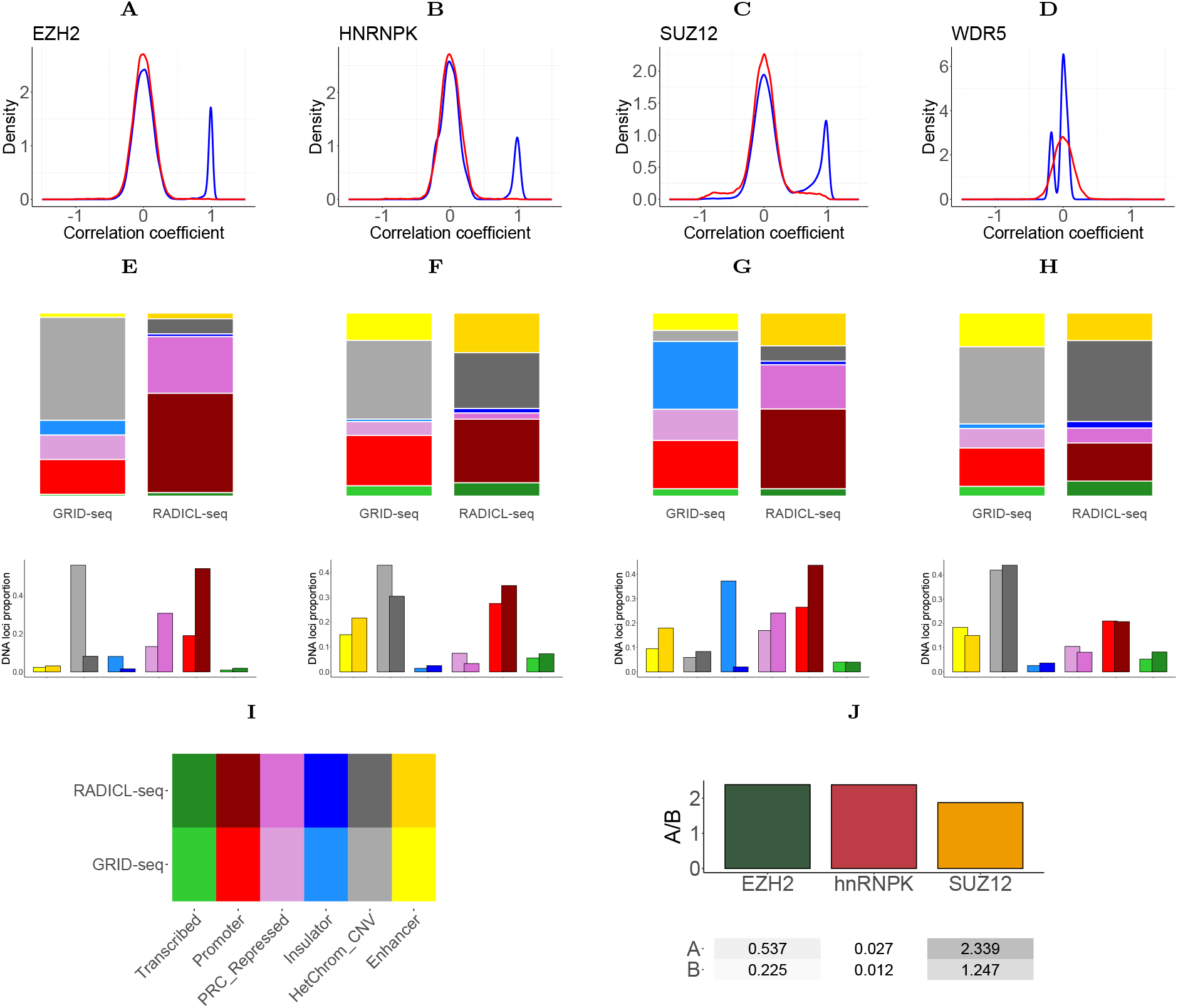
Comparison of EZH2, hnRNPK, SUZ12 and WDR5 protein triads constructed from RADICL- and GRID-Seq RNA-chromatin interaction data. **(A-D)** The correlation coefficient distribution between the extended DNA parts of triads for EZH2 **(A)**, hnRNPK **(B)**, SUZ12 **(C)** and WDR5 **(D)** proteins. **(E-H)** The distribution of DNA-part proportions of triads on ChromHMM states for triads constructed on RADICL-Seq and GRID-Seq RDt-data for EZH2 **(E)**, hnRNPK **(F)**, SUZ12 **(G)** and WDR5 **(H)** proteins. **(I)** A color legend for previous figures E-H. **(J)** The density of consistent triads in RADICL-Seq and GRID-Seq falling into A/B compartments.

The distribution of DNA parts by ChromHMM states was characterized by RDt-contacts constructed on RD-contacts data from RADICL-Seq and GRID-Seq experiments and PR-/PD-contacts for EZH2, hnRNPK, SUZ12 and WDR5 proteins (**Figure 6E-H**). The distributions showed higher consistency when comparing GRID-Seq and RADICL-Seq based triad DNA parts.

The regions that were consistent between the two experiments were characterised in terms of their A/B-compartments annotation. It was demonstrated that the density of triads falling into A-compartments is approximately twice that of triads falling into B-compartments. Density is defined as the number of DNA loci within A-or B-compartments normalized to the total compartment length (**Figure 6J, Table 4**). This suggests that when the chromatin is more open at the site of contact ligation, the contact representation is more likely to be consistent across different protocols. It is also possible that GRID-Seq captures more consistent RD-contacts due to their proportion of total GRID-Seq contacts.

**Table 4.** Ratio of contacts falling into regions of high correlation based on StereoGene results. cnts - contacts.

| Protein | # consistent cnts | % RADICL cnts (total) | % GRID cnts (total) |
| --- | --- | --- | --- |
| EZH2 | 910 | 3.79 (23981) | 21.96 (4144) |
| hnRNPK | 46 | 2.9 (1585) | 27.22 (169) |
| SUZ12 | 4348 | 1.84 (236673) | 1.91 (227292) |

The validation of results from methods used to capture pairwise protein, RNA and DNA interactions is a critical issue. This is particularly true for the RNA-chromatin interactions data generated by different protocols. Although unfiltered all-to-all data has been described as inconsistent [38], our findings suggest that there is a proportion of all-to-all data that is consistent between various protocols. The triad construction method we propose may extract this data.

### Constructed triads may prove orthologous

The presence of conserved RNA-DNA interactions between organisms provides strong support for our triad filtering method. We tested for conserved interaction triads between organisms by comparing the interaction triads constructed from EZH2, hnRNPK, SUZ12, and WDR5 PR- and PD-interaction data for human (K562) and mouse (mESC) cell lines, using available data. We searched for RNA orthologues that form triads in both mouse and human as bidirectional hits with a relaxed significance threshold (**Suppl. Table S16**) using ortho2align [35]. We found 11 such RNA pairs. Next, we lifted over the DNA loci in contact with these RNAs to the opposite genome and searched for the closest DNA segment of the orthologous RNA contact. The distances between the DNA fragments and the DNA fragments in the original genome are substantial (**Suppl. Table S17**). However, we identified 10 pairs of orthologous RNA contacts (**Table 5**) whose DNA fragments are within 2 Mb of one another. Considering the previously mentioned experiment specifics and the issue of inaccurate genomic interval lifting between genomes, this result confirms the reliability of our obtained data.

**Table 5.** Top 10 potentially orthologous protein-mediated RNA-DNA contacts with the closest distance. Distance is defined as the distance between the locus in the mouse genome and the locus lifted from the human genome to the mouse genome or vice versa.

| DNA Chr | Mouse RNA | Human RNA | Distance | Protein | Closest gene |  |
| --- | --- | --- | --- | --- | --- | --- |
|  |  |  |  |  | Mouse gene | Human gene |
| chr16 | Gm24299 | SNORD5 | 71 101 | SUZ12 | Lrrc74b, P2rx6 | XR_933144.3, XYLT1 |
| chr22 | Gm24299 | SNORD5 | 79 851 | SUZ12 | Pi4ka, Serpind1 | AIFM3 |
| chr16 | Gm24265 | RNU4-2 | 114 378 | EZH2 | Rimbp3, Hic2 | XYLT1 |
| chr11 | Rpph1 | RPPH1 | 334 071 | hnRNPK | Tex14, Rnu3b4 | TMEM135 |
| chr16 | Gm24265 | RNU4-2 | 409 387 | SUZ12 | Rimbp3, Hic2 | No close genes |
| chr11 | Gm24265 | RNU4-2 | 627 098 | SUZ12 | Rnf213 | No close genes |
| chr2 | Rpph1 | RPPH1 | 1 529 017 | EZH2 | Gm40034 | No close genes |
| chr10 | Gm24299 | SNORD5 | 1 575 789 | SUZ12 | Cyp27b1, Mettl1, Marchf9 | No close genes |
| chr12 | Gm24299 | SNORD5 | 1 722 366 | SUZ12 | Ttc6 | RPS26 |
| chr19 | Gm24265 | RNU4-2 | 1 920 768 | SUZ12 | Ms4a4a | CACNA1A |

The limited number of orthologous triads and their composition can be explained. Firstly, we analysed significantly different cell types, as there were no better datasets available for similar cell types. Additionally, data obtained for RNA-chromatin contacts is rather noisy and incomplete. Therefore, consistent orthologous triads may not be found. Lastly, non-coding RNAs evolve rapidly, so corresponding orthologs may simply be missed. Therefore, the orthologous triads found are only for RNAs with a large number of contacts.

### Constructed triads are associated with genomic annotations

The RNA-chromatin interactome data is filtered by triad construction, which also selects RNA-associated PDt-contacts (see also **Suppl. Table S10, S11, S12, S13, S14**). We analyzed the annotation of triads PDt-contacts by different chromatin states and compared them with the annotation of ChIP-seq peaks of the same proteins in the K562 cell line (**Figure 7A-C**, **F**; for all proteins see **Suppl. Fig. S12**). We used the 𝒳^2^ criterion to statistically assess the difference in enrichment (see **Suppl. Table S18** for test results for all proteins).

**Fig. 7.**
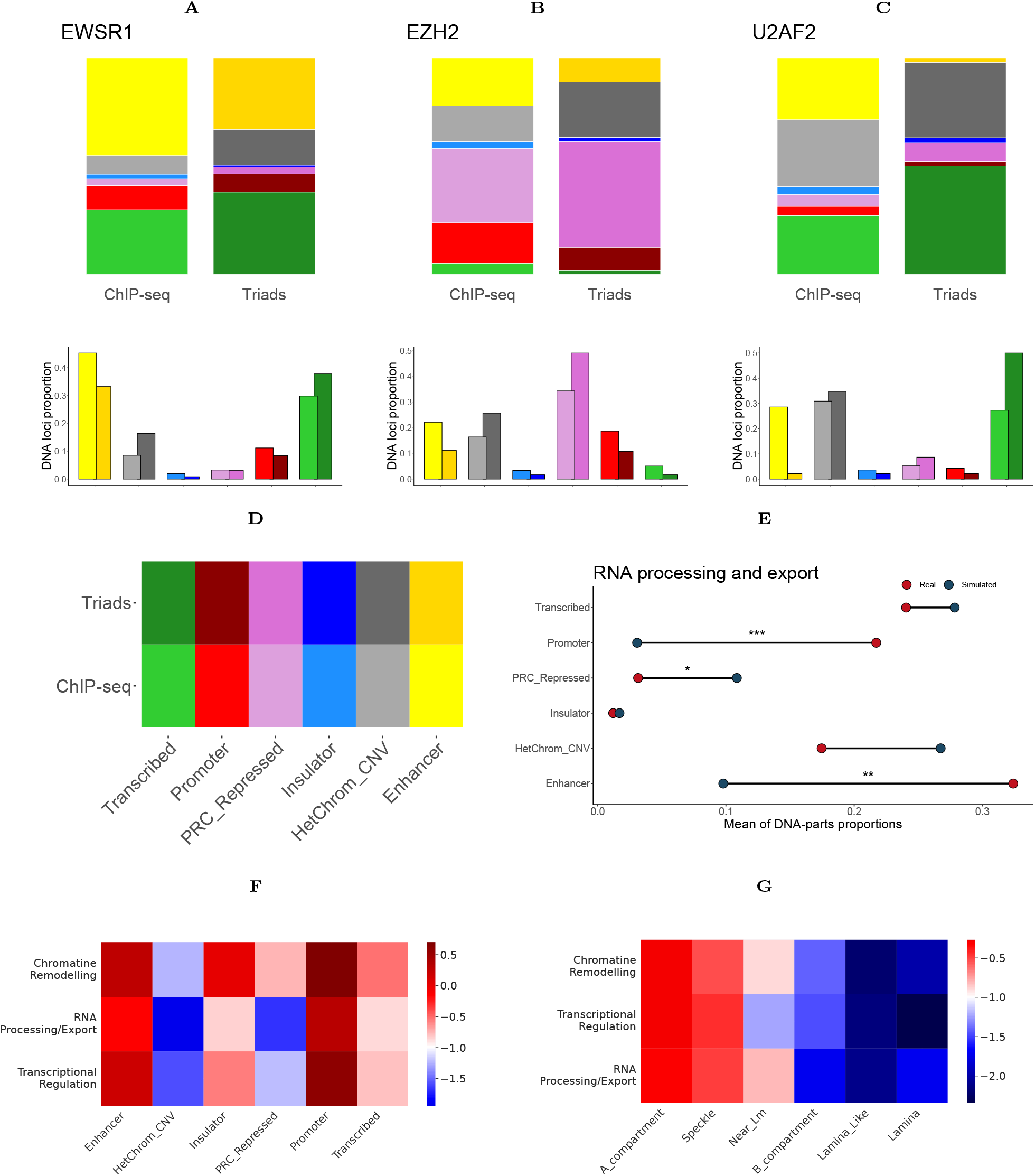
Triads and ChromHMM/SPIN genome annotation. **(A-C)** The distribution of DNA parts of triads mediated by EWSR1 **(A)**, EZH2 **(B)**, and U2AF2 **(C)** proteins across ChromHMM states is presented. **(D)** The figures are accompanied by a color legend. **(E)** The distribution of DNA parts of triads by ChromHMM states constructed on real and simulated DNA- and RNA-protein interaction data for RNA processing and exporting proteins. **(F, G)** Heatmaps for the distribution of DNA parts of protein triads, averaged by functional groups, over ChromHMM **(F)** and SPIN **(G)** annotations. *−* log_10_ of normalized densities is shown. The proportion of DNA parts in the annotation state is calculated by dividing the total length of the annotated regions in nucleotides by the total length of the sequence.

For proteins that do not participate in regulating certain RNAs and their interactomes, it is reasonable to assume that the distribution of ChromHMM states in DNA regions will remain unchanged. This is because the process of constructing triads for these proteins essentially involves randomly sampling from ChIP-Seq loci. However, this may not be the case for proteins involved in RNA-interactome reactions. For certain proteins, we can observe a notable enrichment pattern of states that correspond to the protein’s functional association (e.g. PRC2 repressed chromatin for EZH2 and SUZ12, or transcribed chromatin for SRSF9 or U2AF2). This pattern is in addition to significant statistical test results. The distribution of the DNA parts of triads constructed on real data was compared to the distribution of the DNA parts of triads constructed on our simulated data (**Figure 7E**, see **Suppl. Fig. S14, S15** for all proteins and SPIN annotation.

The proteins were divided into two groups based on the median *p*-value of the 𝒳^2^ goodness-of-fit test to determine if there were any differences in the characteristics of RDt-contacts mediated by the two groups. To address the bias in contact density in the RNA-chromatin interactome, the weight of the contact was defined as:

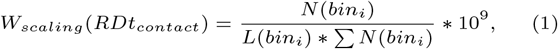

where *N* (*bin*_*i*_) represents the number of contacts of the target RNA falling into a bin, and *L*(*bin*_*i*_) is the bin length. To estimate the difference between two groups, we introduce a significance metric for triads as the harmonic mean of q-values, HMQ **(2)**:

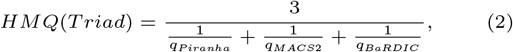

a metric defined as the aggregated significance — q-values — of all pairwise peaks from which the interaction triad is assembled. We used HMQ to estimate importance weights of the contacts.

The comparison of contact weights and significance revealed differences in the significance distributions of the triads. Triads mediated by proteins significantly associated with the RNA-chromatin interactome showed greater significance. However, the distribution of contact scaling weights for the two groups was not different (**Figure 8A, 8B**), indicating that while grouping proteins into two classes helps differentiate more significant contacts, the underlying nature of RDt-contacts present in both groups is the same. The proposed division of proteins based on their association with potential functions of mediating RNA-interactome regulation may be useful for investigating functional regulatory processes in the nucleus.

**Fig. 8.**
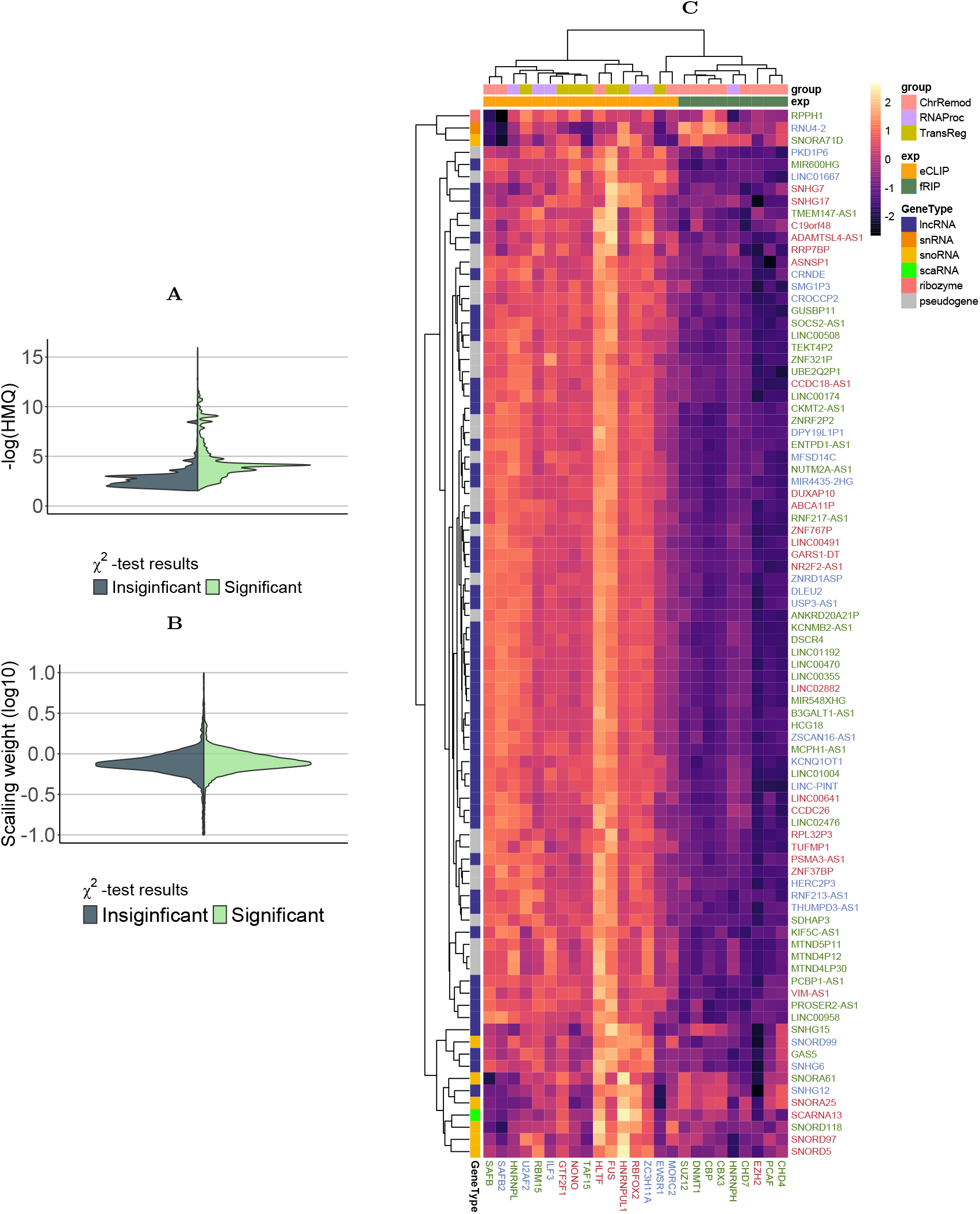
The association of proteins with RD-contacts. **(A)** Differences in HMQ significance between triads formed by proteins that were deemed significant by 𝒳^2^ goodness-of-fit test and those that were not. **(B)** Difference in scaling weights for mentioned groups of proteins. **(C)** A heatmap clusterisation of RNA and proteins by PR-contact fold changes.

To determine the specificity of PR contacts, we identified RNAs that form triad interactions with at least 20 proteins (**Suppl. Fig. S16**). We referred to these RNAs as the ‘common’ RNAs, while the remaining RNAs were referred to as the ‘specific’ RNAs. Next, we divided all RD-contacts from the original RD-dataset into four classes for each protein:

- the RNA in the contact is specific and the DNA part of the contact overlaps with at least one ChIP-Seq locus of the protein;
- the RNA in the contact is common and the DNA part of the contact overlaps with at least one ChIP-Seq locus of the protein;
- the RNA in the contact is specific and the DNA part of the contact does not overlap with any ChIP-Seq locus of the protein;
- the RNA in the contact is common and the DNA part of the contact does not overlap with any ChIP-Seq locus.

Fisher’s exact test was applied to the contingency table obtained from these categories to assess the ability of each protein to form specific PR-contacts (**Suppl. Table S18**).

A group of 25 proteins associated with the RNA interactome was selected based on the consensus of two tests. RNAs forming triads with these proteins were also selected. Each RNA was represented as a vector of its interaction signals with these proteins, defined as the fold change of PR-contact. The vectors of both the RNAs and proteins were then clustered. The results indicate (**Figure 8C**) that proteins cluster based on their functional roles as chromatin remodelers, transcription regulators, and RNA processing and splicing regulators. Similarly, selected RNAs are grouped by function, with snoRNA and lncRNA clustering separately.

The identified proteins associated with the RNA-chromatin interactome may have functionally relevant RNA-chromatin interactions. A detailed analysis of their contacts could prove useful.

### RD-contacts precipitation on CTCF produces results similar to triads

The Red-ChIP experiment is of interest as it validates the interaction triads constructed for the CTCF protein in K562 cells. The experiment involves immunoprecipitation of RD-complexes on the protein. It is important to understand how the results of this experiment relate to the results obtained from constructing the triads on the Red-C experiment RD-contacts data. DNA parts from the following contact groups were correlated using StereoGene: RD-contacts from the Red-C experiment, input from the Red-ChIP (essentially another Red-C replicate), RD-contacts from those experiments after filtering the data with CTCF peaks, RD-contacts precipitated on CTCF (RedChIP Signal), and DNA parts of the interaction triads that were constructed. The consistency of the sets of interacting DNA loci is high between real and predicted CTCF-mediated RNA-DNA contacts (**Figure 9A**).

**Fig. 9.**
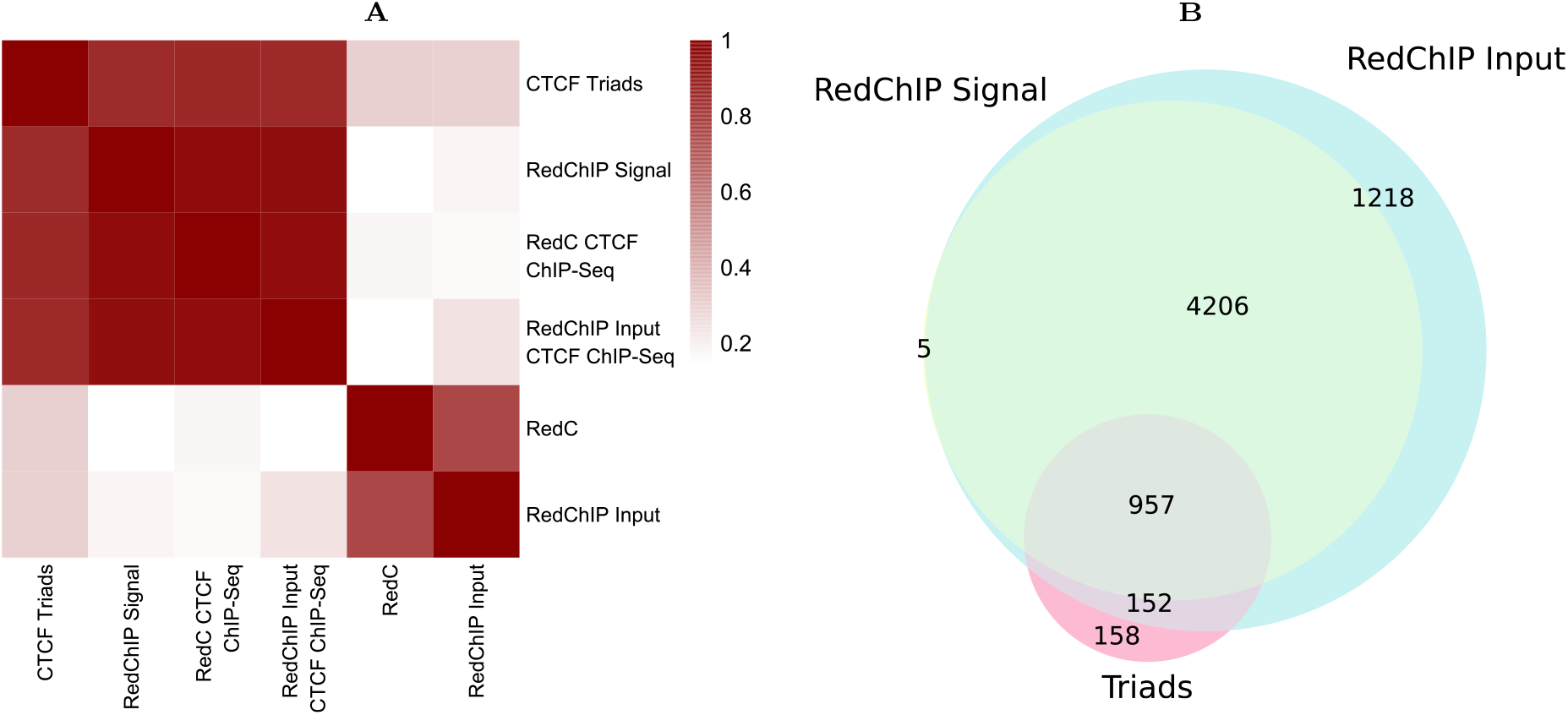
Red-ChIP data supports with predicted triads. **(A)** Stereogene correlations of DNA-parts of triads. **(B)** Shared RNAs between Red-C-based CTCF triads, Red-ChIP input data and Red-ChIP signal data.

It is clear that the data filtered by ChIP-Seq are well-clustered separately from the unfiltered RD-contacts of the Red-C experiment, alongside the Red-ChIP input data. The set of RNAs involved in triads interactions and in the RedChIP RD-contacts overlap well (**Figure 9B**). These results suggest that the computationally predicted RD-contacts in triads represent a subset of real contacts, as the RD-contacts data from the Red-ChIP experiment also exhibit some level of noise.

## Discussion

RNA-chromatin interactome data is known to be noisy in sequence bioinformatics and requires careful processing to filter out nonspecific and noisy contacts. We carefully studied and processed the data of pairwise interactions to ensure that only relevant and specific contacts were included. We used a range of statistical peak calling methods designed for pairwise interactions between RNA, DNA, and proteins to search for functional contacts of the RNA-chromatin interactome mediated by one of the 47 chromatin proteins, using data of PR- and PD-interactions. Our approach enables us to choose a significance level for each level of pairwise interactions used to construct the object of study based on an accumulated significance metric. This significance selection also applies to triads that have already been constructed.

Despite the high levels of noise and large number of cis-contacts and contacts formed by protein-coding RNA in the RD-contacts data, our approach appears to effectively filter out this flawed data. Filtering out too much data can have drawbacks, and it is important to consider statistical assessment of thresholds to work with the data accurately.

The obtained data are meaningfully different from those obtained by chance and contain less noise than the original RNA-chromatin contacts. There are (**Suppl. Table S16**) a significant number of conserved triads between mouse and human organisms, and the functional validation of these interactions remains a question.

We demonstrated that RADICL-Seq NPM phase data can serve as a negative control for the triads construction method, which is a protocol for non-protein-mediated RNA-chromatin contacts. Although the nature of these NPM data is not fully understood, they are useful when searching for RDt-contacts mediated by a particular protein, rather than determining the genome-wide RNA-chromatin interactome. It is demonstrated here that the correlation between RDt-contacts mediated by different proteins decreases when the contacts shared between RADICL 2FA and RADICL NPM sets are filtered out.

We investigated whether there are differences in the constructed triads when starting with data from different pairwise interaction detection protocols for PR-interactions (fRIP-Seq and eCLIP-Seq) and for RD-interactions (namely, GRID-Seq and RADICL-Seq). Triads built on different datasets appear consistent for PR-contacts and almost consistent for RD-contacts. A range of regions can be identified that is conserved between experiments. To draw more data, other types of pairwise interactions need to be incorporated into the analysis. It might be valuable to see if PD-contacts originating from different experiments produce consistent triads for target proteins.

We demonstrated that constructing interaction triads for chromatin proteins enhances RNA-chromatin contacts in regions where DNA loci functional annotation inferred from ChromHMM is associated with protein function. For instance, EZH2-mediated triads DNA-parts are more abundant in PRC2-repressed chromatin regions than the ChIP-Seq loci of this protein. In addition, we verified that the randomly simulated triads were not related to protein function, in the sense of ChromHMM. Triads with a DNA locus located in chromatin states that correspond to protein functions are of interest for studies on regulatory interactions. Additionally, we observed that the enrichment of DNA parts in triads is higher in open chromatin states such as Promoter, Enhancer, Transcribed states from ChromHMM annotation or A-compartment and Speckle states from SPIN annotation compared to other states. The annotation of A/B-compartment of triads is consistent between RADICL-Seq and GRID-Seq experiments, supporting the hypothesis that open chromatin-RNA contacts are better represented in the interactome data.

In addition, the interaction triads constructed are consistent with the results of the Red-ChIP experiment, which serves as a positive control for our approach. While the Red-ChIP experimental procedure allows for the presence of contacts of noisy origin, the number of RNAs shared between the set of RD-contacts of Red-ChIP and RDt-contacts is high, and the set of DNA loci with which these RNAs interact is also shared.

It has been demonstrated that the proposed approach is effective. The main challenge in identifying RNA-chromatin interactions mediated by intranuclear proteins is the lack of relevant data. In this study, we analysed various biological data sources, including ENCODE, NCBI GEO, ReMap, 4D Nucleome, and GTRD databases, and identified a dataset of triads of 47 proteins as successful. To scale this approach, machine learning techniques can be used for data imputation.

## Acknowledgments

The authors thank Asya Mendelevich for their valuable comments on manuscript. The research was supported by RSF (project No. 23-14-00136).

