## Supplementary materials (Suppl. Tables S1-S19, Suppl. Figures S1-S11) for "Comprehensive analysis of RNA-chromatin, RNA- and DNA-protein interactions"


Table S1: Database identifiers for data used in the study. IDs correspond to databases the following way: GSE.../GSM... – NCBI GEO, ENCS... – ENCODE, PEAKS... – GTRD, ERP... – ArrayExpress. . . . . . 3
Table S2: BaRDIC running parameters for peak calling in all- and one-to-all RNA-DNA data. . . . . . 5
Table S3: Triad number magnitude when applying peak extension procedure. . . . . . 6
Table S4: Grouped ChromHMM annotation states. . . . . . 6
Table S5: Grouped SPIN annotation states. . . . . . 7
Table S6: StereoGene parameters changed from defaults when correlating DNA tracks . . . . . . 7
Table S7: Proteins split into three groups according to their putative function inferred from UniProt. . . 8
Table S8: Distribution of the of cis- and trans-contacts abundances in constructed triads for each protein. Unless stated otherwise, triads are based on eCLIP RNA-protein interactions data, Red-C RNA-DNA interactions data for K562 cells. Ratios of mRNA-mediated triads are also shown. . 8
Table S9: cis-/trans- contacts ratio statistics for selected RNAs for K562 Red-C experiment. . . . . . 11
Table S10: Triads and ChIP-Seq composition by ChromHMM annotation of DNA loci for all proteins split by function. *cnt* – number of observed ChIP-Seq loci in a state; *tri* – number of triads; $Z$ – z-score of $\frac{tri}{cnt}$ distribution over all ChromHMM states . . . . . . 12
Table S11: Triads and RD-contacts DNA parts composition by ChromHMM annotation of DNA loci for all proteins split by function. *cnt* – number of observed DNA-parts in a state; *tri* – number of triads; $Z$ – z-score of $\frac{tri}{cnt}$ distribution over all ChromHMM states . . . . . . 13
Table S12: Triads and ChIP-Seq composition by SPIN annotation of DNA loci for all proteins split by function. *cnt* – number of observed ChIP-Seq loci in a state; *tri* – number of triads; $Z$ – z-score of $\frac{tri}{cnt}$ distribution over all SPIN states . . . . . . 13
Table S13: Triads and and RD-contacts DNA parts composition by SPIN annotation of DNA loci for all proteins split by function. *cnt* – number of of observed DNA-parts in a state; *tri* – number of triads; $Z$ – z-score of $\frac{tri}{cnt}$ distribution over all SPIN states . . . . . . 14
Table S14: Triads and and RD-contacts composition by RNA biotypes for all proteins split by function. *cnt* – number of of observed RNA of a given biotype; *tri* – number of triads; $Z$ – z-score of $\frac{tri}{cnt}$ distribution over all RNA biotypes . . . . . . 14
Table S15: $p$-values for the real number of triads on the background distribution of the number of triads obtained on simulated data. The background distribution for triads number was fit using a gamma distribution, and the $p$-value defined as the probability of obtaining the real number of triads for protein or a value greater from the fit distribution. . . . . . 14
Table S16: RNA orthologs found by ortho2align (selected as bidirectional best hit) . . . . . . 17
Table S17: DNA contacts distances in original genomes for RNA orthologs. *"RNA from source genome"* stands for RNA, for which coordinates of DNA locus that it interacts with were unchanged. *"RNA for liftOver"* stands for RNA, for which coordinates of DNA locus were lifted over to the opposing genome. . . . . . 17
Table S18: Results for the association tests: $\chi^2$- goodness-of-fit of ChromHMM annotation of DNA-parts and Fisher's exact test for association of PR- and PD-contacts data of protein with RD-contacts. 20
Table S19: Results for the association test: $\chi^2$- goodness-of-fit of SPIN annotation of DNA-parts of triads and proteins ChIP-Seq loci. . . . . . 22
Figure S1: Circos diagrams for constructed protein mediated RNA-DNA interactions, or triads. Legend for diagrams is shown in the main text. . . . . . 28
Figure S2: Protein clusterization based on their "related" proteins . . . . . . 29
Figure S3: Strand cross-correlation for WDR5 ChIP-Seq. . . . . . 29
Figure S4: Ridgeline plot for number of simulated triads for chromatine remodelling proteins. . . . . . 30
Figure S5: Ridgeline plot for number of simulated triads for RNA processing and export proteins. . . . . . 31
Figure S6: Ridgeline plot for number of simulated triads for transcription regulating proteins. . . . . . 32
Figure S7: Triads number desaturation when filtering the 2FA contacts shared with NPM phase. . . . . . 33
Figure S8: Number of triads simulated for RADICL-Seq 2FA and RADICL-Seq NPM datasets differs significantly with the latter number being larger. . . . . . 33
Figure S9: Biotype content of RNAs forming triads in all fRIP-based triads **(A)**, all eCLIP-based triads **(B)**, triads shared by two experiments **(C)** and triads present only in one of datasets **(D)**. . . 34
Figure S10: SUZ12 DNA-parts tracks correlation. **(A)** Distribution of correlations across all bins; **(B)** Cross-correlation function decline. blue — observed correlation; red — background correlations 35
Figure S11: Common RNA between contacts of RADICL-Seq and GRID-Seq in triads formed by EZH **(A)**, hnRNPK **(B)**, SUZ12 **(C)** and WDR5 **(D)** proteins. . . . . . 35

Figure S12: Stacked barplots and paired barplots representing DNA parts distribution along the ChromHMM annotation states for triads of proteins in both cell lines. . . . . . . . . . . . . . . 39
Figure S13: Stacked barplots and paired barplots representing DNA parts distribution along the SPIN annotation states for triads of proteins in both cell lines. . . . . . . . . . . . . . . . . . . . . . . . . 43
Figure S14: DNA parts of triads distribution by ChromHMM states obtained using real and simulated DNA- and RNA-protein interaction data for RNA processing and exporting proteins. . . . . . 44
Figure S15: DNA parts of triads distribution by SPIN states obtained using real and simulated DNA- and RNA-protein interaction data. . . . . . . . . . . . . . . . . . . . . . . . . . . . . . . . . . . . . . . 44
Figure S16: RNA participation in triads, i.e. given an RNA that if forming triads, what is the possible range of proteins number it forms triads with? . . . . . . . . . . . . . . . . . . . . . . . . . . . . 45

**Supplementary Table S1.** Database identifiers for data used in the study. IDs correspond to databases the following way: GSE. . . /GSM. . . – NCBI GEO, ENCS. . . – ENCODE, PEAKS. . . – GTRD, ERP. . . – ArrayExpress.

| Protein | RNA-protein data | ChIP-Seq data |
|---|---|---|
| CBP | K562 fRIP: GSE67963 | K562: ENCSR000ATT |
| CBX3 | K562 fRIP: GSE67963 | K562: ENCSR000ATV |
| CHD1 | K562 fRIP: GSE67963 | K562: ENCSR000AQD |
| CHD4 | K562 fRIP: GSE67963 | K562: ENCSR000ATL |
| CHD7 | K562 fRIP: GSE67963 | K562: ENCSR000AVD |
| CTCF | K562 fRIP: GSE67963 | K562: ENCSR000AKO |
| DNMT1 | K562 fRIP: GSE67963 | K562: ENCSR987PBI |
| EWSR1 | K562 eCLIP: ENCSR887LPK | K562: ENCSR142YYA |
| EZH2 | K562 fRIP: GSE67963; mESC eCLIP: GSE141700 | K562: ENCSR000AQE; mESC: PEAKS042326 |
| FUS | K562 eCLIP: ENCSR069EVH | K562: ENCSR051DXE |
| GTF2F1 | K562 eCLIP: ENCSR736AAG | K562: ENCSR000EHC |
| HDAC1 | K562 fRIP: GSE67963 | K562: ENCSR000AQF |
| HLTF | K562 eCLIP: ENCSR589YHM | K562: ENCSR090JNM |
| hnRNPC | K562 eCLIP: ENCSR249ROI | K562: PEAKS057732 |
| hnRNPH | K562 fRIP: GSE67963 | K562: GSE120104 |
| hnRNPK | K562 eCLIP: ENCSR268ETU; mESC eCLIP: GSE87233 | K562: ENCSR014RCS; mESC: GSE129471 |
| hnRNPL | K562 eCLIP: ENCSR795CAI | K562: ENCSR594BNR |
| hnRNPUL1 | K562 eCLIP: ENCSR571VHI | K562: ENCSR296MXW |

See next page

| Protein | RNA-protein data | ChIP-Seq data |
|---|---|---|
| ILF3 | K562 eCLIP: ENCSR438KWZ | K562: ENCSR632TJQ |
| KHSRP | K562 eCLIP: ENCSR438GZQ | K562: ENCSR686EYO |
| LARP7 | K562 eCLIP: ENCSR456KXI | K562: ENCSR288MOZ |
| LSD1 | K562 fRIP: GSE67963 | K562: PEAKS048159, PEAKS048160 |
| MORC2 | K562 eCLIP: ENCSR168VUE | K562: GSE95374 |
| NONO | K562 eCLIP: ENCSR861PAR | K562: ENCSR010KFT |
| NUP98 | K562 fRIP: GSE67963 | K562: PEAKS048332 |
| PCAF | K562 fRIP: GSE67963 | K562: ENCSR000ATZ |
| PCBP1 | K562 eCLIP: ENCSR922WJV | K562: ENCSR052PTN |
| PHF8 | K562 fRIP: GSE67963 | K562: ENCSR000AQH |
| PTBP1 | K562 eCLIP: ENCSR981WKN | K562: ENCSR948KMB |
| RBBP5 | K562 fRIP: GSE67963 | K562: ENCSR000AQI |
| RBFOX2 | K562 eCLIP: ENCSR756CKJ | K562: ENCSR822LBD |
| RBM15 | K562 eCLIP: ENCSR196INN | K562: ENCSR850WUE |
| RBM22 | K562 eCLIP: ENCSR295OKT | K562: ENCSR848AOP |
| SAFB | K562 eCLIP: ENCSR484LAB | K562: ENCSR072VUO |
| SAFB2 | K562 eCLIP: ENCSR943MHU | K562: ENCSR724GSW |
| SRSF1 | K562 eCLIP: ENCSR432XUP | K562: ENCSR832DMO |
| SRSF7 | K562 eCLIP: ENCSR468FSW | K562: ENCSR222MYK |

See next page

**Supplementary Table S1** – continued from previous page

| Protein | RNA-protein data | ChIP-Seq data |
|---|---|---|
| SRSF9 | K562 eCLIP: ENCSR381VNR | K562: ENCSR814AJH |
| SUZ12 | K562 fRIP: GSE67963; mESC eCLIP: GSE141700; mESC fRIP: ERP113531 | K562: ENCSR000AUC; mESC: PEAKS057610 |
| TAF15 | K562 eCLIP: ENCSR568DZW | K562: ENCSR047LSJ |
| TARDBP | K562 eCLIP: ENCSR720BJU | K562: ENCSR033VAZ |
| U2AF1 | K562 eCLIP: ENCSR862QCH | K562: ENCSR690GUG |
| U2AF2 | K562 eCLIP: ENCSR893RAV | K562: ENCSR479QAJ |
| WDR5 | K562 fRIP: GSE67963; mESC fRIP: GSE53035; | K562: PEAKS050903, PEAKS050904; mESC: GSM566279 |
| YBX3 | K562 eCLIP: ENCSR529FKI | K562: ENCSR567JEU |
| ZC3H11A | K562 eCLIP: ENCSR712IAG | K562: ENCSR000EFR |
| ZC3H8 | K562 eCLIP: ENCSR181NRW | K562: ENCSR494PWZ |

**Supplementary Table S2.** BaRDIC running parameters for peak calling in all- and one-to-all RNA-DNA data.

| Option | Value |
|---|---|
| `-min_contacts` | 1000 |
| `-trans_min` | 1000 |
| `-trans_max` | 1000000 |
| `-trans_step` | 100 |
| `-cis_min` | 1.1 |
| `-cis_max` | 2 |
| `-cis_step` | 0.01 |

See next page

**Supplementary Table S2** – continued from previous page

| Option | Value |
|---|---|
| `-cis_start` | 100 |
| `-tolerance` | 0.01 |
| `-window` | 1 |
| `-ifactor` | 0.01 |
| `-degree` | 3 |
| `-max_threshold` | 0.05 |
| `-fill_value` | 1 |
| `-qval_threshold` | 0.05 |
| `-bgtype` | custom |

**Supplementary Table S3.** Triad number magnitude when applying peak extension procedure.

| DNA extend both | RNA extend dowstream | TRIADS w mRNAs | TRIADS w/o mRNAs |
|---|---|---|---|
| 0 | 0 | 257 749 | 210 681 |
| 0 | 50 | 263 862 | 214 112 |
| 0 | 100 | 272 556 | 220 127 |
| 1000 | 0 | 1 271 470 | 1 038 752 |
| 1000 | 50 | 1 301 594 | 1 055 370 |
| 1000 | 100 | 1 343 721 | 1 084 776 |
| 2000 | 0 | 2 163 250 | 1 770 970 |
| 2000 | 50 | 2 214 383 | 1 799 471 |
| 2000 | 100 | 2 284 709 | 1 848 423 |

**Supplementary Table S4.** Grouped ChromHMM annotation states.

| Grouped State | Original ChromHMM State |
|---|---|
| Promoter | Active Promoter, Weak Promoter, Inactive/poised Promoter |
| Enhancer | Strong enhancer, Weak/poised enhancer |

See next page

**Supplementary Table S4** – continued from previous page

| Grouped State | Original ChromHMM State |
|---|---|
| Transcribed | Transcriptional transition, Transcriptional elongation, Weak transcription |
| Insulator | Insulator |
| PRC_Repressed | Polycomb-repressed |
| HetChrom_CNV | Heterochromatin; low signal, Repetitive/Copy Number Variation |

**Supplementary Table S5.** Grouped SPIN annotation states.

| Grouped State | Original SPIN State |
|---|---|
| A_compartment | Interior_Act1, Interior_Act2, Interior_Act3 |
| B_compartment | Interior_Repr1, Interior_Repr2 |
| Lamina | Lamina |
| Lamina_like | Lamina_like |
| Near_Lm | Near_Lm1, Near_Lm2 |
| Speckle | Speckle |

**Supplementary Table S6.** StereoGene parameters changed from defaults when correlating DNA tracks

| Option | Value |
|---|---|
| `Rscript` | 1 |
| `wSize` | 300000 |
| `wStep` | 300000 |
| `maxNA` | 99.9 |
| `maxZero` | 99.9 |
| `kernelSigma` | 2000 |
| `bin` | 200 |
| `outLC` | 1 |

**Supplementary Table S7.** Proteins split into three groups according to their putative function inferred from UniProt.

| Protein group | Grouped proteins |
|---|---|
| Chromatin modification and remodelling | CBP, CBX3, CHD1, CHD4, CHD7, CTCF, DNMT1, EZH2, HDAC1, HLTF, LSD1, MORC2, PCAF, PHF8, RBBP5, SAFB, SAFB2, SUZ12, WDR5 |
| RNA processing and export | FUS, hnRNPC, hnRNPH, hnRNPL, hnRNPUL1, ILF3, KHSRP, NONO, NUP98, PTBP1, RBFOX2, RBM15, RBM22, SRSF1, SRSF7, SRSF9, TARDBP, U2AF1, U2AF2, ZC3H11A |
| Transcription regulation | EWSR1, FUS, GTF2F1, hnRNPK, hnRNPUL1, LARP7, NONO, PCBP1, TAF15, U2AF2, YBX3, ZC3H8 |

**Supplementary Table S8.** Distribution of the of cis- and trans-contacts abundances in constructed triads for each protein. Unless stated otherwise, triads are based on eCLIP RNA-protein interactions data, Red-C RNA-DNA interactions data for K562 cells. Ratios of mRNA-mediated triads are also shown.

| Protein | Cis-contacts number | Trans-contacts number | mRNA contacts ratio |
|---|---|---|---|
| CBP fRIP | 26 | 497 | 0.4608 |
| CBX3 fRIP | 136 | 745 | 0.4721 |
| CHD1 fRIP | 856 | 10396 | 0.4878 |
| CHD4 fRIP | 200 | 2953 | 0.5839 |
| CHD7 fRIP | 99 | 782 | 0.6139 |
| CTCF fRIP | 2368 | 29571 | 0.4818 |
| DNMT1 fRIP | 52 | 715 | 0.6698 |
| EWSR1 | 157 | 1045 | 0.3135 |
| EZH2 GRID-Seq mESC | 664 | 3480 | 0.0443 |
| EZH2 RADICL-Seq 2FA mESC | 2965 | 21016 | 0.1435 |
| EZH2 RADICL-Seq NPM mESC | 2976 | 35917 | 0.0799 |

See next page

**Supplementary Table S8** – continued from previous page

| Protein | Cis-contacts number | Trans-contacts number | mRNA contacts ratio |
|---|---|---|---|
| EZH2 fRIP | 44 | 710 | 0.6370 |
| FUS | 124 | 95 | 0.6451 |
| GTF2F1 | 238 | 2173 | 0.3867 |
| HDAC1 fRIP | 522 | 5490 | 0.5099 |
| HLTF | 46 | 1220 | 0.3788 |
| HNRNPC | 10 | 17 | 0.4490 |
| HNRNPH fRIP | 22 | 36 | 0.5538 |
| HNRNPK GRID-Seq mESC | 16 | 153 | 0.1592 |
| HNRNPK RADICL-Seq 2FA mESC | 162 | 1423 | 0.1587 |
| HNRNPK RADICL-Seq NPM mESC | 76 | 2879 | 0.0817 |
| HNRNPK | 631 | 7096 | 0.3692 |
| HNRNPL | 57 | 323 | 0.4501 |
| HNRNPUL1 | 243 | 1451 | 0.2754 |
| ILF3 | 23 | 562 | 0.1667 |
| KHSRP | 28 | 145 | 0.7882 |
| LARP7 | 725 | 7210 | 0.3677 |
| LSD1 fRIP | 1359 | 20011 | 0.4914 |
| MORC2 | 98 | 1802 | 0.1453 |
| NONO | 239 | 1974 | 0.5931 |
| NUP98 fRIP | 290 | 4153 | 0.4960 |
| PCAF fRIP | 29 | 321 | 0.6881 |
| PCBP1 | 864 | 9524 | 0.4099 |
| PHF8 fRIP | 1931 | 20136 | 0.4601 |
| PTBP1 | 497 | 4780 | 0.3508 |

See next page

| Protein | Cis-contacts number | Trans-contacts number | mRNA contacts ratio |
|---|---|---|---|
| RBBP5 fRIP | 1362 | 19515 | 0.4110 |
| RBFOX2 | 257 | 1169 | 0.5750 |
| RBM15 | 15 | 229 | 0.4106 |
| RBM22 | 772 | 10296 | 0.3857 |
| SAFB2 | 14 | 36 | 0.1935 |
| SAFB | 122 | 480 | 0.3348 |
| SRSF1 | 25 | 185 | 0.6111 |
| SRSF7 | 4 | 81 | 0.5874 |
| SRSF9 | 5 | 21 | 0.0370 |
| SUZ12 GRID-Seq mESC | 3899 | 56261 | 0.0987 |
| SUZ12 fRIP GRID-Seq mESC | 3012 | 47458 | 0.0917 |
| SUZ12 RADICL-Seq 2FA mESC | 17403 | 314152 | 0.1865 |
| SUZ12 fRIP RADICL-Seq 2FA mESC | 14355 | 254532 | 0.2487 |
| SUZ12 RADICL-Seq NPM mESC | 28444 | 608381 | 0.0453 |
| SUZ12 fRIP RADICL-Seq NPM mESC | 10813 | 198167 | 0.1739 |
| SUZ12 fRIP | 29 | 194 | 0.7525 |
| TAF15 | 76 | 359 | 0.4208 |
| TARDBP | 580 | 9031 | 0.1606 |
| U2AF1 | 343 | 3644 | 0.3795 |
| U2AF2 | 11 | 26 | 0.5698 |
| WDR5 fRIP GRID-Seq mESC | 1 | 5 | 0.8182 |

See next page

**Supplementary Table S8** – continued from previous page

| Protein | Cis-contacts number | Trans-contacts number | mRNA contacts ratio |
|---|---|---|---|
| WDR5 fRIP RADICL-Seq 2FA mESC | 4 | 419 | 0.5490 |
| WDR5 fRIP RADICL-Seq NPM mESC | 1 | 497 | 0.3656 |
| WDR5 fRIP | 733 | 9944 | 0.4998 |
| YBX3 | 2 | 38 | 0.8326 |
| ZC3H11A | 82 | 637 | 0.4524 |
| ZC3H8 | 877 | 9851 | 0.3657 |

**Supplementary Table S9.** cis-/trans- contacts ratio statistics for selected RNAs for K562 Red-C experiment.

| RNA | RNA Type | Red-C cis-/trans-contacts ratio | Triads cis-/trans-contacts ratio |
|---|---|---|---|
| 1840_HUVEC | vlinc | 0.193256 | 0.130892 |
| 2125_K562 | vlinc | 0.177960 | 0.120279 |
| 2286_K562 | vlinc | 0.213551 | 0.170541 |
| AC005208.1 | lncRNA | 0.168325 | 0.136384 |
| AC012494.1 | lncRNA | 0.182849 | 0.064323 |
| AC020687.1 | lncRNA | 0.290336 | 0.163065 |
| AC097528.1 | lncRNA | 0.194500 | 0.031450 |
| AL390334.1 | pseudogene | 0.252462 | 0.090108 |
| AL445255.1 | lncRNA | 0.271939 | 0.073506 |
| AL713998.1 | lncRNA | 0.074473 | 0.078996 |
| B3GALT1-AS1 | lncRNA | 0.195343 | 0.098824 |
| C22orf34 | lncRNA | 0.421999 | 0.474465 |
| CCDC26 | lncRNA | 0.217844 | 0.394440 |
| DSCR4 | lncRNA | 0.512160 | 0.730741 |
| FTX | lncRNA | 0.085945 | 0.089114 |

See next page

**Supplementary Table S9** – continued from previous page

| RNA | RNA Type | Red-C cis-/trans-contacts ratio | Triads cis-/trans-contacts ratio |
|---|---|---|---|
| KCNQ1OT1 | lncRNA | 0.293347 | 0.352229 |
| LINC00355 | lncRNA | 0.273391 | 0.023474 |
| LINC00470 | lncRNA | 0.231691 | 0.160800 |
| LINC02476 | lncRNA | 0.243996 | 0.458085 |
| MIR4435-2HG | lncRNA | 0.214966 | 0.153130 |
| MIR548XHG | lncRNA | 0.431875 | 0.222322 |
| RMRP_2 | ribozyme | 0.037462 | 0.033039 |
| RPPH1 | ribozyme | 0.047262 | 0.020858 |
| U2_728 | snRNA | 0.055386 | 0.057798 |
| X_13_10942_a_hg38 | Xrna | 0.036283 | 0.011467 |
| X_2_7832_a_hg38 | Xrna | 0.198941 | 0.124180 |
| X_4_12686_b_hg38 | Xrna | 0.205287 | 0.082588 |

**Supplementary Table S10.** Triads and ChIP-Seq composition by ChromHMM annotation of DNA loci for all proteins split by function. *cnt* – number of observed ChIP-Seq loci in a state; *tri* – number of triads; *Z* – z-score of $\frac{tri}{cnt}$ distribution over all ChromHMM states

| **State** | $\frac{cnt}{10^4}$ | % | **All proteins** | | | **Remodelling** | | **Tr.Reg** | | **RNA proc.** | |
|---|---|---|---|---|---|---|---|---|---|---|---|
| | | | $\frac{tri}{10^3}$ | % | Z | $\frac{tri}{10^3}$ | Z | $\frac{tri}{10^3}$ | Z | $\frac{tri}{10^3}$ | Z |
| Promoter | 56.51 | 23.58 | 848.70 | 34.69 | -0.22 | 806.37 | -0.23 | 24.10 | 0.38 | 18.23 | 0.54 |
| Enhancer | 108.77 | 45.39 | 455.95 | 18.63 | -0.64 | 398.61 | -0.63 | 32.86 | -0.35 | 24.48 | -0.24 |
| Transcribed | 40.59 | 16.94 | 191.08 | 7.81 | -0.62 | 158.96 | -0.62 | 14.47 | -0.03 | 17.64 | 1.45 |
| Insulator | 6.93 | 2.89 | 90.35 | 3.69 | -0.30 | 88.84 | -0.28 | 0.84 | -1.42 | 0.67 | -1.28 |
| PRC_Repressed | 7.95 | 3.32 | 582.11 | 23.79 | 2.01 | 578.35 | 2.01 | 2.59 | -0.21 | 1.17 | -0.87 |
| HetChrom_CNV | 18.90 | 7.89 | 278.60 | 11.39 | -0.23 | 260.74 | -0.24 | 12.08 | 1.63 | 5.78 | 0.41 |
| TOTAL | 239.66 | 100.00 | 2446.79 | 100.00 | -0.41 | 2291.87 | -0.41 | 86.94 | 0.01 | 67.98 | 0.23 |

**Supplementary Table S11.** Triads and RD-contacts DNA parts composition by ChromHMM annotation of DNA loci for all proteins split by function. *cnt* – number of observed DNA-parts in a state; *tri* – number of triads; *Z* – z-score of $\frac{tri}{cnt}$ distribution over all ChromHMM states

| **State** | $\frac{cnt}{10^6}$ | % | **All proteins** | | | **Remodelling** | | **Tr.Reg** | | **RNA proc.** | |
|---|---|---|---|---|---|---|---|---|---|---|---|
| | | | $\frac{tri}{10^3}$ | % | Z | $\frac{tri}{10^3}$ | Z | $\frac{tri}{10^3}$ | Z | $\frac{tri}{1000}$ | Z |
| Promoter | 6.36 | 2.78 | 848.70 | 34.69 | 1.98 | 806.37 | 1.97 | 24.10 | 1.78 | 18.23 | 1.76 |
| Enhancer | 16.48 | 7.20 | 455.95 | 18.63 | -0.23 | 398.61 | -0.27 | 32.86 | 0.56 | 24.48 | 0.51 |
| Transcribed | 20.31 | 8.88 | 191.08 | 7.81 | -0.61 | 158.96 | -0.63 | 14.47 | -0.31 | 17.64 | -0.05 |
| Insulator | 2.72 | 1.19 | 90.35 | 3.69 | -0.12 | 88.84 | -0.09 | 0.84 | -0.58 | 0.67 | -0.62 |
| PRC_Repressed | 21.64 | 9.46 | 582.11 | 23.79 | -0.25 | 578.35 | -0.22 | 2.59 | -0.71 | 1.17 | -0.79 |
| HetChrom_CNV | 161.30 | 70.49 | 278.60 | 11.39 | -0.77 | 260.74 | -0.77 | 12.08 | -0.74 | 5.78 | -0.81 |
| TOTAL | 228.81 | 100.00 | 2446.79 | 100.00 | -0.58 | 2291.87 | -0.58 | 86.94 | -0.54 | 67.98 | -0.57 |

**Supplementary Table S12.** Triads and ChIP-Seq composition by SPIN annotation of DNA loci for all proteins split by function. *cnt* – number of observed ChIP-Seq loci in a state; *tri* – number of triads; *Z* – z-score of $\frac{tri}{cnt}$ distribution over all SPIN states

| **State** | $\frac{cnt}{10^4}$ | % | **All proteins** | | | **Remodelling** | | **Tr.Reg** | | **RNA proc.** | |
|---|---|---|---|---|---|---|---|---|---|---|---|
| | | | $\frac{tri}{10^3}$ | % | Z | $\frac{tri}{10^3}$ | Z | $\frac{tri}{10^3}$ | Z | $\frac{tri}{10^3}$ | Z |
| A_compartment | 21.58 | 53.87 | 235.24 | 54.56 | 0.90 | 153.07 | 0.99 | 42.79 | 0.75 | 39.38 | 0.69 |
| B_compartment | 1.68 | 4.19 | 13.09 | 3.04 | -0.40 | 9.57 | -0.14 | 2.08 | -0.72 | 1.44 | -0.54 |
| Lamina | 0.62 | 1.56 | 3.76 | 0.87 | -1.14 | 2.53 | -1.45 | 0.96 | -0.13 | 0.27 | -1.09 |
| Lamina_like | 0.32 | 0.79 | 2.26 | 0.52 | -0.69 | 1.58 | -0.73 | 0.37 | -0.90 | 0.32 | -0.36 |
| Near_Lm | 4.73 | 11.81 | 39.60 | 9.18 | -0.16 | 28.85 | 0.18 | 6.06 | -0.64 | 4.70 | -0.37 |
| Speckle | 11.13 | 27.79 | 137.18 | 31.82 | 1.49 | 81.27 | 1.15 | 27.12 | 1.64 | 28.79 | 1.67 |
| TOTAL | 40.07 | 100.00 | 431.13 | 100.00 | 0.84 | 276.87 | 0.84 | 79.37 | 0.74 | 74.90 | 0.75 |

**Supplementary Table S13.** Triads and and RD-contacts DNA parts composition by SPIN annotation of DNA loci for all proteins split by function. *cnt* – number of of observed DNA-parts in a state; *tri* – number of triads; *Z* – z-score of $\frac{tri}{cnt}$ distribution over all SPIN states

| State | $\frac{cnt}{10^5}$ | % | All proteins | | | Remodelling | | Tr.Reg | | RNA proc. | |
|---|---|---|---|---|---|---|---|---|---|---|---|
| | | | $\frac{tri}{10^3}$ | % | Z | $\frac{tri}{10^3}$ | Z | $\frac{tri}{10^3}$ | Z | $\frac{tri}{10^3}$ | Z |
| A_compartment | 205.61 | 37.63 | 235.24 | 54.56 | 0.22 | 153.07 | 0.28 | 42.79 | 0.18 | 39.38 | 0.11 |
| B_compartment | 49.58 | 9.08 | 13.09 | 3.04 | -0.50 | 9.57 | -0.49 | 2.08 | -0.50 | 1.44 | -0.51 |
| Lamina | 100.68 | 18.43 | 3.76 | 0.87 | -0.68 | 2.53 | -0.72 | 0.96 | -0.63 | 0.27 | -0.61 |
| Lamina_like | 8.91 | 1.63 | 2.26 | 0.52 | -0.50 | 1.58 | -0.51 | 0.37 | -0.50 | 0.32 | -0.48 |
| Near_Lm | 139.20 | 25.48 | 39.60 | 9.18 | -0.48 | 28.85 | -0.47 | 6.06 | -0.49 | 4.70 | -0.49 |
| Speckle | 42.33 | 7.75 | 137.18 | 31.82 | 1.94 | 81.27 | 1.92 | 27.12 | 1.95 | 28.79 | 1.97 |
| TOTAL | 546.32 | 100.00 | 431.13 | 100.00 | -0.07 | 276.87 | -0.05 | 79.37 | -0.08 | 74.90 | -0.10 |

**Supplementary Table S14.** Triads and and RD-contacts composition by RNA biotypes for all proteins split by function. *cnt* – number of of observed RNA of a given biotype; *tri* – number of triads; *Z* – z-score of $\frac{tri}{cnt}$ distribution over all RNA biotypes

| Biotypes | $\frac{cnt}{10^4}$ | % | All proteins | | | Remodelling | | Tr.Reg | | RNA proc. | |
|---|---|---|---|---|---|---|---|---|---|---|---|
| | | | $\frac{tri}{10^3}$ | % | Z | $\frac{tri}{10^3}$ | Z | $\frac{tri}{10^3}$ | Z | $\frac{tri}{10^3}$ | Z |
| mRNA | 87.14 | 66.56 | 471.91 | 19.61 | -0.45 | 409.31 | -0.44 | 32.69 | -0.60 | 29.90 | -0.49 |
| rRNA | 0.23 | 0.18 | 853.58 | 35.47 | 2.62 | 850.18 | 2.62 | 3.19 | 0.44 | 0.21 | -0.45 |
| snRNA | 0.39 | 0.30 | 216.48 | 8.99 | 0.00 | 186.63 | -0.05 | 15.25 | 2.39 | 14.60 | 2.50 |
| snoRNA | 0.38 | 0.29 | 217.21 | 9.03 | 0.02 | 206.78 | 0.01 | 5.25 | 0.44 | 5.19 | 0.58 |
| lncRNA | 22.67 | 17.32 | 113.43 | 4.71 | -0.45 | 96.34 | -0.44 | 9.13 | -0.60 | 7.97 | -0.49 |
| X-RNA | 11.52 | 8.80 | 431.17 | 17.92 | -0.42 | 414.85 | -0.41 | 8.57 | -0.57 | 7.75 | -0.46 |
| miRNA | 0.76 | 0.58 | 20.27 | 0.84 | -0.43 | 20.23 | -0.42 | 0.04 | -0.63 | 0.00 | -0.52 |
| pseudo | 6.08 | 4.65 | 20.16 | 0.84 | -0.45 | 11.78 | -0.44 | 2.81 | -0.60 | 2.42 | -0.48 |
| other | 1.74 | 1.33 | 62.49 | 2.60 | -0.43 | 44.41 | -0.42 | 8.08 | -0.27 | 6.87 | -0.19 |
| TOTAL | 130.92 | 100.00 | 2406.70 | 100.00 | -0.44 | 2240.51 | -0.43 | 85.01 | -0.58 | 74.90 | -0.47 |

**Supplementary Table S15.** *p*-values for the real number of triads on the background distribution of the number of triads obtained on simulated data. The background distribution for triads number was fit using a gamma distribution, and the *p*-value defined as the probability of obtaining the real number of triads for protein or a value greater from the fit distribution.

| Triads constructed on | *p*-value of simulation | adjusted *p*-value |
|---|---|---|
| CBP Red-C fRIP | 141.331 | 140.84 |
| CBX3 Red-C fRIP | 102.742 | 102.412 |
| CHD1 Red-C fRIP | 102.54 | 102.224 |
| CHD4 Red-C fRIP | 162.695 | 162.181 |

See next page

**Supplementary Table S15** – continued from previous page

| Triads constructed on | *p*-value of simulation | adjusted *p*-value |
|---|---|---|
| CHD7 Red-C fRIP | 125.304 | 124.854 |
| CTCF Red-C fRIP | 180.472 | 179.91 |
| DNMT1 Red-C fRIP | 18.547 | 18.426 |
| EWSR1 Red-C eCLIP | 104.636 | 104.291 |
| EZH2 GRID eCLIP | 296.277 | 295.631 |
| EZH2 RADICL 2FA eCLIP | 296.277 | 295.631 |
| EZH2 RADICL NPM eCLIP | 296.277 | 295.631 |
| EZH2 Red-C fRIP | 63.279 | 63.054 |
| FUS Red-C eCLIP | 3.27 | 3.248 |
| GTF2F1 Red-C eCLIP | 115.802 | 115.407 |
| HDAC1 Red-C fRIP | 120.077 | 119.647 |
| HLTF Red-C eCLIP | 177.046 | 176.509 |
| HNRNPC Red-C eCLIP | 8.816 | 8.756 |
| HNRNPH Red-C fRIP | 7.009 | 6.964 |
| HNRNPK GRID eCLIP | 52.601 | 52.421 |
| HNRNPK RADICL2FA eCLIP | 296.277 | 295.631 |
| HNRNPK RADICL NPM eCLIP | 296.277 | 295.631 |
| HNRNPK Red-C eCLIP | 54.8 | 54.61 |
| HNRNPL Red-C eCLIP | 2.346 | 2.332 |
| HNRNPUL1 Red-C eCLIP | 110.168 | 109.807 |
| ILF3 Red-C eCLIP | 33.853 | 33.704 |
| KHSRP Red-C eCLIP | 3.504 | 3.475 |
| LARP7 Red-C eCLIP | 119.872 | 119.46 |
| LSD1 Red-C fRIP | 8.851 | 8.783 |
| MORC2 Red-C eCLIP | 66.157 | 65.896 |
| NONO Red-C eCLIP | 27.661 | 27.521 |

See next page

**Supplementary Table S15** – continued from previous page

| Triads constructed on | *p*-value of simulation | adjusted *p*-value |
|---|---|---|
| NUP98 Red-C fRIP | 296.277 | 295.631 |
| PCAF Red-C fRIP | 62.037 | 61.824 |
| PCBP1 Red-C eCLIP | 111.221 | 110.844 |
| PHF8 Red-C fRIP | 296.277 | 295.631 |
| PTBP1 Red-C eCLIP | 296.277 | 295.631 |
| RBBP5 Red-C fRIP | 296.277 | 295.631 |
| RBFOX2 Red-C eCLIP | 56.393 | 56.192 |
| RBM15 Red-C eCLIP | 40.249 | 40.09 |
| RBM22 Red-C eCLIP | 90.91 | 90.636 |
| SAFB2 Red-C eCLIP | 8.024 | 7.972 |
| SAFB Red-C eCLIP | 12.453 | 12.36 |
| SRSF1 Red-C eCLIP | 12.768 | 12.666 |
| SRSF7 Red-C eCLIP | 19.857 | 19.727 |
| SRSF9 Red-C eCLIP | 9.63 | 9.545 |
| SUZ12 GRID eCLIP | 296.277 | 295.631 |
| SUZ12 GRID fRIP | 296.277 | 295.631 |
| SUZ12 RADICL 2FA eCLIP | 296.277 | 295.631 |
| SUZ12 RADICL 2FA fRIP | 296.277 | 295.631 |
| SUZ12 RADICL NPM eCLIP | 99.435 | 99.134 |
| SUZ12 RADICL NPM fRIP | 192.606 | 191.989 |
| SUZ12 Red-C fRIP | 3.823 | 3.787 |
| TAF15 Red-C eCLIP | 65.584 | 65.336 |
| TARDBP Red-C eCLIP | 296.277 | 295.631 |
| U2AF1 Red-C eCLIP | 98.987 | 98.7 |
| U2AF2 Red-C eCLIP | 9.369 | 9.292 |
| WDR5 GRID fRIP | 1.653 | 1.653 |

See next page

**Supplementary Table S15** – continued from previous page

| Triads constructed on | *p*-value of simulation | adjusted *p*-value |
|---|---|---|
| WDR5 RADICL 2FA fRIP | 52.543 | 52.374 |
| WDR5 RADICL NPM fRIP | 64.79 | 64.554 |
| WDR5 Red-C fRIP | 14.914 | 14.802 |
| YBX3 Red-C eCLIP | 1.811 | 1.804 |
| ZC3H11A Red-C eCLIP | 126.45 | 125.98 |
| ZC3H8 Red-C eCLIP | 191.491 | 190.903 |

**Supplementary Table S16.** RNA orthologs found by ortho2align (selected as bidirectional best hit)

| Human RNA | Mouse RNA | In triads with | Human biotype | Mouse biotype |
|---|---|---|---|---|
| U79 | Gas5 | EZH2 | CDBox | processed transcript |
| | Gm50452 | EZH2 | CDBox | snoRNA |
| RPPH1 | Rpph1 | EZH2, hnRNPK | ribozyme | ribozyme |
| SNORD58B | Gm23301 | EZH2, hnRNPK | snoRNA | snoRNA |
| RNU4-2 | Gm24265 | EZH2, SUZ12 | snRNA | snRNA |
| C19orf48 | Gm26247 | EZH2 | pseudogene | snoRNA |
| | Snord88a | EZH2 | pseudogene | snoRNA |
| | Gm23991 | EZH2 | pseudogene | snoRNA |
| DLEU2 | Dleu2 | hnRNPK | lncRNA | processed transcript |
| RNF213-AS1 | A930037H05Rik | SUZ12 | lncRNA | processed transcript |
| SNORD5 | Gm24299 | SUZ12 | snoRNA | snoRNA |

**Supplementary Table S17.** DNA contacts distances in original genomes for RNA orthologs. *"RNA from source genome"* stands for RNA, for which coordinates of DNA locus that it interacts with were unchanged. *"RNA for liftOver"* stands for RNA, for which coordinates of DNA locus were lifted over to the opposing genome.

| DNA Chr | RNA from source genome | RNA for liftOver | Distance, nt | Triads formed by |
|---|---|---|---|---|
| chr16 | Gm24299 | SNORD5 | 71101 | SUZ12 |
| chr22 | SNORD5 | Gm24299 | 79851 | SUZ12 |

See next page

**Supplementary Table S17** – continued from previous page

| DNA Chr | RNA from source genome | RNA for liftOver | Distance, nt | Triads formed by |
|---|---|---|---|---|
| chr16 | Gm24265 | RNU4-2 | 114378 | EZH2 |
| chr11 | Rpph1 | RPPH1 | 334071 | HNRNPK |
| chr16 | Gm24265 | RNU4-2 | 409387 | SUZ12 |
| chr11 | Gm24265 | RNU4-2 | 627098 | SUZ12 |
| chr2 | Rpph1 | RPPH1 | 1529017 | EZH2 |
| chr10 | Gm24299 | SNORD5 | 1575789 | SUZ12 |
| chr12 | SNORD5 | Gm24299 | 1722366 | SUZ12 |
| chr19 | RNU4-2 | Gm24265 | 1920768 | SUZ12 |
| chr11 | Rpph1 | RPPH1 | 2543554 | EZH2 |
| chr7 | RNU4-2 | Gm24265 | 2644315 | SUZ12 |
| chr6 | Gm24265 | RNU4-2 | 2704283 | SUZ12 |
| chr12 | Gm26247 | C19orf48 | 4866317 | EZH2 |
| chr17 | Gm50452 | U79 | 5350920 | EZH2 |
| chr15 | Rpph1 | RPPH1 | 5502957 | EZH2 |
| chr14 | C19orf48 | Gm26247 | 6136940 | EZH2 |
| chr17 | RNU4-2 | Gm24265 | 6459825 | SUZ12 |
| chr6 | U79 | Gm50452 | 6513447 | EZH2 |
| chr12 | RPPH1 | Rpph1 | 6675254 | EZH2 |
| chr2 | Gas5 | U79 | 7172769 | EZH2 |
| chr6 | RPPH1 | Rpph1 | 7203675 | EZH2 |
| chr3 | Rpph1 | RPPH1 | 7558544 | EZH2 |
| chr15 | Gm24265 | RNU4-2 | 7737350 | SUZ12 |
| chr7 | RNU4-2 | Gm24265 | 8161423 | SUZ12 |
| chr2 | U79 | Gas5 | 8529496 | EZH2 |
| chr22 | RPPH1 | Rpph1 | 9440164 | EZH2 |
| chr2 | U79 | Gas5 | 9528528 | EZH2 |

See next page

| DNA Chr | RNA from source genome | RNA for liftOver | Distance, nt | Triads formed by |
|---|---|---|---|---|
| chr10 | Rpph1 | RPPH1 | 9815790 | EZH2 |
| chr17 | Rpph1 | RPPH1 | 10880342 | EZH2 |
| chr5 | RPPH1 | Rpph1 | 11601727 | EZH2 |
| chr2 | Rpph1 | RPPH1 | 11693085 | EZH2 |
| chr10 | RPPH1 | Rpph1 | 12013038 | EZH2 |
| chr2 | RPPH1 | Rpph1 | 12542982 | EZH2 |
| chr16 | Gm24265 | RNU4-2 | 12618598 | SUZ12 |
| chr5 | Rpph1 | RPPH1 | 13398672 | EZH2 |
| chr11 | Gm23301 | SNORD58B | 13574192 | HNRNPK |
| chr11 | Gm24265 | RNU4-2 | 17752338 | SUZ12 |
| chr16 | Gm50452 | U79 | 18906235 | EZH2 |
| chr12 | RPPH1 | Rpph1 | 19771353 | EZH2 |
| chr22 | RNU4-2 | Gm24265 | 20303830 | SUZ12 |
| chr7 | SNORD58B | Gm23301 | 20517250 | EZH2 |
| chr8 | Gm24265 | RNU4-2 | 20957564 | SUZ12 |
| chr7 | RNU4-2 | Gm24265 | 21869558 | SUZ12 |
| chr11 | Gm23301 | SNORD58B | 22013524 | EZH2 |
| chr3 | RPPH1 | Rpph1 | 23617849 | EZH2 |
| chr10 | U79 | Gm50452 | 24060208 | EZH2 |
| chr1 | Gm23301 | SNORD58B | 26949946 | EZH2 |
| chr8 | Rpph1 | RPPH1 | 28768814 | EZH2 |
| chr17 | SNORD58B | Gm23301 | 30342819 | EZH2 |
| chr2 | Gas5 | U79 | 33594729 | EZH2 |
| chr14 | Gm24265 | RNU4-2 | 34242552 | SUZ12 |
| chr18 | Gm23301 | SNORD58B | 34937500 | EZH2 |
| chr11 | RPPH1 | Rpph1 | 35518520 | EZH2 |

See next page

**Supplementary Table S17** – continued from previous page

| DNA Chr | RNA from source genome | RNA for liftOver | Distance, nt | Triads formed by |
|---|---|---|---|---|
| chr17 | Gm24265 | RNU4-2 | 39573176 | SUZ12 |
| chr11 | Rpph1 | RPPH1 | 40362990 | EZH2 |
| chr7 | Rpph1 | RPPH1 | 41092890 | EZH2 |
| chr9 | Gm50452 | U79 | 44981079 | EZH2 |
| chr6 | Gm24299 | SNORD5 | 45606058 | SUZ12 |
| chr4 | Gm50452 | U79 | 46787576 | EZH2 |
| chr7 | SNORD5 | Gm24299 | 51881334 | SUZ12 |
| chr2 | Rpph1 | RPPH1 | 52531323 | EZH2 |
| chr3 | Rpph1 | RPPH1 | 55440322 | EZH2 |
| chr16 | Gm24265 | RNU4-2 | 58216738 | SUZ12 |
| chr2 | Rpph1 | RPPH1 | 60112309 | EZH2 |
| chr20 | U79 | Gas5 | 62122460 | EZH2 |
| chr16 | RNU4-2 | Gm24265 | 66900238 | SUZ12 |
| chr17 | Rpph1 | RPPH1 | 70320379 | EZH2 |
| chr6 | RNU4-2 | Gm24265 | 74855836 | SUZ12 |
| chr5 | Gm50452 | U79 | 88642071 | EZH2 |
| chr1 | Gas5 | U79 | 103951727 | EZH2 |
| chr5 | Gm50452 | U79 | 114030242 | EZH2 |
| chr1 | Rpph1 | RPPH1 | 114229243 | EZH2 |
| chr7 | U79 | Gm50452 | 131126867 | EZH2 |

**Supplementary Table S18.** Results for the association tests: $\chi^2$- goodness-of-fit of ChromHMM annotation of DNA-parts and Fisher's exact test for association of PR- and PD-contacts data of protein with RD-contacts.

| Protein | q-value($\chi^2$) | q-value(Fisher) |
|---|---|---|
| CBP | $5.10 * 10^{-18}$ | $7.86 * 10^{-6}$ |
| CBX3 | $2.36 * 10^{-61}$ | $7.41 * 10^{-5}$ |
| CHD1 | $3.14 * 10^{-214}$ | $1.25 * 10^{-1}$ |

See next page

**Supplementary Table S18** – continued from previous page

| **Protein** | **q-value($\chi^2$)** | **q-value(Fisher)** |
|---|---|---|
| CHD4 | $8.27 * 10^{-57}$ | $1.01 * 10^{-3}$ |
| CHD7 | $3.68 * 10^{-36}$ | $3.61 * 10^{-4}$ |
| CTCF | 0 | $4.45 * 10^{-1}$ |
| DNMT1 | $1.50 * 10^{-6}$ | $9.62 * 10^{-5}$ |
| EWSR1 | $3.68 * 10^{-36}$ | $9.62 * 10^{-5}$ |
| EZH2 | $5.30 * 10^{-40}$ | 0.001060326675 |
| FUS | $1.37 * 10^{-31}$ | $7.87 * 10^{-4}$ |
| GTF2F1 | 0 | $3.14 * 10^{-4}$ |
| HDAC1 | $1.56 * 10^{-97}$ | 0.01977 |
| HLTF | $8.24 * 10^{-19}$ | 0.000173 |
| HNRNPC | $7.74 * 10^{-5}$ | $3.59 * 10^{-2}$ |
| HNRNPH | $5.87 * 10^{-6}$ | $1.73 * 10^{-8}$ |
| HNRNPK | $3.23 * 10^{-303}$ | $6.74 * 10^{-2}$ |
| HNRNPL | $4.09 * 10^{-9}$ | 0.0001024 |
| HNRNPUL1 | $7.20 * 10^{-94}$ | $9.62 * 10^{-5}$ |
| ILF3 | $7.37 * 10^{-5}$ | 0.003031 |
| KHSRP | 0.78792 | 0.0004923 |
| LARP7 | $1.30 * 10^{-118}$ | $9.46 * 10^{-2}$ |
| LSD1 | 0 | 0.4440061483 |
| MORC2 | $1.33 * 10^{-105}$ | $5.50 * 10^{-3}$ |
| NONO | $2.65 * 10^{-9}$ | $7.87 * 10^{-4}$ |
| NUP98 | $2.01 * 10^{-86}$ | 0.016093 |
| PCAF | $5.71 * 10^{-14}$ | 0.0007874 |
| PCBP1 | $6.08 * 10^{-115}$ | $9.50 * 10^{-2}$ |
| PHF8 | 0 | 0.36820 |
| PTBP1 | $1.28 * 10^{-91}$ | 0.01575443137 |

See next page

**Supplementary Table S18** – continued from previous page

| Protein | q-value($\chi^2$) | q-value(Fisher) |
|---|---|---|
| RBBP5 | $1.11 * 10^{-188}$ | $3.68 * 10^{-1}$ |
| RBFOX2 | $1.95 * 10^{-60}$ | 0.001525 |
| RBM15 | 0.000103582 | 0.0004245 |
| RBM22 | 0 | $1.25 * 10^{-1}$ |
| SAFB | $2.65 * 10^{-31}$ | $5.02 * 10^{-4}$ |
| SAFB2 | 0.0002162 | $4.78 * 10^{-4}$ |
| SRSF1 | 0.071047 | $1.24 * 10^{-6}$ |
| SRSF7 | 0.210818 | $1.76 * 10^{-5}$ |
| SRSF9 | 0.039051 | 1.00 |
| SUZ12 | $7.62 * 10^{-15}$ | $3.99 * 10^{-4}$ |
| TAF15 | $1.49 * 10^{-41}$ | $7.86 * 10^{-6}$ |
| TARDBP | $2.41 * 10^{-93}$ | $9.50 * 10^{-2}$ |
| U2AF1 | $2.55 * 10^{-48}$ | 0.011595 |
| U2AF2 | 0.000608817 | $1.73 * 10^{-8}$ |
| WDR5 | $9.06 * 10^{-221}$ | 0.21304 |
| YBX3 | 0.68586 | 0.0016564 |
| ZC3H11A | $1.47 * 10^{-63}$ | $3.03 * 10^{-5}$ |
| ZC3H8 | $1.44 * 10^{-98}$ | $1.25 * 10^{-1}$ |

**Supplementary Table S19.** Results for the association test: $\chi^2$- goodness-of-fit of SPIN annotation of DNA-parts of triads and proteins ChIP-Seq loci.

| Protein | p-value($\chi^2$) | q-value($\chi^2$) |
|---|---|---|
| CBP | $2.51 * 10^{-34}$ | $6.95 * 10^{-34}$ |
| CBX3 | $1.06 * 10^{-22}$ | $2.07 * 10^{-22}$ |
| CHD1 | $1.72 * 10^{-84}$ | $1.01 * 10^{-83}$ |
| CHD4 | $1.00 * 10^{-20}$ | $1.89 * 10^{-20}$ |
| CHD7 | $6.01 * 10^{-32}$ | $1.57 * 10^{-31}$ |

See next page

**Supplementary Table S19** – continued from previous page

| Protein | p-value($\chi^2$) | q-value($\chi^2$) |
|---|---|---|
| CTCF | 0 | 0 |
| DNMT1 | $4.02 * 10^{-14}$ | $6.99 * 10^{-14}$ |
| EWSR1 | $2.26 * 10^{-74}$ | $1.06 * 10^{-73}$ |
| EZH2 | $1.13 * 10^{-25}$ | $2.52 * 10^{-25}$ |
| FUS | $2.62 * 10^{-6}$ | $3.84 * 10^{-6}$ |
| GTF2F1 | $1.46 * 10^{-27}$ | $3.43 * 10^{-27}$ |
| HDAC1 | $1.29 * 10^{-28}$ | $3.19 * 10^{-28}$ |
| HLTF | $1.63 * 10^{-5}$ | $2.25 * 10^{-5}$ |
| HNRNPC | $2.93 * 10^{-2}$ | $3.36 * 10^{-2}$ |
| HNRNPH | $3.80 * 10^{-3}$ | $4.71 * 10^{-3}$ |
| HNRNPK | $1.23 * 10^{-240}$ | $2.90 * 10^{-239}$ |
| HNRNPL | $1.52 * 10^{-1}$ | $1.59 * 10^{-1}$ |
| HNRNPUL1 | $1.03 * 10^{-18}$ | $1.87 * 10^{-18}$ |
| ILF3 | $7.11 * 10^{-2}$ | $7.77 * 10^{-2}$ |
| KHSRP | $1.43 * 10^{-2}$ | $1.68 * 10^{-2}$ |
| LARP7 | $1.33 * 10^{-69}$ | $5.20 * 10^{-69}$ |
| LSD1 | $3.12 * 10^{-122}$ | $2.10 * 10^{-121}$ |
| MORC2 | $3.04 * 10^{-62}$ | $1.10 * 10^{-61}$ |
| NONO | $2.65 * 10^{-54}$ | $8.31 * 10^{-54}$ |
| NUP98 | $1.49 * 10^{-81}$ | $7.78 * 10^{-81}$ |
| PCAF | $1.76 * 10^{-7}$ | $2.66 * 10^{-7}$ |
| PCBP1 | $8.67 * 10^{-186}$ | $1.02 * 10^{-184}$ |
| PHF8 | $1.12 * 10^{-153}$ | $8.80 * 10^{-153}$ |
| PTBP1 | $2.21 * 10^{-23}$ | $4.52 * 10^{-23}$ |
| RBBP5 | $1.43 * 10^{-185}$ | $1.35 * 10^{-184}$ |
| RBFOX2 | $1.93 * 10^{-13}$ | $3.23 * 10^{-13}$ |

See next page

**Supplementary Table S19** – continued from previous page

| **Protein** | **p-value($\chi^2$)** | **q-value($\chi^2$)** |
|---|---|---|
| RBM15 | $5.92 * 10^{-10}$ | $9.28 * 10^{-10}$ |
| RBM22 | $3.31 * 10^{-11}$ | $5.36 * 10^{-11}$ |
| SAFB | $5.37 * 10^{-3}$ | $6.47 * 10^{-3}$ |
| SAFB2 | $8.39 * 10^{-2}$ | $8.96 * 10^{-2}$ |
| SRSF1 | $8.30 * 10^{-5}$ | $1.05 * 10^{-4}$ |
| SRSF7 | $2.93 * 10^{-1}$ | $3.00 * 10^{-1}$ |
| SRSF9 | $3.90 * 10^{-5}$ | $5.09 * 10^{-5}$ |
| SUZ12 | $1.64 * 10^{-24}$ | $3.50 * 10^{-24}$ |
| TAF15 | $3.05 * 10^{-1}$ | $3.05 * 10^{-1}$ |
| TARDBP | $1.66 * 10^{-54}$ | $5.59 * 10^{-54}$ |
| U2AF1 | $3.09 * 10^{-48}$ | $9.08 * 10^{-48}$ |
| U2AF2 | $2.18 * 10^{-5}$ | $2.92 * 10^{-5}$ |
| WDR5 | $1.99 * 10^{-210}$ | $3.11 * 10^{-209}$ |
| YBX3 | $4.18 * 10^{-2}$ | $4.67 * 10^{-2}$ |
| ZC3H11A | $1.59 * 10^{-5}$ | $2.25 * 10^{-5}$ |
| ZC3H8 | $6.35 * 10^{-70}$ | $2.71 * 10^{-69}$ |

CBP Red-C fRIP-Seq K562

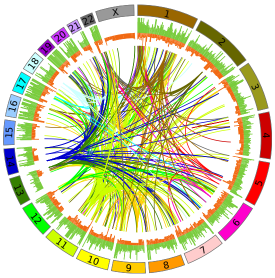

CBX3 Red-C fRIP-Seq K562

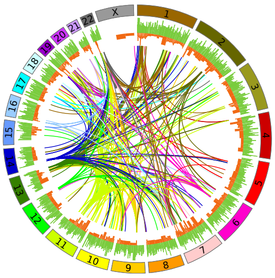

CHD1 Red-C fRIP-Seq K562

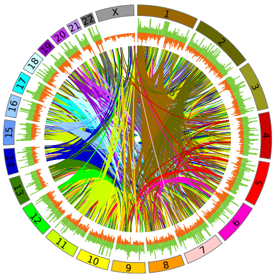

CHD4 Red-C fRIP-Seq K562

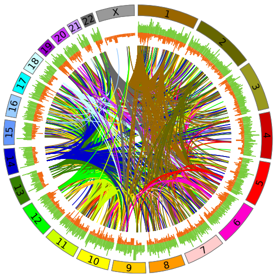

CHD7 Red-C fRIP-Seq K562

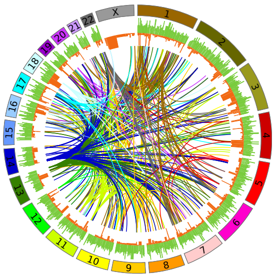

CTCF Red-C fRIP-Seq K562

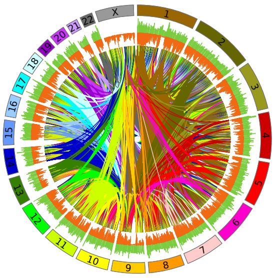

DNMT1 Red-C fRIP-Seq K562

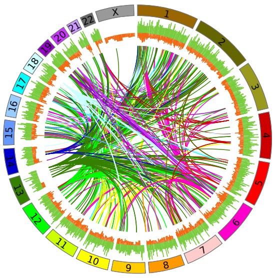

EWSR1 Red-C eCLIP-Seq K562

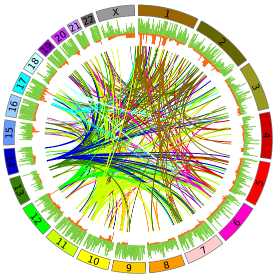

EZH2 GRID-Seq eCLIP-Seq mESC

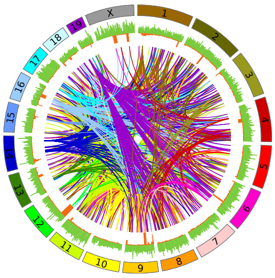

EZH2 RADICL-Seq 2FA eCLIP-Seq mESC

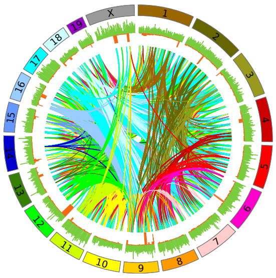

EZH2 RADICL-Seq NPM eCLIP-Seq mESC

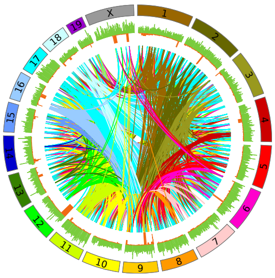

EZH Red-C fRIP-Seq K562

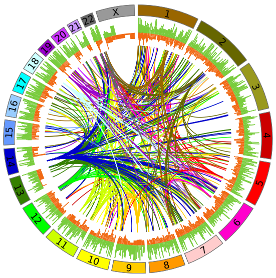

FUS Red-C eCLIP-Seq K562

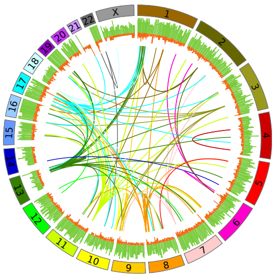

GTF2F1 Red-C eCLIP-Seq K562

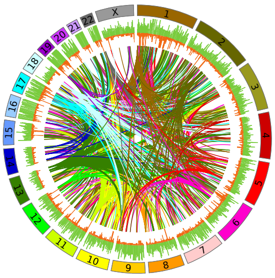

HDAC1 Red-C fRIP-Seq K562

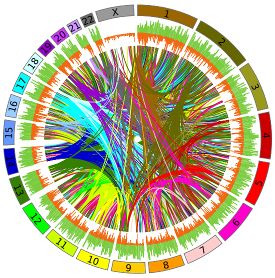

HLTF Red-C eCLIP-Seq K562

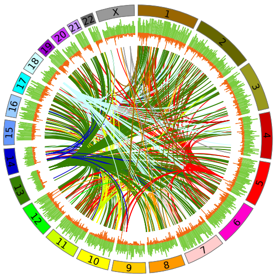

hnRNPC Red-C
eCLIP-Seq K562

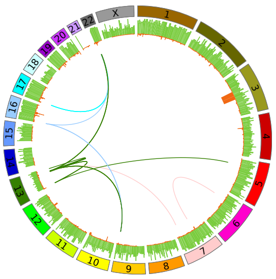

hnRNPH Red-C
fRIP-Seq K562

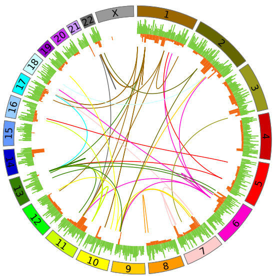

hnRNPK GRID-Seq
eCLIP-Seq mESC

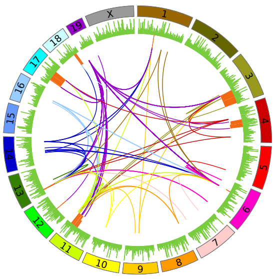

hnRNPK
RADICL-Seq 2FA
eCLIP-Seq mESC

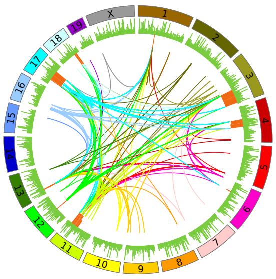

hnRNPK
RADICL-Seq NPM
eCLIP-Seq K562

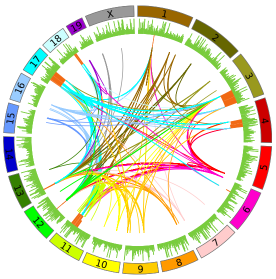

hnRNPK Red-C
eCLIP-Seq

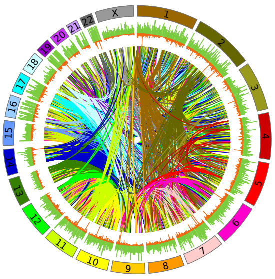

hnRNPL Red-C
eCLIP-Seq K562

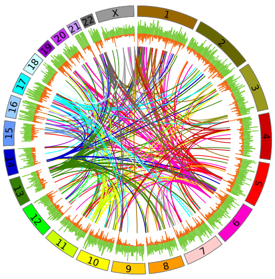

hnRNPUL1 Red-C
eCLIP-Seq K562

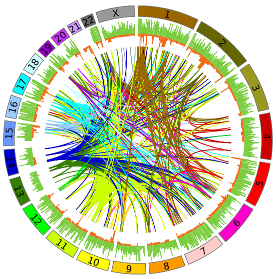

ILF3 Red-C
eCLIP-Seq K562

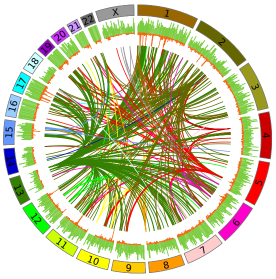

KHSRP Red-C
eCLIP-Seq K562

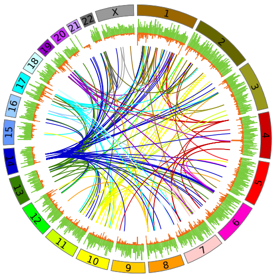

LARP7 Red-C
eCLIP-Seq K562

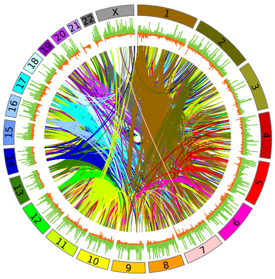

LSD1 Red-C fRIP-Seq
K562

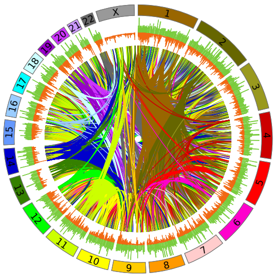

MORC2 Red-C
eCLIP-Seq K562

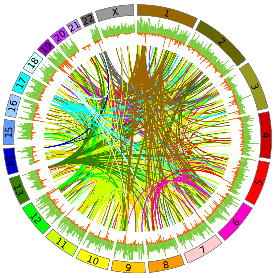

NONO Red-C
eCLIP-Seq K562

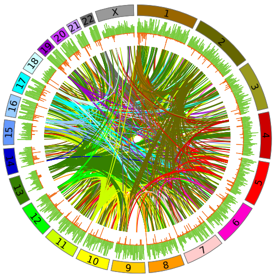

NUP98 Red-C
fRIP-Seq K562

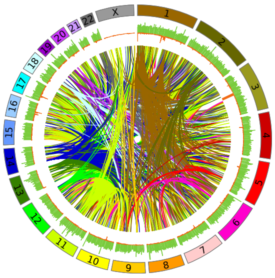

PCAF Red-C
fRIP-Seq K562

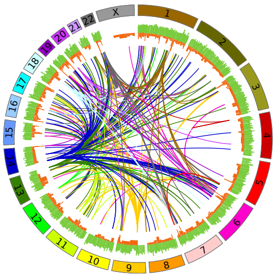

PCBP1 Red-C eCLIP-Seq K562

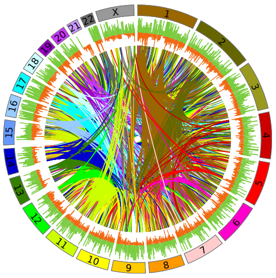

PHF8 Red-C fRIP-Seq K562

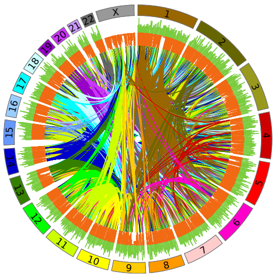

PTBP1 Red-C eCLIP-Seq K562

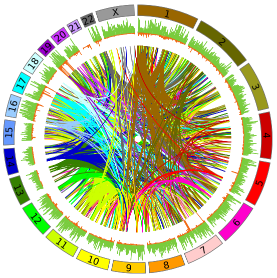

RBBP5 Red-C fRIP-Seq K562

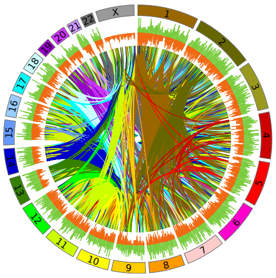

RBFOX2 Red-C eCLIP-Seq K562

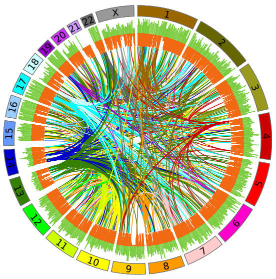

RBM15 Red-C eCLIP-Seq K562

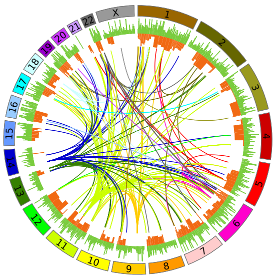

RBM22 Red-C eCLIP-Seq K562

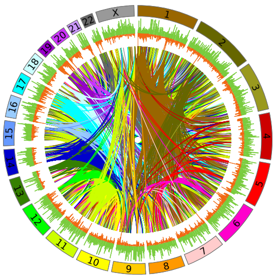

SAFB Red-C eCLIP-Seq K562

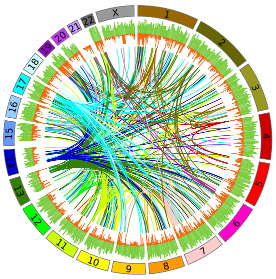

SAFB2 Red-C eCLIP-Seq K562

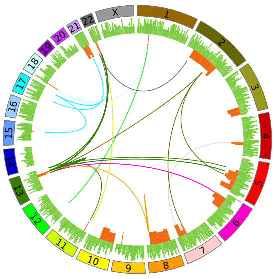

SRSF1 Red-C eCLIP-Seq K562

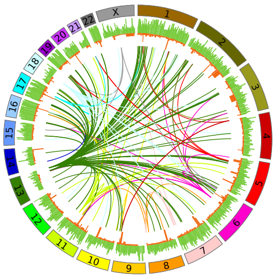

SRSF7 Red-C eCLIP-Seq K562

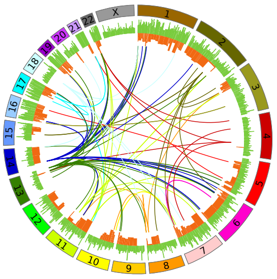

SRSF9 Red-C eCLIP-Seq K562

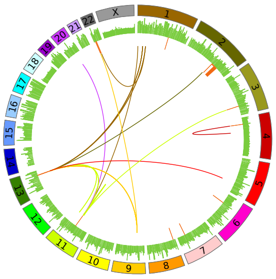

SUZ12 GRID-Seq fRIP-Seq mESC

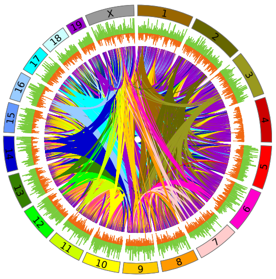

SUZ12 GRID-Seq eCLIP-Seq mESC

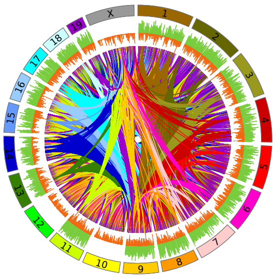

SUZ12 RADICL-Seq 2FA fRIP-Seq mESC

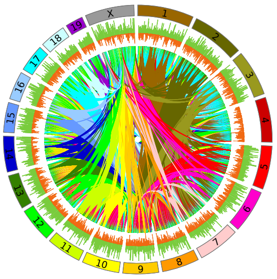

SUZ12 RADICL-Seq 2FA eCLIP-Seq mESC

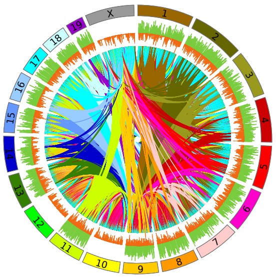

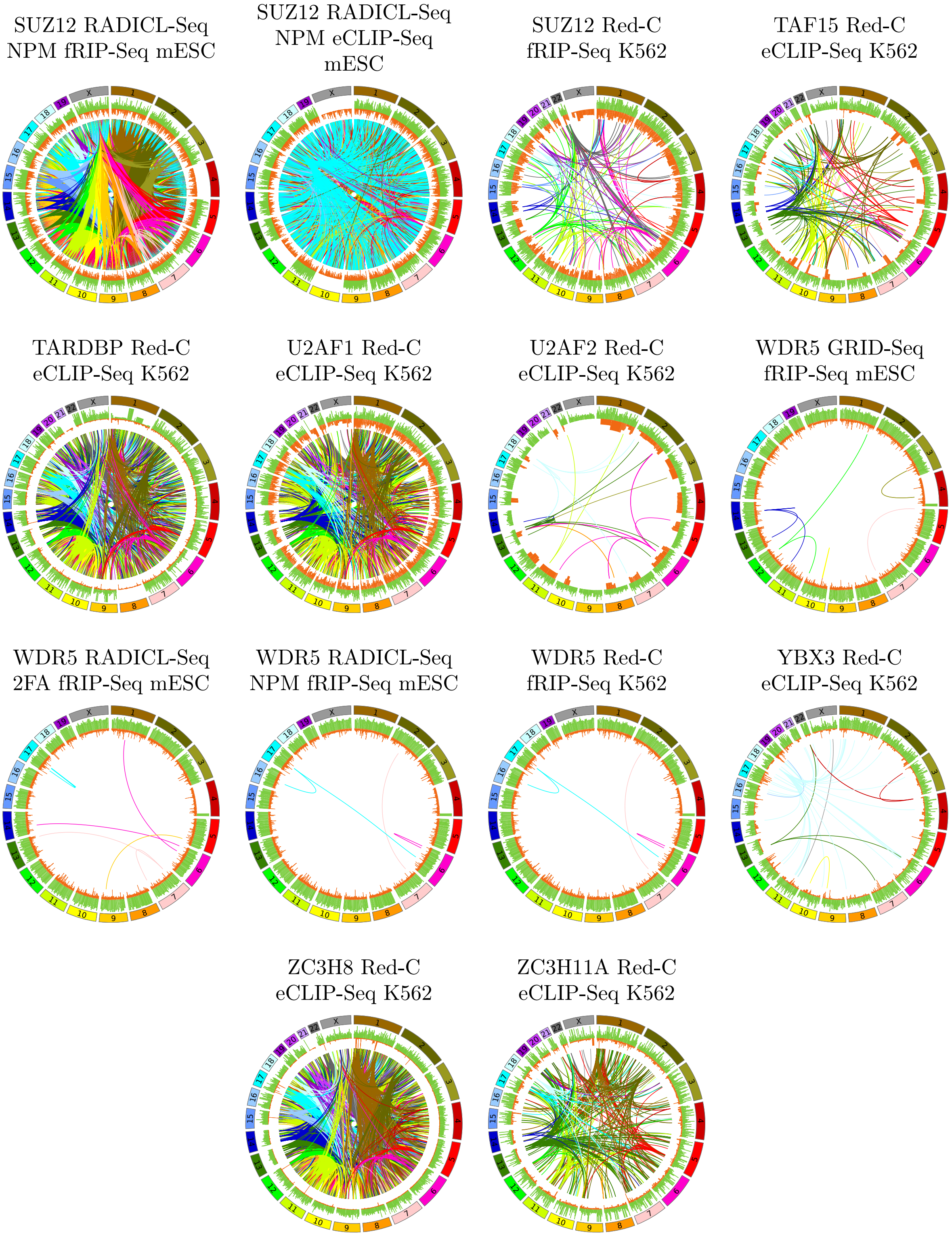


**Supplementary Figure S1.** Circos diagrams for constructed protein mediated RNA-DNA interactions, or triads. Legend for diagrams is shown in the main text.

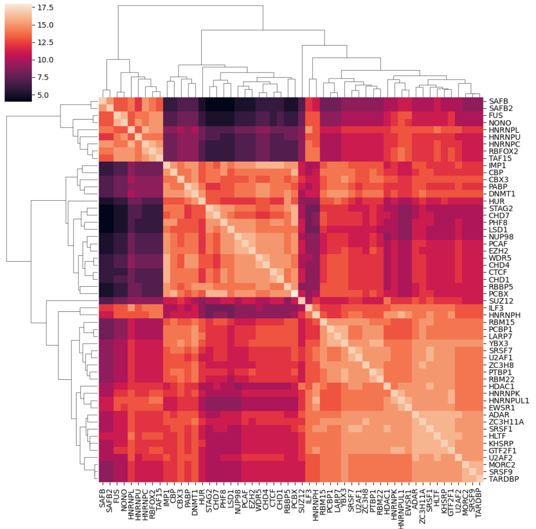


**Supplementary Figure S2.** Protein clusterization based on their "related" proteins

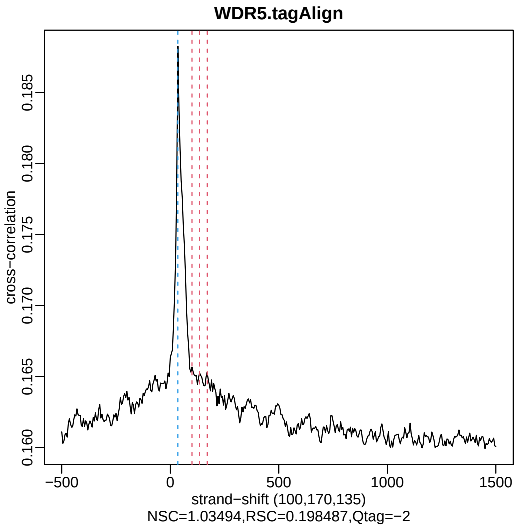


**Supplementary Figure S3.** Strand cross-correlation for WDR5 ChIP-Seq.

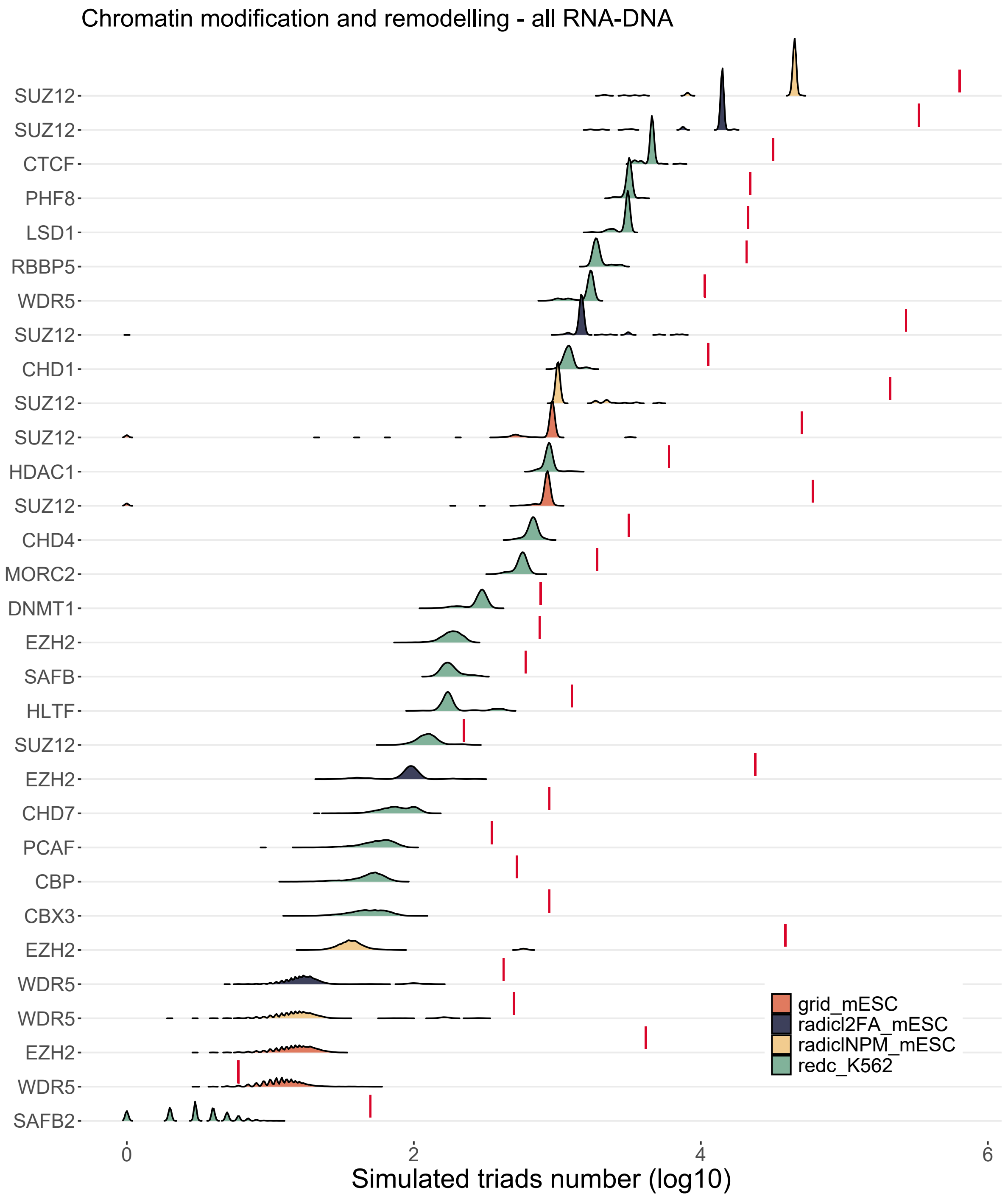


**Supplementary Figure S4.** Ridgeline plot for number of simulated triads for chromatine remodelling proteins.

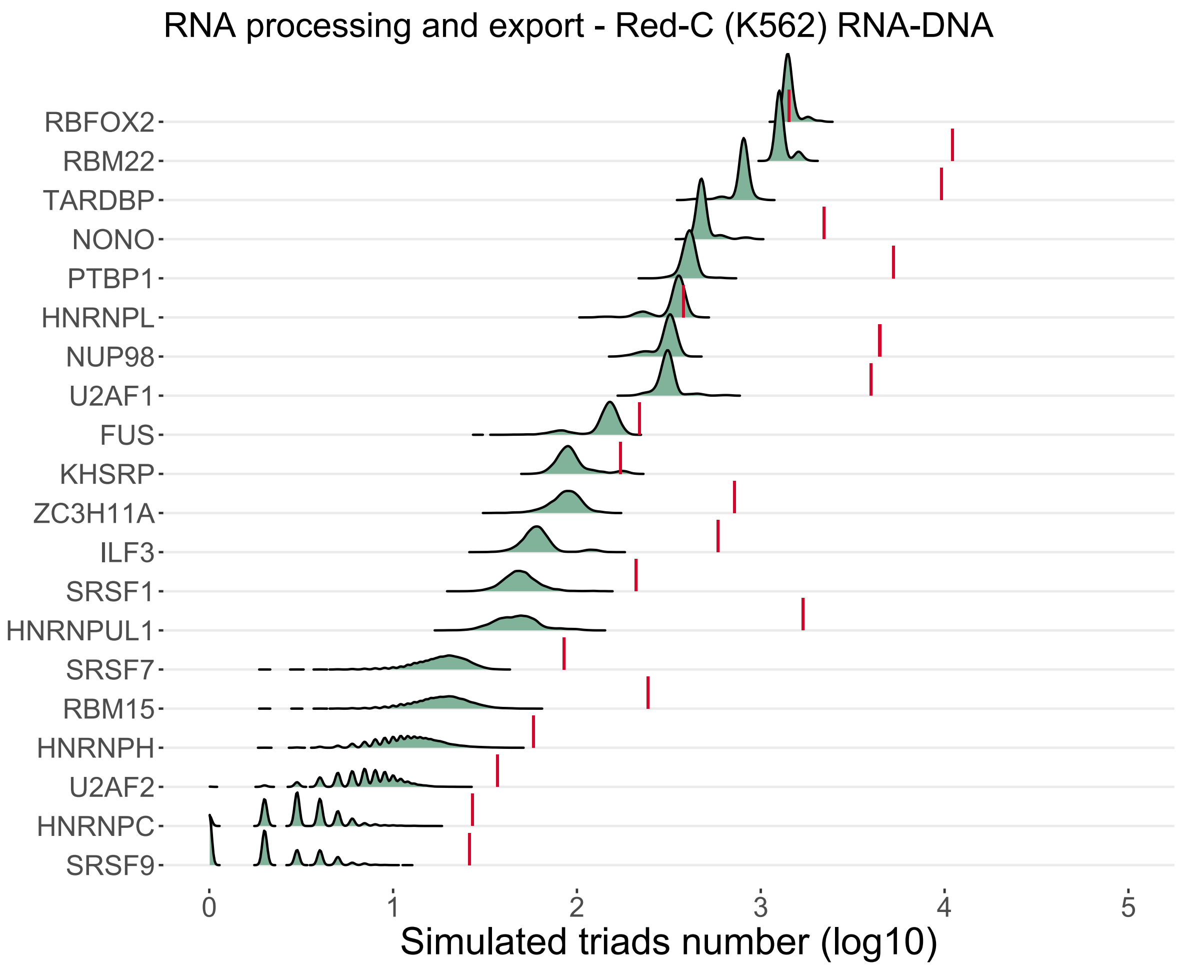


**Supplementary Figure S5.** Ridgeline plot for number of simulated triads for RNA processing and export proteins.

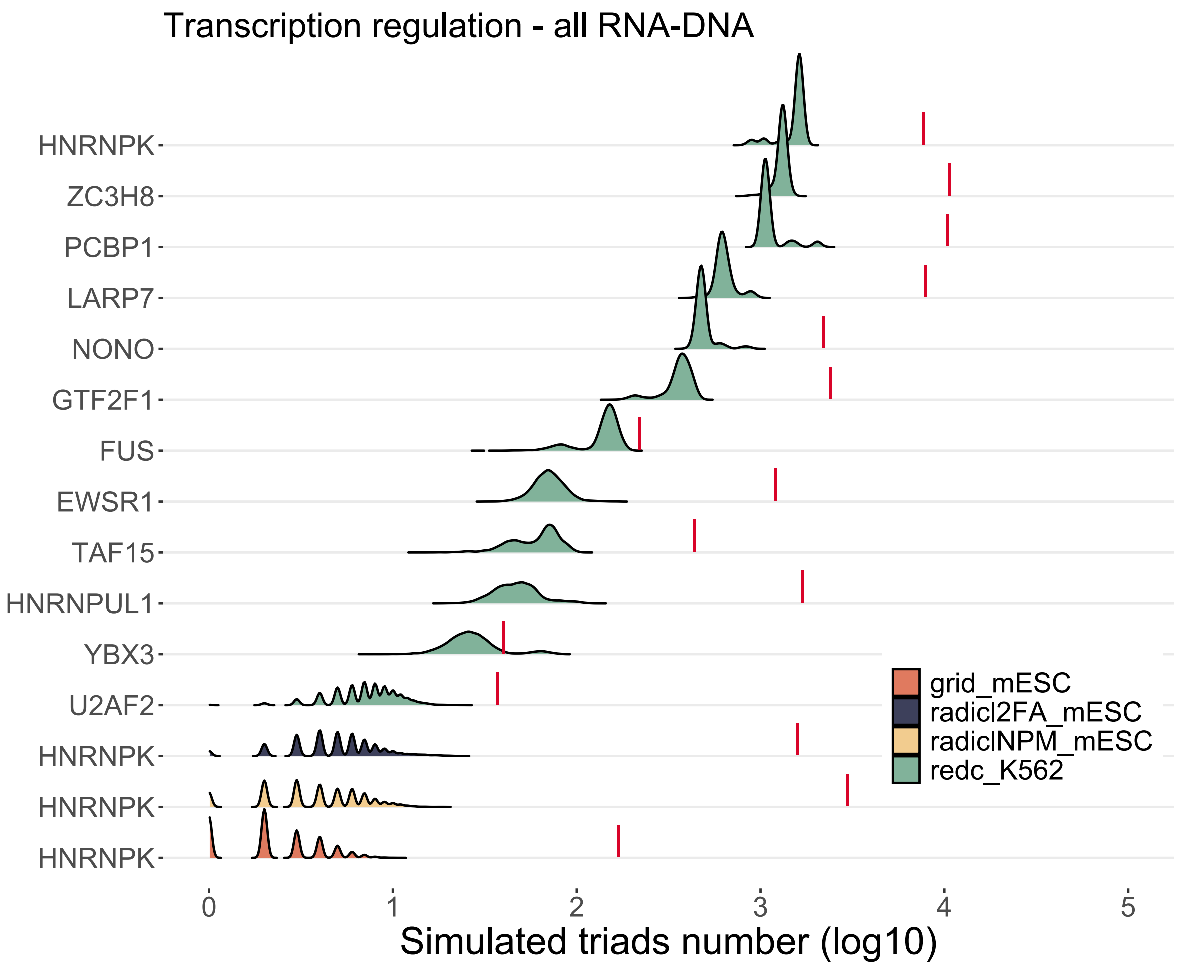


**Supplementary Figure S6.** Ridgeline plot for number of simulated triads for transcription regulating proteins.

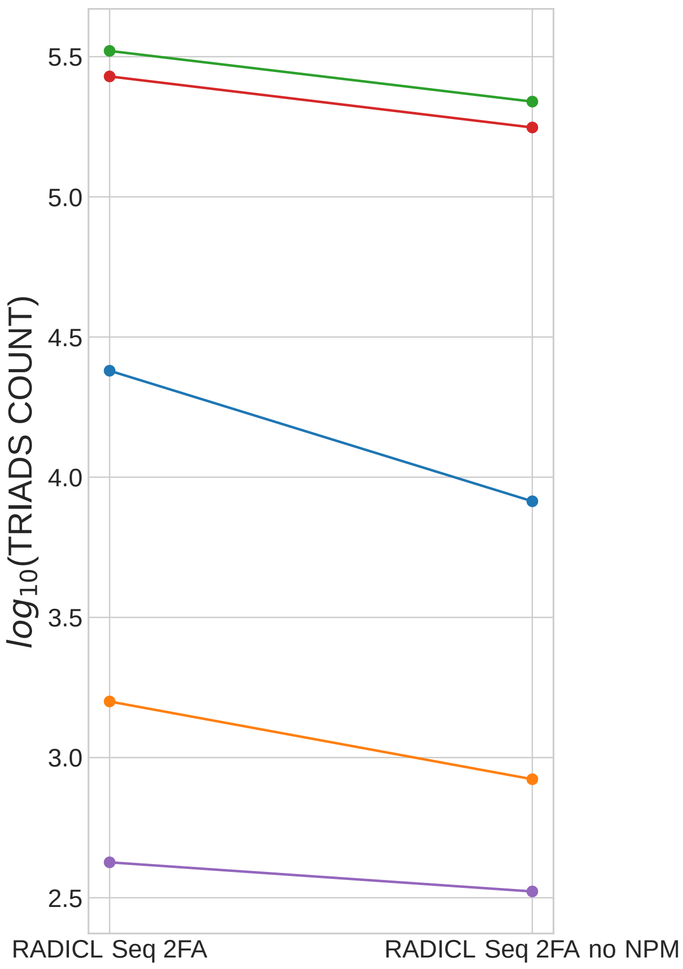


**Supplementary Figure S7.** Triads number desaturation when filtering the 2FA contacts shared with NPM phase.

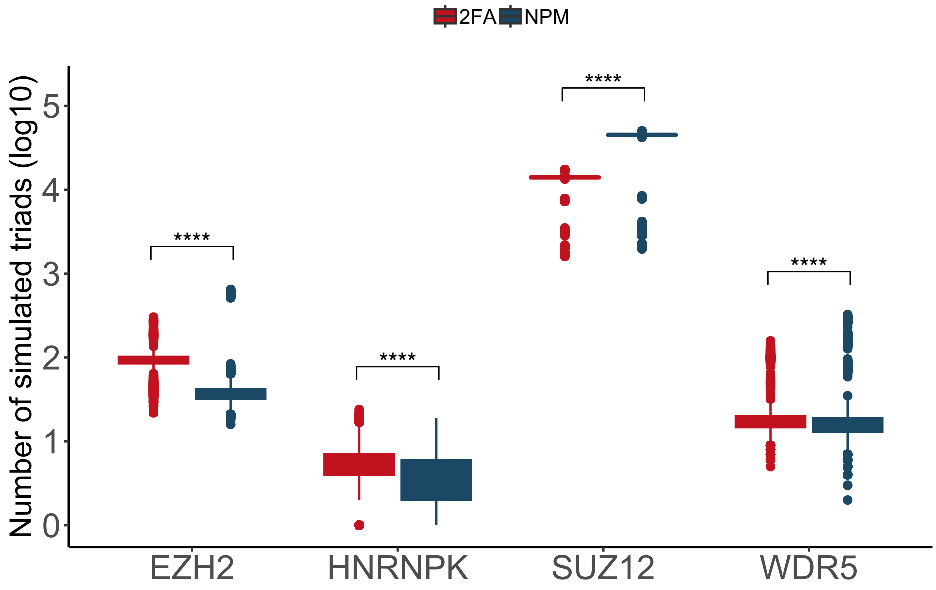


**Supplementary Figure S8.** Number of triads simulated for RADICL-Seq 2FA and RADICL-Seq NPM datasets differs significantly with the latter number being larger.

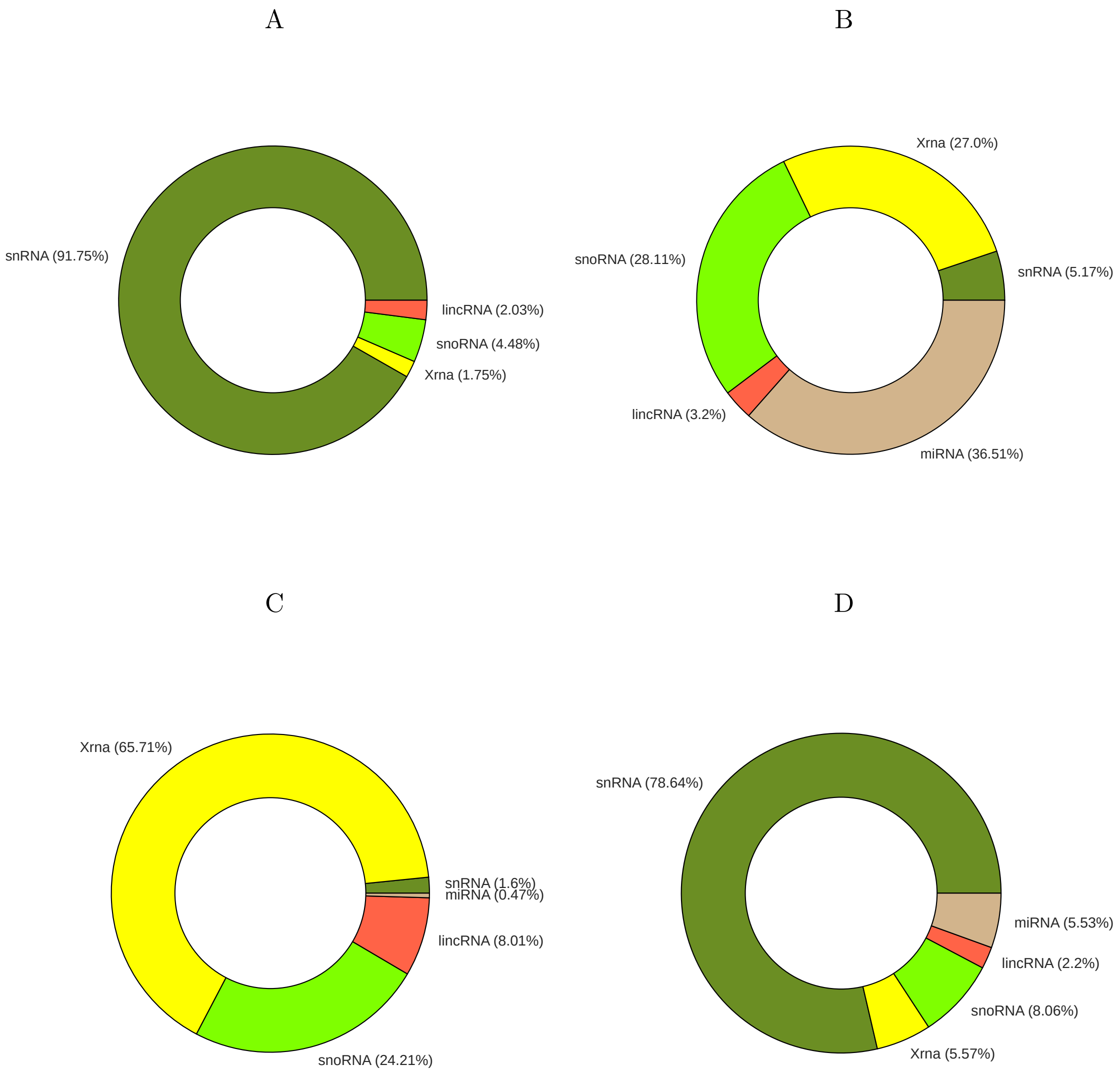


**Supplementary Figure S9.** Biotype content of RNAs forming triads in all fRIP-based triads **(A)**, all eCLIP-based triads **(B)**, triads shared by two experiments **(C)** and triads present only in one of datasets **(D)**.

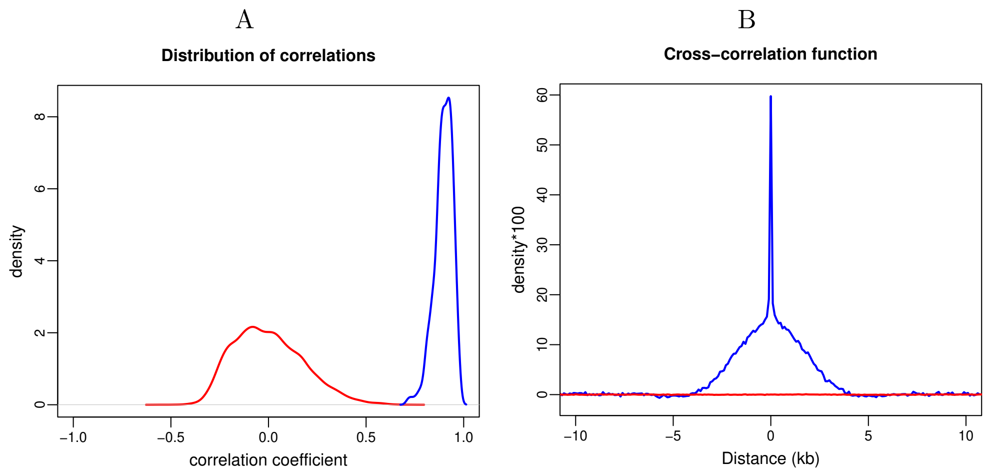


**Supplementary Figure S10.** SUZ12 DNA-parts tracks correlation. **(A)** Distribution of correlations across all bins; **(B)** Cross-correlation function decline. blue — observed correlation; red — background correlations

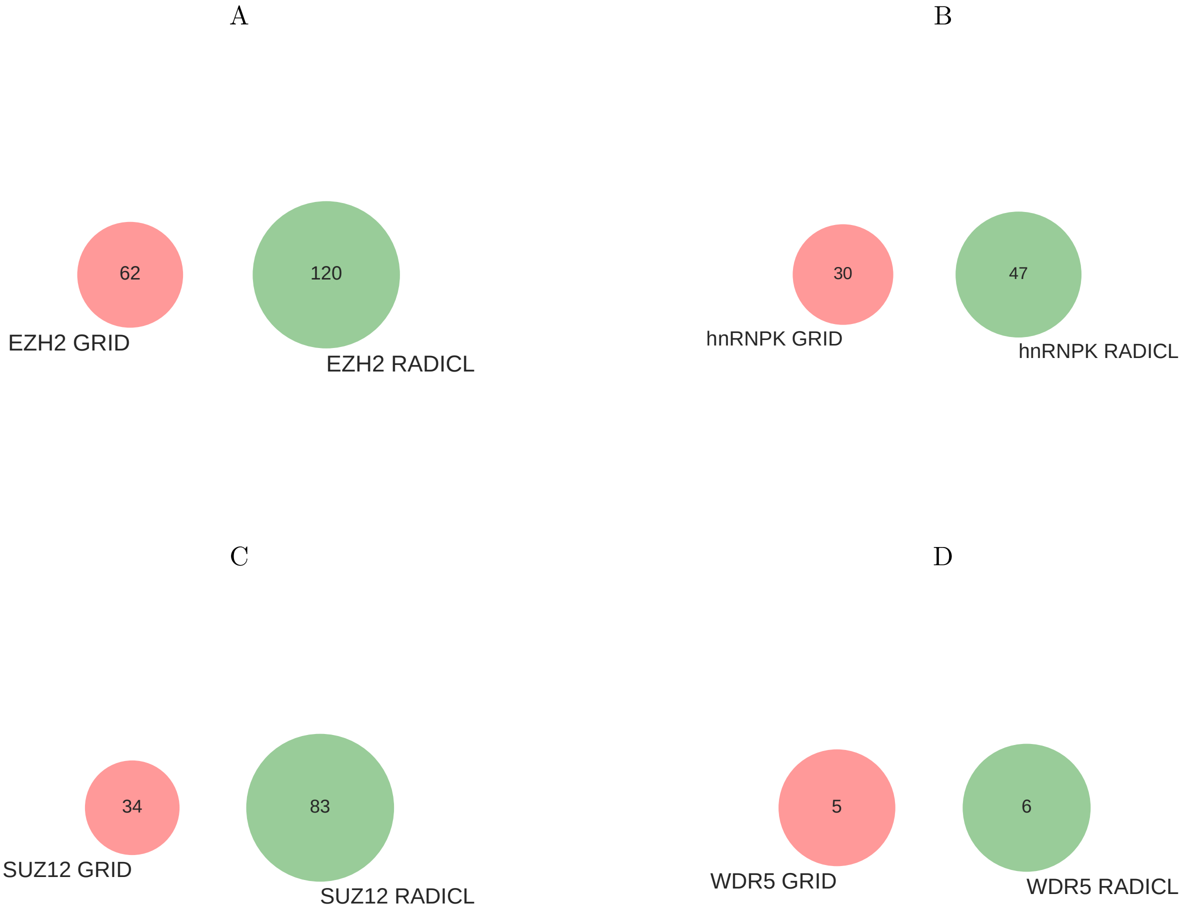


**Supplementary Figure S11.** Common RNA between contacts of RADICL-Seq and GRID-Seq in triads formed by EZH **(A)**, hnRNPK **(B)**, SUZ12 **(C)** and WDR5 **(D)** proteins.

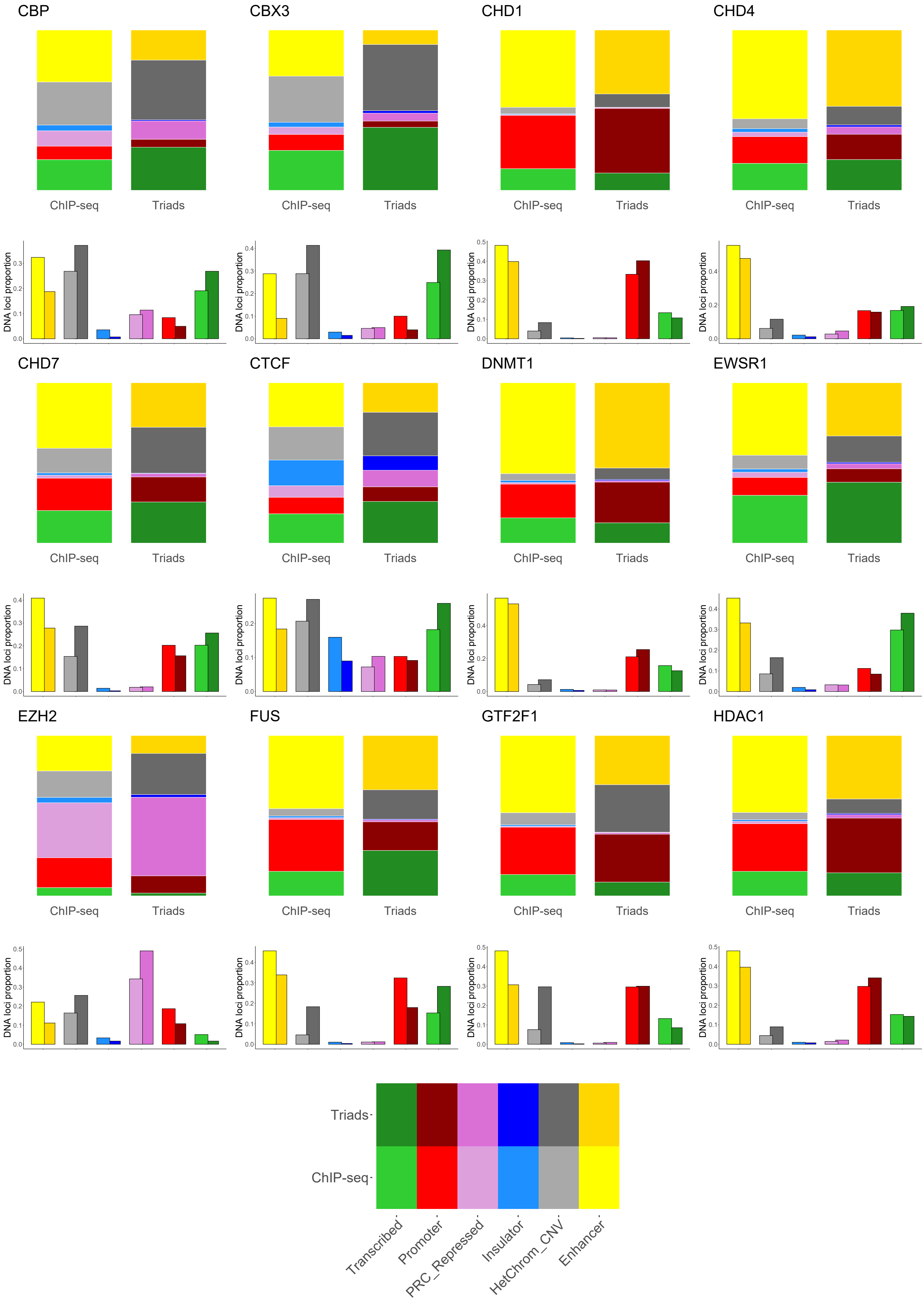
CBP
CBX3
CHD1
CHD4
CHD7
CTCF
DNMT1
EWSR1
EZH2
FUS
GTF2F1
HDAC1
ChIP-seq
Triads
DNA loci proportion
Triads
ChIP-seq
Transcribed
Promoter
PRC_Repressed
Insulator
HetChrom_CNV
Enhancer

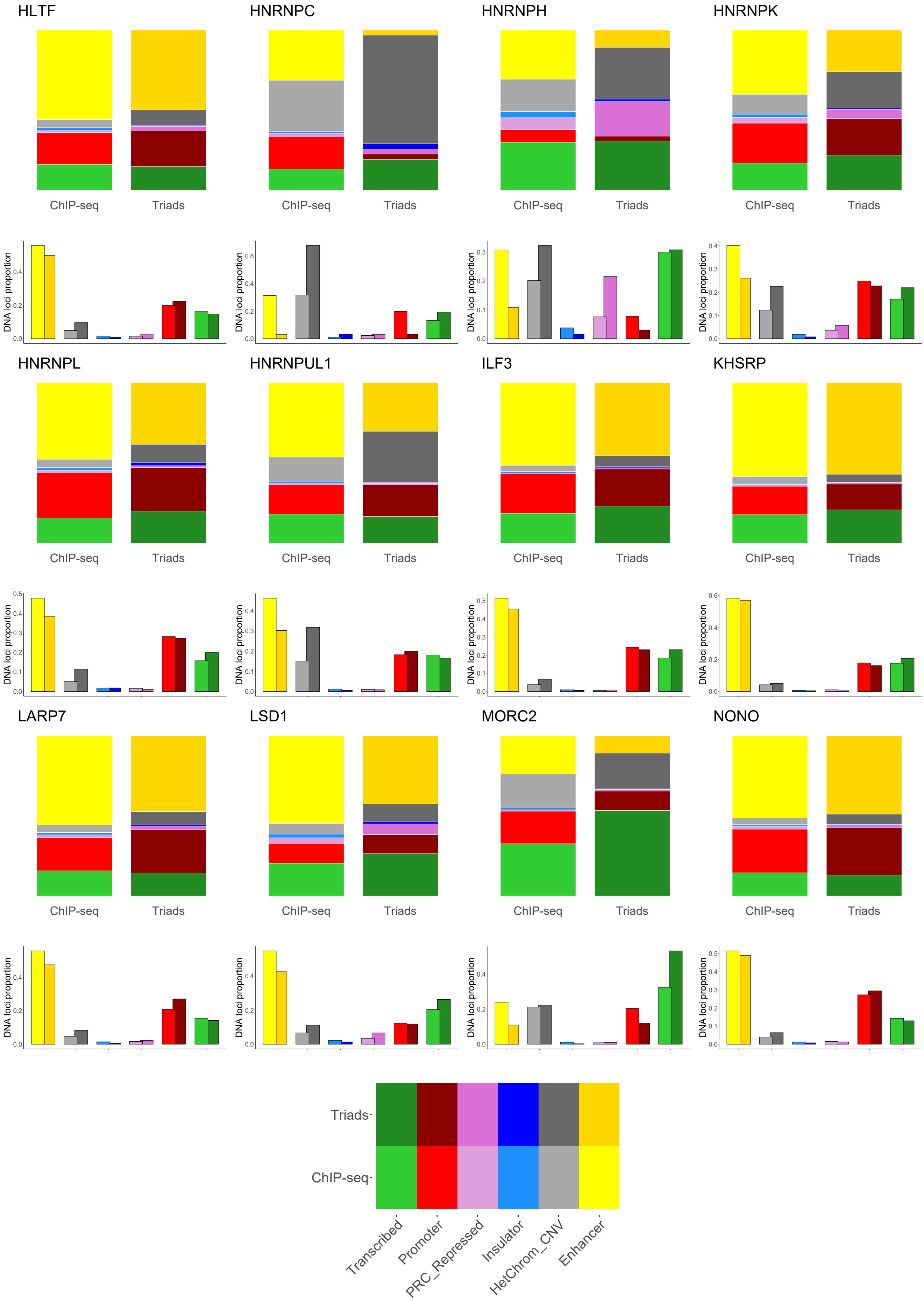
HLTF
HNRNPC
HNRNPH
HNRNPK
HNRNPL
HNRNPUL1
ILF3
KHSRP
LARP7
LSD1
MORC2
NONO
ChIP-seq
Triads
DNA loci proportion
Transcribed
Promoter
PRC_Repressed
Insulator
HetChrom_CNV
Enhancer

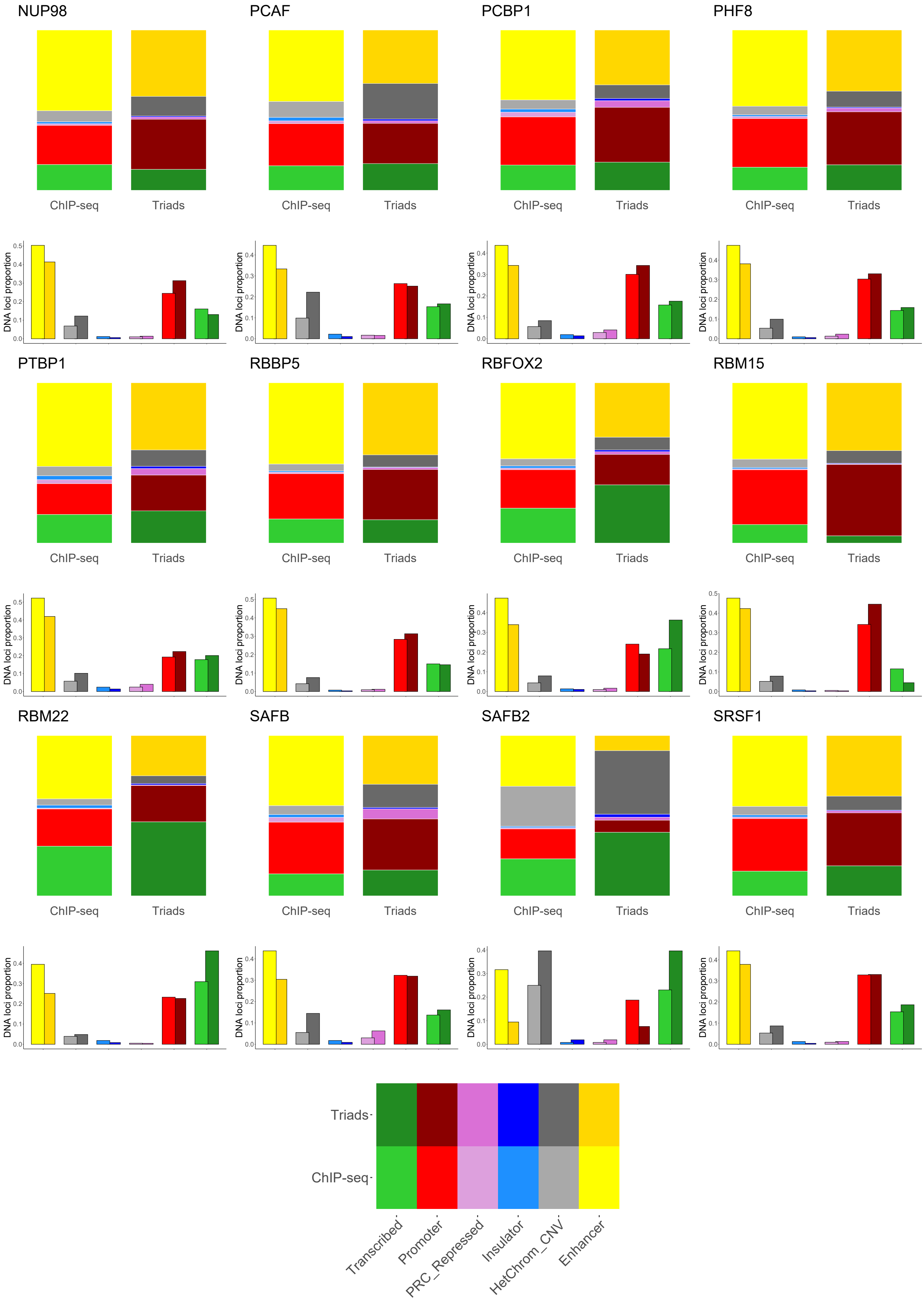
NUP98
PCAF
PCBP1
PHF8
PTBP1
RBBP5
RBFOX2
RBM15
RBM22
SAFB
SAFB2
SRSF1
ChIP-seq
Triads
DNA loci proportion
Transcribed
Promoter
PRC_Repressed
Insulator
HetChrom_CNV
Enhancer

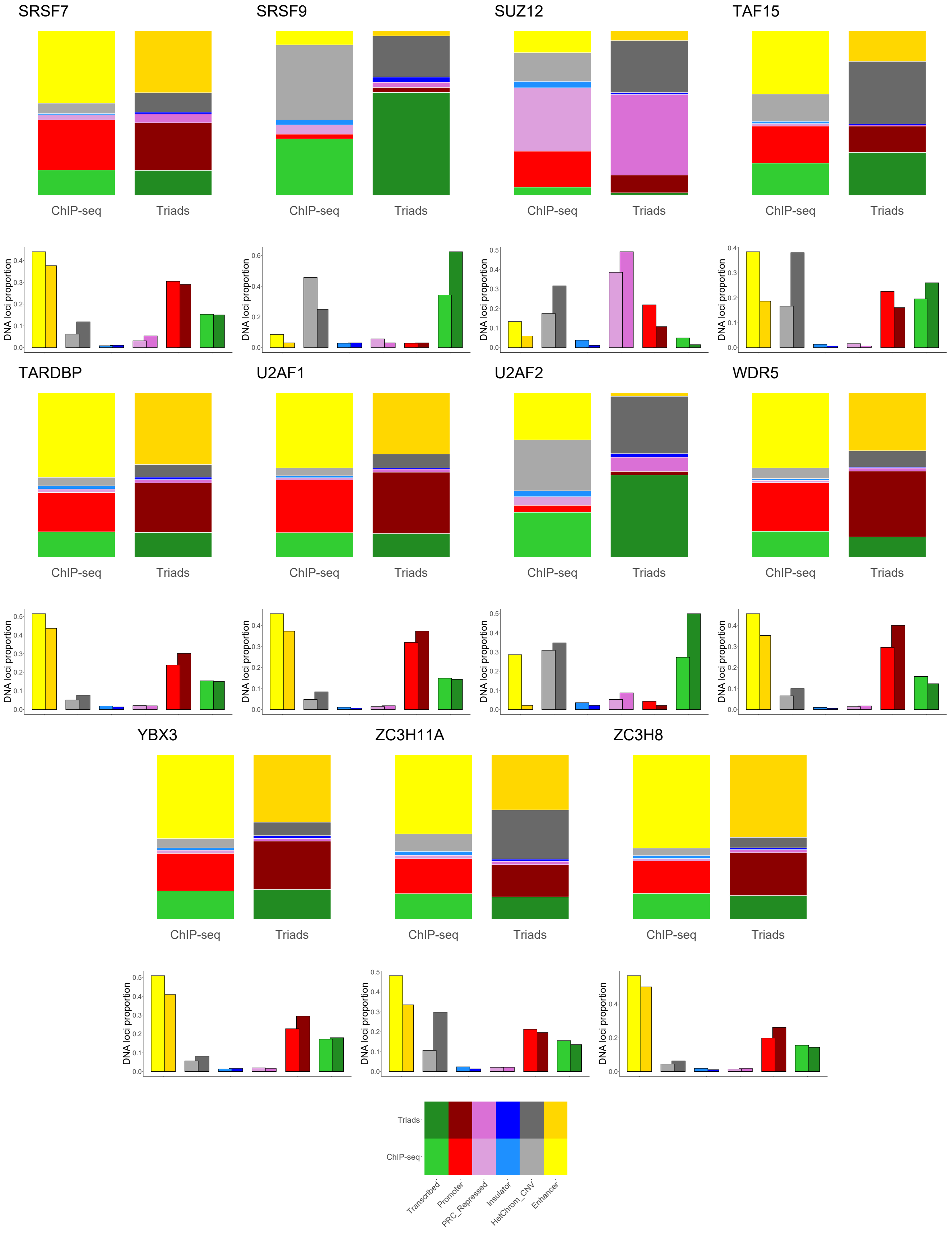


**Supplementary Figure S12.** Stacked barplots and paired barplots representing DNA parts distribution along the ChromHMM annotation states for triads of proteins in both cell lines.

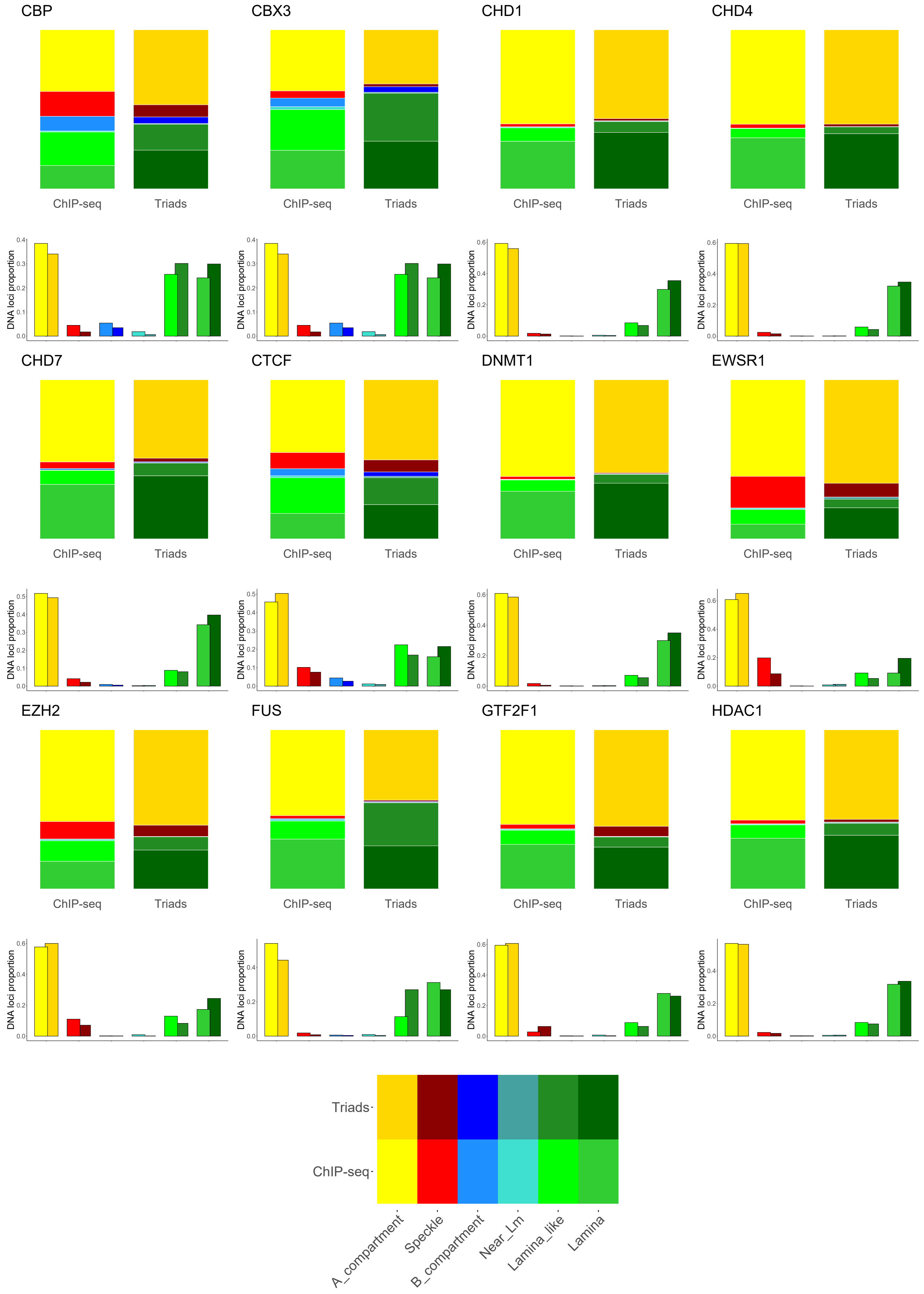
CBP
CBX3
CHD1
CHD4
CHD7
CTCF
DNMT1
EWSR1
EZH2
FUS
GTF2F1
HDAC1
ChIP-seq
Triads
DNA loci proportion
Triads
ChIP-seq
A_compartment
Speckle
B_compartment
Near_Lm
Lamina_like
Lamina

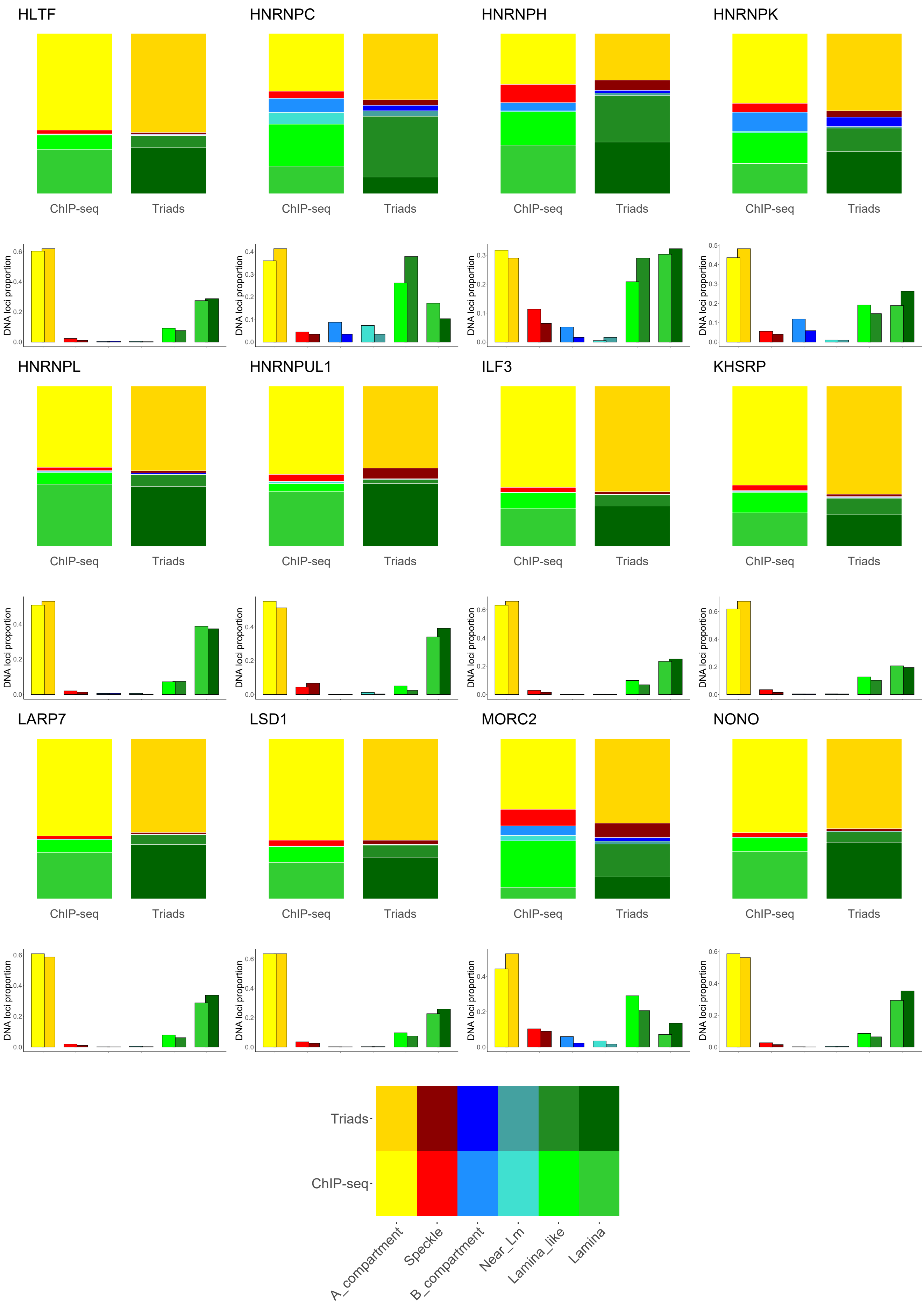
HLTF
HNRNPC
HNRNPH
HNRNPK
HNRNPL
HNRNPUL1
ILF3
KHSRP
LARP7
LSD1
MORC2
NONO
ChIP-seq
Triads
DNA loci proportion
Triads
ChIP-seq
A_compartment
Speckle
B_compartment
Near_Lm
Lamina_like
Lamina

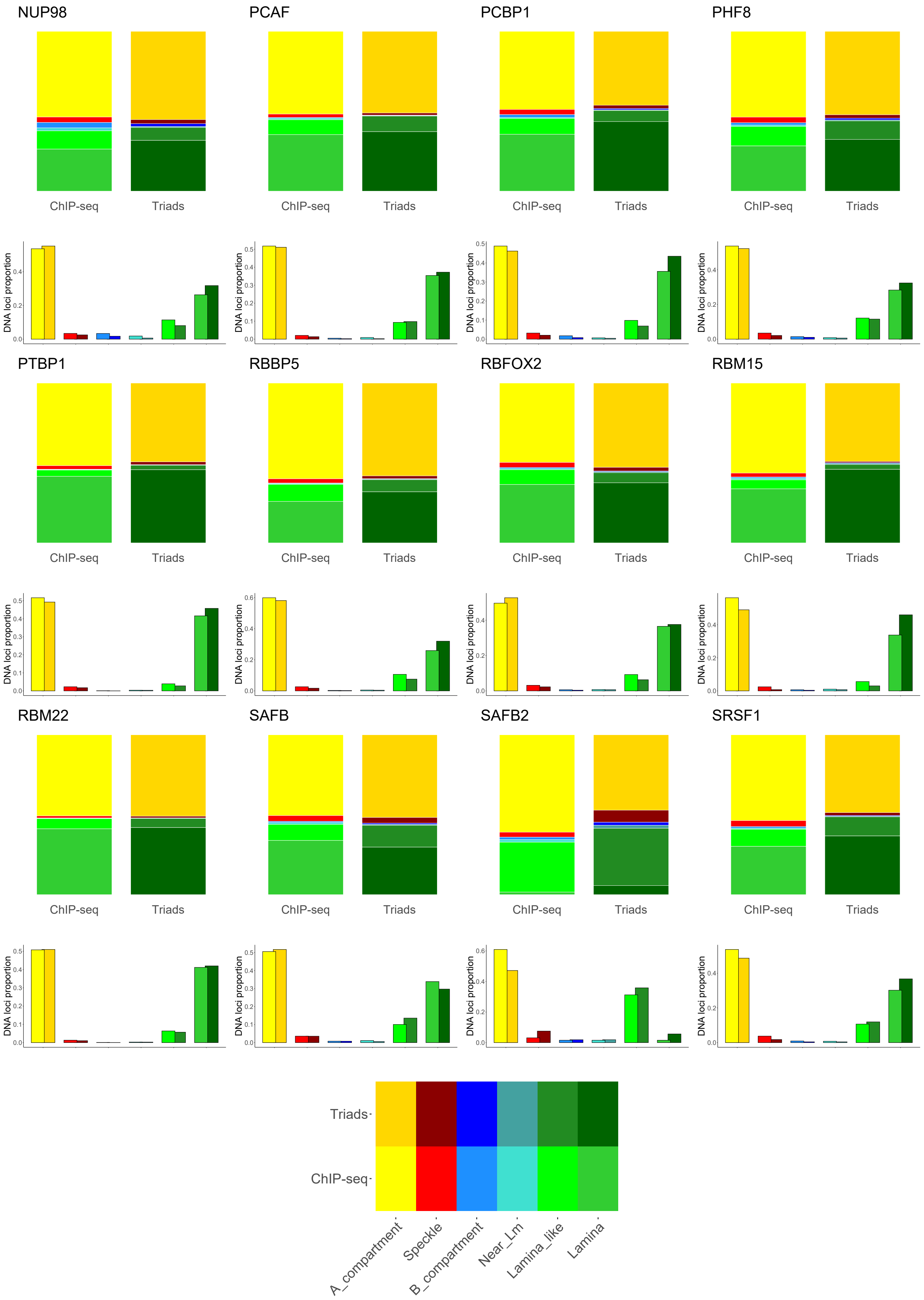
NUP98
PCAF
PCBP1
PHF8
PTBP1
RBBP5
RBFOX2
RBM15
RBM22
SAFB
SAFB2
SRSF1
ChIP-seq
Triads
DNA loci proportion
A_compartment
Speckle
B_compartment
Near_Lm
Lamina_like
Lamina

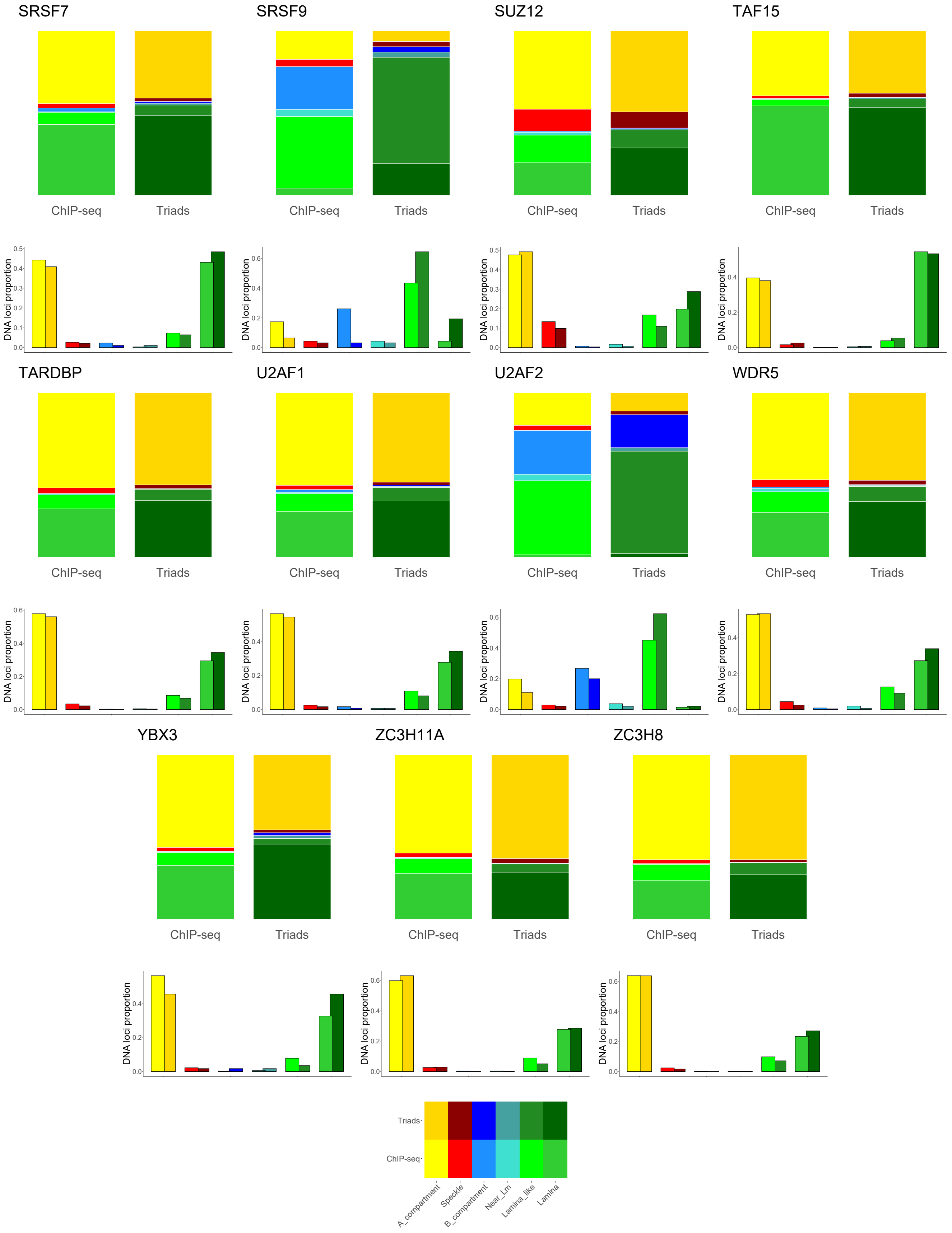


**Supplementary Figure S13.** Stacked barplots and paired barplots representing DNA parts distribution along the SPIN annotation states for triads of proteins in both cell lines.

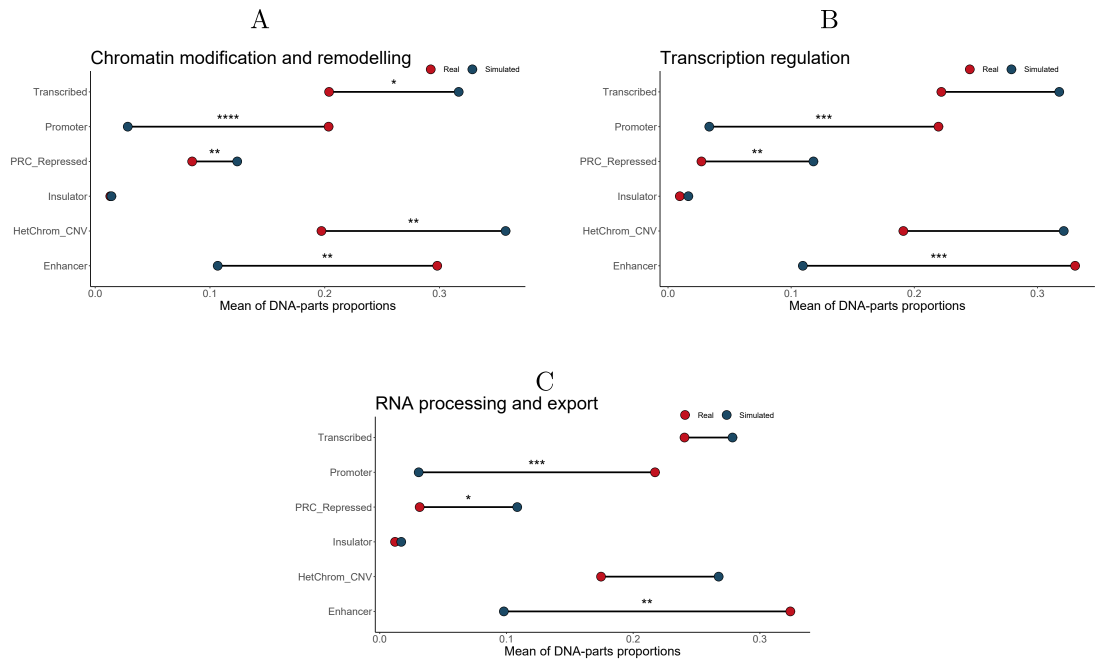


**Supplementary Figure S14.** DNA parts of triads distribution by ChromHMM states obtained using real and simulated DNA- and RNA-protein interaction data for RNA processing and exporting proteins.

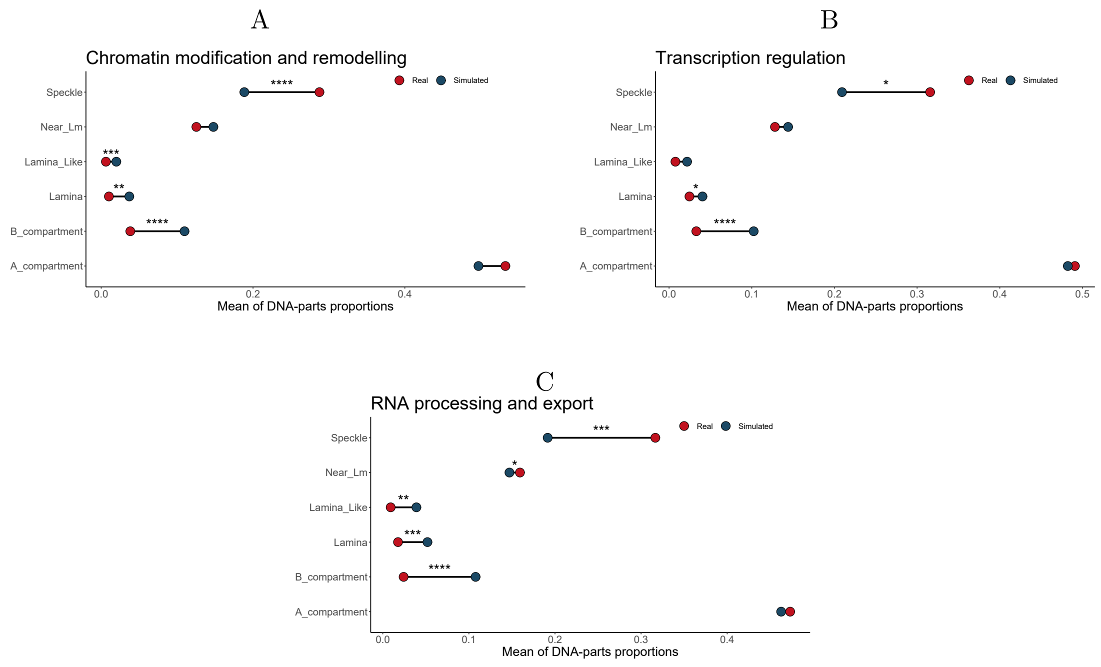


**Supplementary Figure S15.** DNA parts of triads distribution by SPIN states obtained using real and simulated DNA- and RNA-protein interaction data.

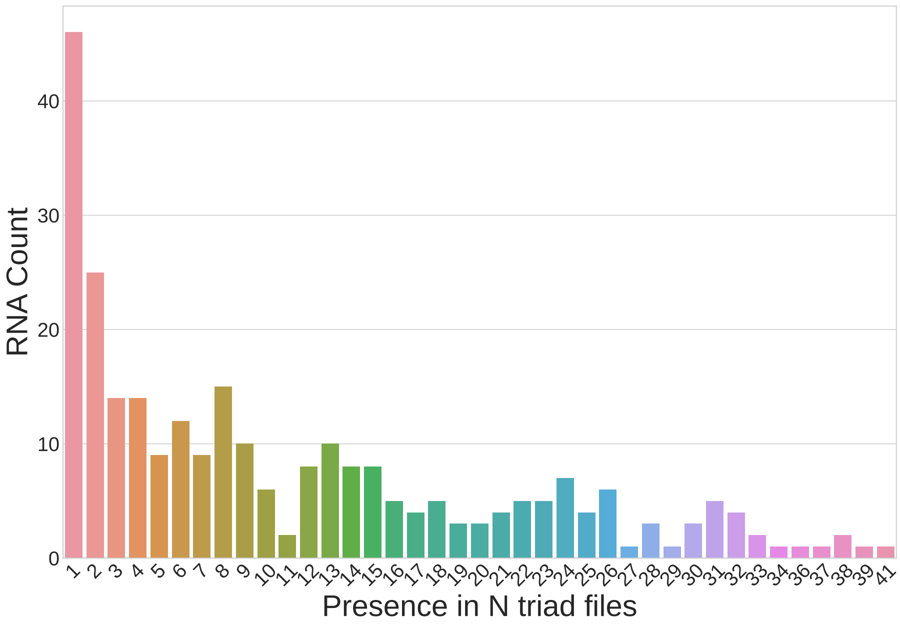


**Supplementary Figure S16.** RNA participation in triads, i.e. given an RNA that if forming triads, what is the possible range of proteins number it forms triads with?
